# A subtype-specific gene regulatory program controls differential circuit integration of deep CA3 pyramidal neurons

**DOI:** 10.64898/2026.11.03.763963

**Authors:** Danie Daaboul, Jeroen Vandensteen, Dan Dascenco, Blanca Lorente-Echeverría, Joris Vandenbempt, Elke Leysen, Malou Reverendo, Ine Vlaeminck, Lise Vervloessem, Nicolas Alcaraz, Marta Wojno, Suresh Poovathingal, Kristofer Davie, Pedro Magalhães, Niklas Kempynck, Gabriela Michel, Nelson Spruston, Keimpe Wierda, Stein Aerts, Joris de Wit

## Abstract

How closely related neuronal subtypes integrate into different circuits remains poorly understood. Deep and superficial hippocampal CA3 pyramidal neurons (PNs) differ in intrinsic firing and mossy fiber (MF) input at specialized postsynaptic structures termed thorny excrescences (TEs). Combining developmental single-nucleus multiomics with genetic targeting, morphological analysis and electrophysiology, we identify the transcription factor ST18 as a regulator of deep CA3 subtype identity. ST18 marks sparsely thorny and burst-firing neurons, and gene regulatory analyses predict that it represses a superficial-associated program involved in dendritic spine development, synaptic organization, and electrophysiological properties. Postnatal ST18 loss shifts the transcriptional and chromatin landscape of deep CA3 PNs toward a superficial-like state. These molecular changes are accompanied by increased TE elaboration approaching superficial levels, enhanced excitatory synaptic activity, and reduced burst-firing. Together, these findings identify ST18-dependent restriction of an alternative subtype program as a mechanism linking molecular identity to distinct input connectivity patterns and intrinsic firing properties during postnatal circuit maturation.

## Introduction

How closely related neuronal subtypes develop distinct connectivity patterns is a central question in circuit development. Pyramidal neuron (PN) subtypes in the cortex and hippocampus can occupy the same anatomical region yet become embedded in distinct circuits. Despite growing knowledge about their molecular, cellular and functional diversity, the gene regulatory programs that shape PN subtype-specific circuit integration are largely unknown.

In the canonical hippocampal trisynaptic circuit, CA3 PNs occupy a central position, receiving mossy fiber (MF) input from dentate gyrus granule cells and relaying information to CA1 PNs through Schaffer collateral projections^1^. MF inputs contact thorny excrescences (TEs), unusually elaborate, branched dendritic spine complexes on CA3 dendrites that form highly specialized postsynaptic interfaces with large MF boutons^2–6^. However, recent studies show that this canonical description only partially captures the wiring complexity of the CA3 circuit. CA3 PNs exhibit substantial heterogeneity across different spatial axes, including the radial axis that separates deep from superficial cells^7^. Differences in MF innervation are a prominent feature of this radial organization. Superficial CA3 is enriched in regular-spiking PNs with prominent TEs. In contrast, deep CA3 is enriched in burst-firing PNs with sparse or absent TEs, termed sparsely thorny or athorny cells, respectively^8–11^. Athorny CA3 PNs were shown to lack functional MF input^10^. In addition, recurrent excitation and inhibitory innervation show a subtype-specific organization across this axis in the CA3 circuit^11,12^. Together, these findings suggest that CA3 PN diversity shapes how dentate gyrus input is distributed into and processed within the recurrent CA3 network, raising the question of how these distinct circuit roles emerge during development. Although differences in gene expression patterns between CA3 PNs have been identified along the septal-temporal and proximal-distal axes^13^, the gene regulatory programs that shape deep and superficial CA3 PN identities and their associated synaptic input remain unresolved.

In cortical PN subtypes, transcription factors (TFs) have been shown to regulate axonal projection patterns through downstream effector genes encoding axon guidance receptors^14–16^. FEZF2 in deep cortical layer 5 PNs regulates expression of the guidance receptor EphB1 to control corticospinal axon targeting^14^. SATB2 in upper layer 5 cortical PNs controls callosal axon targeting via its downstream effectors EphA4 and Unc5H3^16^. These studies provide a framework for linking subtype identity to patterns of output connectivity. In contrast, much less is known about transcriptional regulation of subtype-specific input connectivity. In *C. elegans*, the transcriptional repressor UNC-4 specifies VA motor neuron inputs by suppressing a transcriptional program associated with VB neurons^17–19^. In mouse, ectopic FEZF2 expression alters excitatory inputs onto L4 neurons and inhibitory inputs onto L2/3 neurons^20,21^. These studies demonstrate that TFs can shape input connectivity. However, the downstream effectors mediating these changes remain poorly defined. How TF-effector relationships are organized into developmental gene regulatory programs linking subtype identity to input connectivity patterns remains poorly understood.

Here, we combined single-nucleus multiomic profiling, genetic access to a defined neuronal subtype, and functional perturbation to investigate how gene regulatory programs shape CA3 PN subtype diversification during postnatal development. We generated a developmental atlas of hippocampal transcriptional and chromatin dynamics, providing a resource for resolving the regulatory programs underlying neuronal maturation and subtype identity. Within CA3, we identified the TF ST18 as a defining marker of deep PNs and generated an *St18*-Cre driver line to access this population. *St18*-expressing CA3 PNs exhibited sparse or absent TEs and burst-firing properties. Gene regulatory analyses predicted ST18 as a candidate repressor of a superficial PN program associated with dendritic spine development, synaptic organization, and electrophysiological properties. Postnatal loss of ST18 shifted deep CA3 PNs toward a superficial-like transcriptional and chromatin state, accompanied by an increase in TE density to levels observed in superficial PNs, enhanced excitatory synaptic activity, and a decrease in burst-firing. Together, these findings establish a framework for linking developmental gene regulatory programs to neuronal diversity and identify ST18 as a regulator coordinating CA3 subtype identity, synaptic input, and intrinsic physiology during postnatal circuit development.

## Results

### snMO analysis reveals transcriptional dynamics and cell-type-specific regulatory programs during postnatal hippocampal development

To investigate the gene regulatory dynamics underlying CA3 PN subtype diversification, we expanded our snMO atlas of the developing mouse hippocampus^22^ spanning key stages of postnatal maturation: axonogenesis (postnatal day P5); establishment of initial synaptic contacts (P10); peak synaptogenesis (P15), and synaptic refinement and stabilization (P28) (Fig. 1a). We added replicates at each of the P15 and P28 time points, ensuring representation of both sexes at these ages. After quality control (Extended Data Fig. 1a-c), we integrated gene expression data from all samples and annotated these by label transfer from the Allen Brain Cell (ABC) Atlas (). Nuclei primarily segregated according to cellular identity rather than developmental stage (Fig. 1b, c). We identified 12 major cell classes, with each cell class showing defined cell-type-specific marker expression patterns (Fig. 1c, Extended Fig. 1d). Cell type annotation could be further refined at the subclass level, revealing 44 major hippocampal subclasses, with several populations showing expected age-dependent changes in abundance (Fig. 1d).

**Figure 1.**
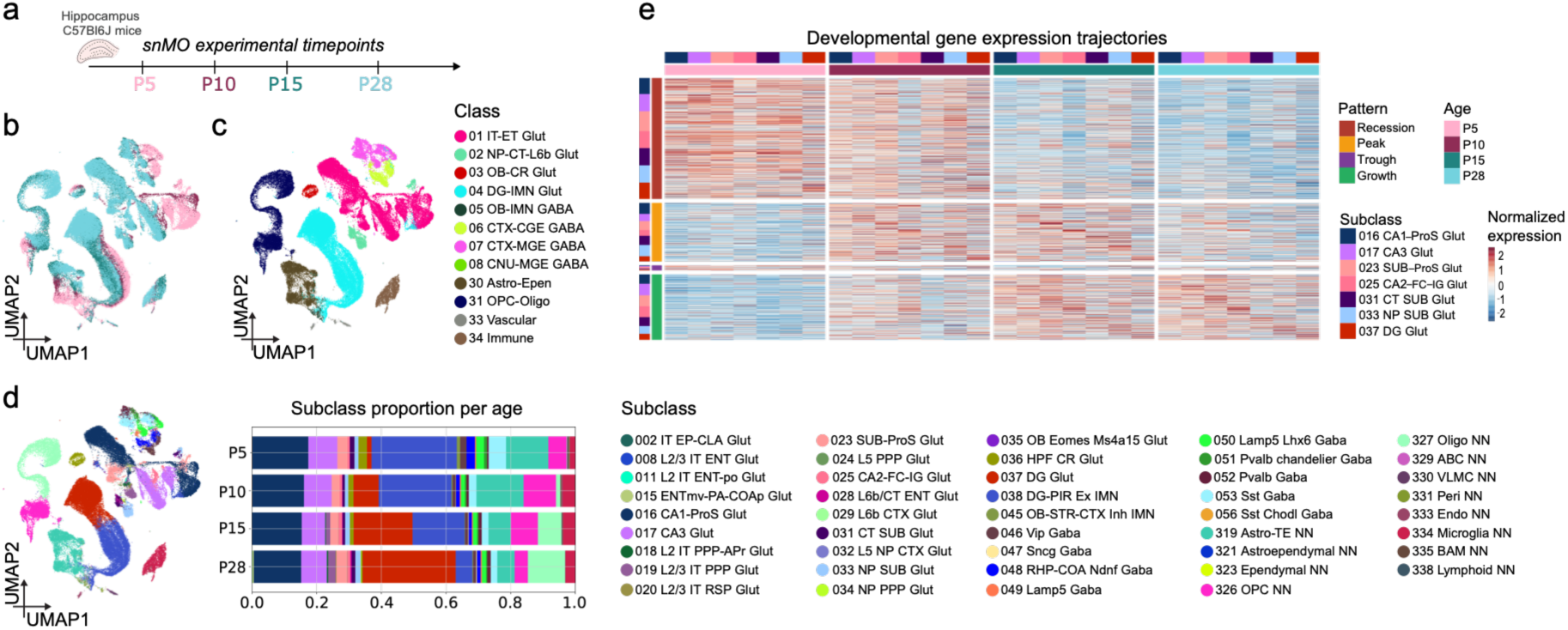
Single-nucleus multiome atlas of the postnatal mouse hippocampus. **(a)** Schematic of the postnatal developmental stages sampled for hippocampal snMO profiling (postnatal day (P)5, P10, P15 and P28). **(b)** UMAP of integrated snMO profiles colored by developmental age. **(c)** UMAP colored by major cell class following label transfer from the Allen Brain Cell (ABC) Atlas reference. **(d)** Left, UMAP colored by subclass annotation. Right, relative proportions of hippocampal subclasses across developmental timepoints. **(e)** Heatmap of temporally dynamic genes across hippocampal glutamatergic subclasses and developmental timepoints, grouped into four expression patterns: recession, peak, trough and growth. Columns are annotated by developmental age and subclass, and colors indicate normalized gene expression.

Temporal transcriptional dynamics analysis across hippocampal glutamatergic subclasses segregated dynamic genes within individual subclasses into four predominant developmental trajectories: early declining (“recession”), transiently activated (“peak”), progressively increasing (“growth”), and intermediate downregulated (“trough”) expression programs (Fig. 1e). Within the CA3 glutamatergic subclass, enrichment analysis revealed a temporal progression in which CA3 neurons first downregulate axon guidance programs, transiently activate synapse-assembly genes, and then progressively induce molecular pathways supporting stable synaptic transmission, signaling, and mature hippocampal circuit function (Extended Fig. 2a). The transcriptional dynamics of effector genes were paralleled by changes in expression of transcriptional regulators (Extended Fig. 2b) and structured patterns of chromatin accessibility across subclasses and developmental stages (Extended Fig. 2c, d).

A sequence-to-function model (CREsted)^23^ trained to predict chromatin accessibility directly from genomic sequence identified combinations of regulatory sequence features associated with specific hippocampal cell types and developmental stages (Extended Data Fig. 3a-c), while regulatory network inference identified corresponding cell-type-specific gene regulatory modules, including an early program and a distinct DG/CA3/CA2-associated module (Extended Data Fig. 3d). Together, these analyses provide a multiomic atlas of the postnatal mouse hippocampus, reveal coordinated transcriptional and regulatory programs underlying cell-type-specific maturation, and establish a framework for dissecting CA3 PN diversification.

### Molecular and spatial organization of CA3 subtypes

We next asked whether molecularly distinct CA3 glutamatergic subpopulations could be resolved in the postnatal hippocampal snMO dataset and whether these populations corresponded to known anatomical axes of CA3 organization. Leiden clustering resolved CA3-annotated cells into four main subpopulations with comparable RNA quality metrics (Fig. 2a, Extended Data Fig. 4a). Annotation at the supertype level (ABC Atlas)^24^ revealed multiple supertypes across these CA3 clusters, including CA3-Glut1, CA3-Glut2, CA3-Glut3, CA3-Glut4, and CA3-Glut5 supertypes (Fig. 2b). All five supertypes were already represented at P5 and persisted across postnatal development through P28 (Fig. 2c).

**Figure 2.**
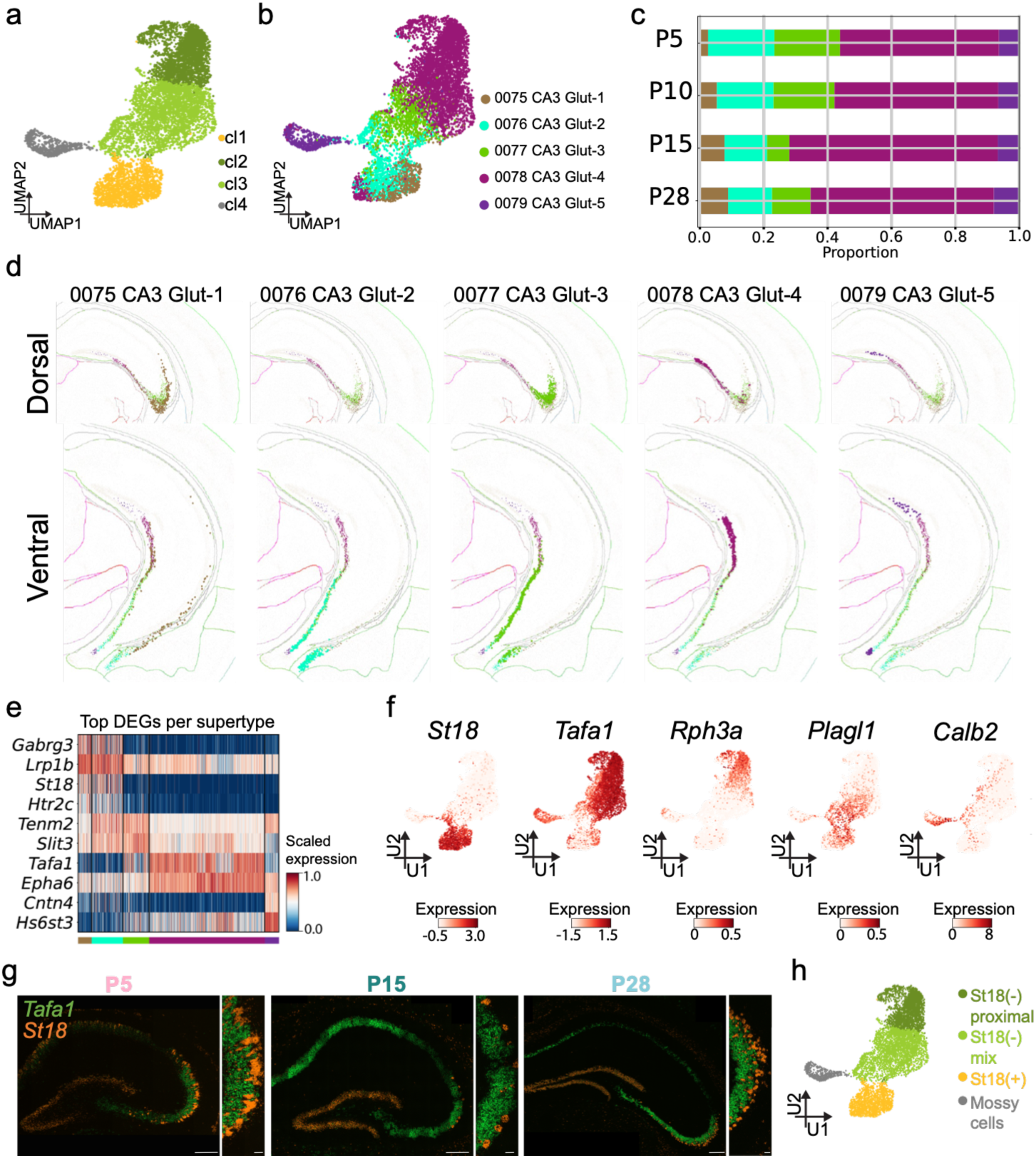
Molecular and spatial heterogeneity of CA3 pyramidal neurons across postnatal development. (a, **b)** UMAP of CA3 neurons colored by unsupervised clusters (a) and Allen Brain Cell (ABC) Atlas supertype annotations (b). **(c)** Relative proportions of CA3 supertypes across developmental timepoints. **(d)** Spatial distribution of CA3 supertypes in dorsal (top) and ventral (bottom) hippocampal sections from the ABC Atlas MERFISH dataset. **(e)** Heatmap of top differentially expressed genes between CA3 supertypes (Wilcoxon rank-sum test). **(f)** UMAPs showing expression of selected CA3 marker genes identified by pairwise comparisons between CA3 supertypes. **(g)** smFISH images showing complementary expression of *St18* (orange) and *Tafa1* (green) in CA3 at P5, P15 and P28, with higher-magnification views shown alongside. **(h)** UMAP of CA3 neurons annotated into *St18*-positive, *St18*-negative proximal, *St18*-negative mix and mossy cell populations. Scale bars in (g), 200 μm (overview) and 25 μm (magnified views).

To relate these molecular identities to anatomical organization, we examined the spatial distribution of the corresponding ABC Atlas MERFISH supertypes^24^. CA3 supertypes showed clear spatial organization in the dorsal and ventral hippocampus (Fig. 2d). CA3-Glut1 was concentrated in distal CA3 and occupied a deeper position along the radial axis of the CA3 pyramidal layer. CA3-Glut2 was largely absent from dorsal CA3 and was instead predominantly distributed in the ventral hippocampus. CA3-Glut3 was enriched in distal CA3 (CA3a-b) and occupied a more superficial position. CA3-Glut4 localized predominantly to proximal CA3 (CA3c). These spatial relationships were also apparent in ventral CA3, where CA3-Glut1, CA3-Glut3, and CA3-Glut4 remained differentially positioned along the deep-superficial and proximodistal axes. CA3-Glut5 was distinct from the other supertypes, localizing primarily to the hilus in both dorsal and ventral hippocampus, suggesting a mossy cell identity rather than CA3 principal neurons (Fig. 2d).

We next identified molecular markers capturing this spatial organization in our postnatal dataset. Top differentially expressed genes per CA3 supertype included genes involved in transcriptional regulation, signaling, adhesion, neurotransmission, and axon guidance^25–29^, consistent with differences in neuronal identity and connectivity (Fig. 2e). Expression patterns of several markers extended across individual supertypes, defining broader CA3 domains (Fig. 2e). *St18* was shared across cells annotated as CA3-Glut1 and some CA3-Glut2 cells, suggesting a deep CA3 program, whereas *Tafa1* spanned distal and proximal supertypes (CA3-Glut3 and CA3-Glut4 cells), indicative of a broader superficial pattern (Fig. 2e, f). Pairwise differential expression further resolved markers with more restricted expression, including *Rph3a* in proximal CA3 and *Plagl1* in distal and ventral populations (Fig. 2f, Extended Data Fig. 4b). *Calb2* was selectively expressed in CA3-Glut5 cells (Fig. 2f), supporting their mossy cell identity^30^.

*St18* and *Tafa1* showed highly complementary expression patterns that suggested a segregation along the deep-superficial axis (Fig. 2f). *St18* encodes a MYT-family zinc-finger transcription factor previously described as a transcriptional repressor^25^, whereas *Tafa1* encodes a secreted protein belonging to the TAFA family^31^. To test whether they indeed define spatially segregated domains across postnatal development, we performed single-molecule fluorescent in situ hybridization (smFISH). *St18* expression was enriched in the deep CA3 pyramidal layer, adjacent to stratum oriens (SO), whereas *Tafa1* was expressed in superficial CA3 (Fig. 2g). This spatially organized segregation was observed at P5, P15 and P28. *St18* and *Tafa1* were also expressed in neighboring hippocampal cell types: *St18* expression was more notable in the granule cell layer, whereas *Tafa1* was enriched in CA1 (Fig. 2g). At P5, *St18* also displayed a radial bias within CA1, with stronger expression in deep than in superficial PNs. Neither *St18*- nor *Tafa1*-expressing cells co-expressed *Vgat* in the pyramidal layer (Extended Data Fig. 4c), supporting their identity as glutamatergic rather than GABAergic neurons. Integrating these molecular and spatial annotations, we classified CA3-annotated nuclei into four distinct marker-defined groups: (1) an St18-positive cluster, comprising deep cells from the CA3-Glut1 supertype and a subset of CA3-Glut2 ventral cells; (2) St18-negative proximal CA3 neurons, corresponding to CA3-Glut4; (3) St18-negative mix CA3 neurons, comprising CA3-Glut4 proximal superficial, CA3-Glut3 distal superficial, and CA3-Glut2 ventral cells; and (4) a mossy cell cluster (Fig. 2h). Together, these results resolve CA3 PNs into molecularly and spatially distinct subtypes and identify *Tafa1* and *St18* as markers of superficial and deep CA3 PNs, respectively.

### Generation of a transgenic mouse line for identifying and targeting *St18*-expressing CA3 neurons

Having established *St18* as a marker of deep-layer CA3 PNs, we generated an *St18*-Cre mouse line to determine whether this gene delineates the previously described CA3 PN population characterized by sparse or absent TEs and burst-firing properties^6–10,308–12,32^. A T2A-Cre cassette was inserted immediately upstream of the endogenous *St18* stop codon, thereby placing Cre recombinase under the control of the endogenous *St18* promoter (Fig. 3a). To map the spatial distribution of *St18*-expressing cells, we injected P0 *St18*-Cre mice with AAV2/9-hSyn-DIO-NLS-tdTomato, resulting in nuclear tdTomato expression specifically in Cre-expressing cells (Fig. 3b).

**Figure 3.**
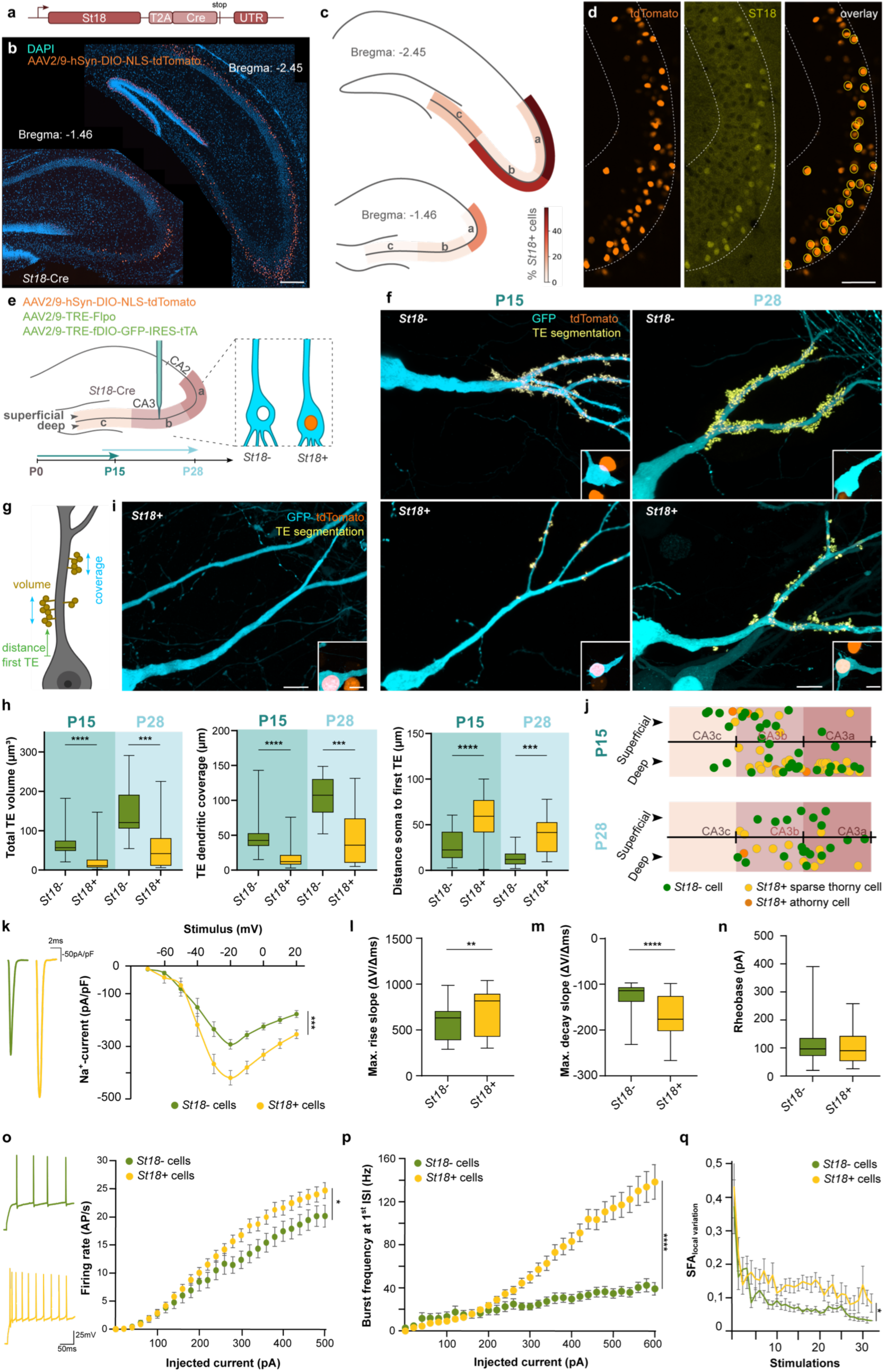
*St18* marks deep CA3 PNs with sparse or absent TEs and burst-firing properties. **(a)** Schematic of the *St18*-Cre allele. **(b)** Representative dorsal (left) and ventral (right) hippocampal sections showing Cre-dependent nuclear tdTomato labeling following injection of AAV2/9-hSyn-DIO-NLS-tdTomato into P0 *St18*-Cre mice. **(c)** Schematic distribution of *St18*-Cre-labeled cells along the proximodistal, deep-superficial, and dorsoventral axes of CA3. Color intensities indicate the percentage of *St18*-Cre-labelled cells (n=1-3 sections). **(d)** Representative images showing *St18*-Cre driven nuclear tdTomato labeling (orange) and endogenous ST18 immunoreactivity (yellow) in CA3. Yellow outlines identify ST18-immunopositive cells in overlay panel. **(e)** Strategy for sparse GFP labeling of CA3 PNs and Cre-dependent nuclear tdTomato identification of *St18*-positive neurons. **(f)** Representative apical dendrites of *St18*-negative (top) and *St18*-positive (bottom) CA3 PNs at P15 (left) and P28 (right), with segmented TEs overlaid in yellow. Insets show the corresponding somata and nuclear tdTomato labeling status. **(g)** Schematic of quantified morphological parameters. **(h)** Morphological quantification of *St18*-negative and *St18*-positive CA3 PNs at P15 and P28. (p-values P15/P28: total TE volume, p<0.0001/0.001; TE dendritic coverage, p<0.0001/0.0002; soma-to-first-TE distance, p<0.0001/0.0009. Mann– Whitney tests, except for TE dendritic coverage at P28 and soma-to-first-TE distance: at both ages Welch’s t-tests were used. P15: n=33 *St18*-negative and 42 *St18*-positive cells from 4 brains; P28: n = 21 *St18*-negative and 13 *St18*-positive cells from 5 brains). **(i)** Representative athorny *St18*-positive CA3 PN. Inset as in (f). **(j)** Normalized somatic positions of CA3 PNs along the proximodistal and deep-superficial axes at P15 and P28. Each point represents one neuron, classified as *St18*-negative (green), sparsely thorny *St18*-positive (yellow), or athorny *St18*-positive (orange). **(k)** Representative Na⁺ current traces (left) and Na^+^ currents normalized to cell capacitance (right) (two-way repeated-measures ANOVA, p=0.0005; n=43 *St18-*negative and 52 *St18*-positive cells). **(l, m)** Maximum action potential rise slope (l; p=0.003) and decay slope (m; p<0.0001) (Mann–Whitney tests; n=30 *St18*-negative and 38 *St18*-positive cells). **(n)** Measured rheobase (Mann-Whitney test, p=0.637; n= 48 *St18-* negative and 66 *St18*-positive cells). **(o)** Representative action potential firing traces from a single cell during current-clamp recordings (left) and firing rates in response to increasing depolarizing current injections (right; mixed-effects model, p=0.037; n= 55 *St18-*negative and 62 *St18*-positive cells). **(p)** First interspike interval (ISI) burst frequency to increasing depolarizing current (mixed-effects model, p<0.0001; n= 55 *St18-*negative and 62 *St18*-positive cells). **(q)** Local variation spike frequency accommodation corrected for rheobase differences (mixed-effects model, p=0.012; n=55 *St18-*negative and 62 *St18*-positive cells). Scale bars, 200 μm (b), 50 μm (d), 10 μm (f, i).

Analysis of serial brain sections spanning the dorsoventral hippocampal axis revealed a preferential localization of tdTomato-positive cells to deep-distal CA3 dorsally and to the deep mid-distal CA3 region more ventrally (Fig. 3c). The density of tdTomato-positive neurons gradually increased along the dorsoventral axis, reaching its highest levels in ventral CA3 (Fig. 3c). This observed spatial distribution of *St18*-Cre-labeled cells is consistent with the spatial organization of CA3-Glut1 (Fig. 2d) and the *St18* expression pattern confirmed by smFISH (Fig. 2g).

To confirm the specificity of the *St18*-Cre mouse line, we performed immunohistochemistry (IHC) with a previously validated anti-ST18 antibody (Huang C, Dev cell 2018) on hippocampal sections of AAV2/9-hSyn-DIO-NLS-tdTomato-injected *St18*-Cre mice. We found that every ST18-immunolabeled CA3 neuron also expressed nuclear tdTomato (Fig. 3d), demonstrating that *St18*-Cre-labeled cells express endogenous ST18. A small subset of tdTomato-positive neurons lacked detectable ST18 immunoreactivity (Fig. 3d), which is most likely explained by differences in labeling intensity between the bright tdTomato and weaker ST18 IHC signals. Together, these findings validate the *St18*-Cre mouse line as a reliable genetic tool for identifying and targeting *St18*-expressing CA3 PNs.

### *St18* expression marks sparsely thorny and athorny deep CA3 neurons

Taking advantage of the *St18*-Cre strategy to specifically access deep CA3 PNs, we next analyzed the morphology of these cells and compared them to superficial CA3 PNs. For sparse labeling and high-resolution morphological reconstruction of CA3 PNs, we used a Supernova-based system^33^ consisting of AAV2/9-TRE-Flpo and AAV2/9-TRE-fDIO-GFP-IRES-tTA vectors injected into *St18*-Cre mice (Fig. 3e). Leakage of the TRE promoter leads to stochastic expression of flippase (Flpo) and the tetracycline-transactivator (tTA), resulting in sparse but bright GFP labeling of both *St18*-positive and *St18*-negative neurons. Co-injection of a Cre-dependent nuclear tdTomato reporter enabled simultaneous visualization of *St18*-positive PNs (Fig. 3e, Extended Data Fig. 5a). Viral vectors were injected at P0 or P14, followed by morphological reconstruction and analysis of TEs at P15 or P28, respectively. During this postnatal period, MF synapses and TEs undergo substantial structural maturation^34^. We imaged both *St18*-positive and *St18*-negative cells and restricted our analysis to dorsal CA3, where CA3 PN morphology and circuit organization have been more extensively characterized.

*St18*-negative CA3 PNs displayed large, densely distributed TEs that occupied a substantial fraction of the apical dendrite, whereas *St18*-positive PNs exhibited smaller, more sparsely distributed TEs with reduced dendritic coverage at P15 (Fig. 3f-h, Extended Data Fig. 5b). TE volume and dendritic coverage increased in both populations at P28 (Fig. 3g-h). Despite this developmental increase in TE complexity, the differences in TE morphology between *St18*-negative and *St18*-positive PNs were maintained across ages (Fig. 3h), indicating that the reduced TE volume and coverage in *St18*-positive neurons is unlikely to reflect a transient delay in synapse maturation. Athorny CA3 PNs were found exclusively among *St18*-positive cells, comprising approximately 10% of this cell population across development (Fig. 3i, Extended Data Fig. 5c). *St18*-positive cells were primarily, but not exclusively, found in deep-distal CA3 (Fig. 3j).

In analyzing additional morphological parameters, we found that the first TE cluster on PN dendrites was located farther from the soma in *St18*-positive neurons compared to *St18*-negative ones, consistent with their deeper somatic position and lower TE density (Fig. 3h). We further examined proximodistal variation in TE architecture in dorsal CA3. Consistent with reported proximodistal differences in TE distribution^35^, *St18*-negative PNs showed a trend of increasing TE organization toward CA3a (Extended Data Fig. 5d). *St18*-positive PNs showed a similar but weaker trend, although interpretation was limited by sparse sampling in CA3c, reflecting their preferential distribution in CA3a/b. Together, these results characterize *St18*-expressing CA3 PNs as sparsely thorny or athorny.

### *St18*-positive CA3 neurons are characterized by a burst-firing phenotype

Previous studies identified preferential burst-firing in deep CA3 neurons with absent or sparse TEs^10,11^. We performed whole-cell patch clamp recordings to characterize the intrinsic electrophysiological properties of *St18*-positive and *St18*-negative CA3 PNs at P13-21, distinguished by respective presence or absence of nuclear tdTomato fluorescence following stereotactic injection of AAV-hSyn-DIO-NLS-tdTom in *St18*-Cre mice at P0.

*St18*-positive neurons were characterized by smaller membrane capacitance, input resistance, together with a shorter membrane time constant, whereas resting membrane potential was similar between populations (Extended Data Fig. 5e-h). Reduced input resistance and membrane time constant of *St18*-positive neurons would result in less responsiveness to small current inputs and return to baseline faster, inherently decreasing cell intrinsic excitability.

*St18*-positive neurons were characterized by an increased inward Na⁺ current, whereas the measured outward K⁺ current-density profiles were similar between the populations (Fig. 3k; Extended Data Fig. 5i, j). Furthermore, action potentials in *St18*-positive neurons were faster with shorter rise times/higher maximum upstroke slopes and faster repolarization kinetics (Fig. 3l, m; Extended Data Fig. 5k). However, no significant differences were detected in action-potential amplitude, voltage threshold, decay time, half-width, or afterhyperpolarization amplitudes (Extended Data Fig. 5l-q). Thus, *St18*-positive neurons differed in specific aspects of action-potential kinetics without detectable changes in overall spike amplitude or half-width. The larger inward Na⁺ current density may underlie a faster action potential upstroke.

While no significant difference in rheobase was detected (Fig. 3n), *St18*-positive neurons exhibited higher mean firing rates at stronger depolarizing current injections compared to *St18*-negative PNs (Fig. 3o), with a particularly pronounced increase in instantaneous firing frequency over the first interspike interval (ISI; Fig. 3p). Higher frequencies were also observed in measurements incorporating the first two or first three ISIs (Extended Data Fig. 5r). These findings support an enhanced propensity for rapid initial spike discharge. Moreover, *St18*-positive neurons exhibited greater variation in interspike intervals during prolonged train activity (i.e. irregular spiking, Fig. 3q). This suggests that firing behavior to suprathreshold stimulation, rather than cell intrinsic excitability, is different in *St*18-positive neurons. Together, *St18*-positive CA3 PNs are characterized by initial burst-firing, followed by an irregular activity profile, utilizing pertinent action-potential kinetics.

### Subtype-specific transcriptional and chromatin remodeling accompanies postnatal maturation of CA3 neurons

To begin identifying the molecular programs underlying the emergence of subtype-specific properties in CA3, we next examined how gene expression and chromatin accessibility change across postnatal development in CA3 PNs. To place each neuron on a continuous developmental axis, we fitted an ordinal regression model to predict developmental age (P5, P10, P15, or P28) from gene expression profiles and used the resulting developmental scores as pseudotime coordinates (see Methods) (Fig. 4a). As expected, cells from P5 were enriched at early pseudotime positions; P10 and P15 cells occupied intermediate positions, and P28 cells were shifted toward later pseudotime (Fig. 4a). In parallel, CA3 neurons were ordered along a molecular axis distinguishing St18-positive from St18-negative identities, reflecting a deep-superficial separation (Fig. 4a). St18-positive/deep and St18-negative/superficial CA3 neurons occupied opposite ends of this axis, supported by fitted expression of *St18* and *Tafa1* (Fig. 4b).

**Fig. 4.**
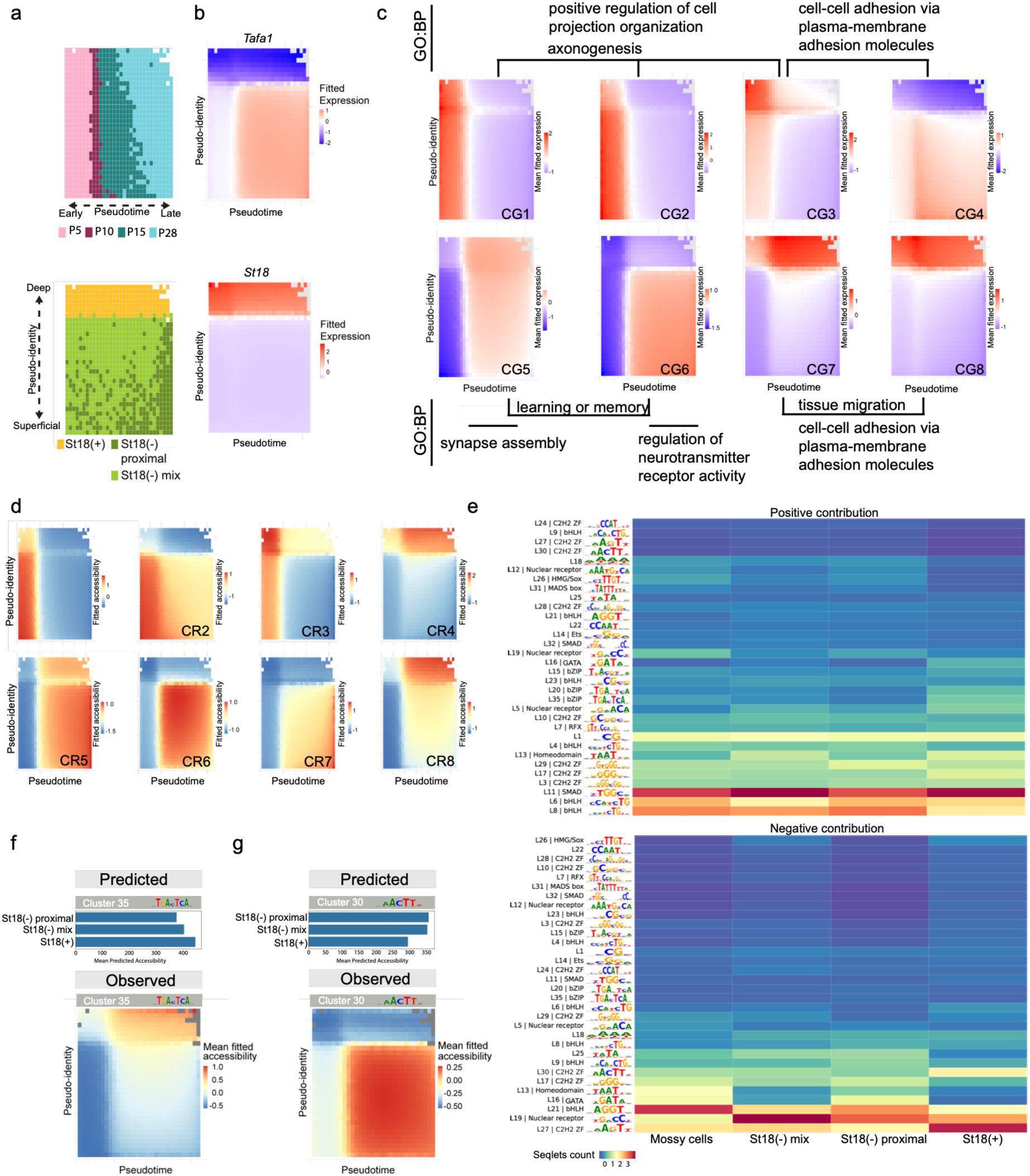
Developmental and identity-specific transcriptional and cis-regulatory programs in CA3 PNs. **(a)** Two-dimensional pseudotime-pseudoidentity representation of CA3 neurons, divided into 32 bins along each axis and colored by the predominant developmental stage (top) or CA3 PN subtype (bottom). **(b)** Generalized additive model (GAM)-fitted expression of *Tafa1* (top) and *St18* (bottom) across pseudotime and pseudo-identity. **(c)** Mean GAM-fitted expression profiles of eight dynamic gene clusters (CG1-CG8), with selected enriched Gene Ontology (GO) biological process terms. **(d)** Mean GAM-fitted chromatin accessibility profiles of eight dynamic regulatory region clusters (CR1-CR8). **(e)** Heatmaps of TF-MINDI seqlet-cluster abundance across CA3 PN subtypes, separated by positive (top) and negative (bottom) contribution scores. Rows are annotated with consensus motif logos and inferred DNA-binding domain (DBD) families. **(f, g)** Mean CREsted-predicted accessibility across CA3 PN subtypes (top) and GAM-fitted observed accessibility across pseudotime and pseudo-identity (bottom) for regions containing seqlets assigned to Leiden clusters 35 (f) and 30 (g), associated with AP-1-like and AAGTT/MYT-like motifs, respectively.

We then modeled gene expression across pseudotime and CA3 molecular identity and grouped genes with similar profiles into clusters, revealing coordinated programs associated with postnatal maturation, CA3 identity, or their interaction (Fig. 4c). Cluster genes (CG)1-CG3 were associated with early developmental states, with CG2 genes preferentially expressed in the St18-negative/superficial CA3 population and CG3 in St18-positive/deep CA3, indicating subtype-specific gene expression programs during early CA3 maturation. These clusters were enriched for cell projection organization, axonogenesis, and cell-cell adhesion (Fig. 4c, Extended Data Fig. 6a), consistent with early programs involved in neurite growth, axonal targeting, and circuit assembly^36,37^. CG4 genes were enriched at intermediate developmental stages and showed a predominantly superficial distribution. In contrast, late clusters CG5-CG8 were enriched for synapse assembly, learning or memory, regulation of neurotransmitter receptor activity, and cell-cell adhesion-related processes (Fig. 4c, Extended Data Fig. 6a), consistent with progressive synaptic maturation across postnatal CA3 development^38,39^. These programs also showed subtype-specific structure, with CG6 and CG7 genes preferentially associated with St18-negative and St18*-*positive CA3, respectively (Fig. 4c). Representative genes per cluster illustrated the identity and temporal profiles, including TFs such as *Trps1, Ar, Sox11*, and *Pou6f2*; guidance and adhesion molecules such as *Sema6d* and *Lrrtm4*; and ion channels such as *Kcnh7* (Extended Data Fig. 6b).

We next investigated whether these expression dynamics were mirrored by coordinated changes in chromatin accessibility. We modeled CA3 chromatin accessibility across the same pseudotime and identity coordinates, identifying dynamic cluster regions (CR)1-CR8 (Fig. 4d). Several region groups showed strong developmental dynamics. CR1-CR3 were most accessible at early pseudotime and decreased with maturation, whereas CR4-CR8 showed the opposite pattern. Several clusters (CR2-CR4, CR6-CR8) showed CA3 PN subtype-biased accessibility, indicating that chromatin state differs between St18-positive/deep and St18-negative/superficial CA3 populations.

Gene expression patterns of genes linked to CR-specific regions (Methods) broadly mirrored the corresponding accessibility dynamics (Extended Data Fig. 6c). CR1 regions, which lost accessibility with development, were linked to genes involved in cell projection organization, neurogenesis, and axon guidance, consistent with neurite growth and circuit assembly (Extended Data Fig. 6c, d). In contrast, CR5 regions gained accessibility over development and were linked to genes involved in membrane potential regulation, synaptic vesicle transport, postsynaptic organization, and synaptic plasticity (Extended Data Fig. 6c, d), consistent with maturation of neuronal signaling and synaptic function. Earlier work suggested that CA3 morpho-functional diversity is partly rooted in embryonic birth-timing, with earlier-born PNs preferentially occupying deeper positions within CA3a-b^38,40^. We therefore asked whether the molecularly defined deep and superficial CA3 populations identified here were associated with distinct embryonic time periods. We compared the transcriptional signatures of these deep and superficial CA3 populations with those of CA3 neurons labeled by *in utero* viral injection of AAV1-CAG-GFP at E14 and E17, followed by cell sorting at P30-40 and transcriptional profiling. The E14 injection is expected to label earlier-born CA3 neurons, whereas the E17 injection will label a mixed population of earlier-and later-born neurons. Genes enriched among E14-labeled neurons preferentially overlapped with the deep CA3 signature, whereas genes enriched among E17-labeled neurons showed greater overlap with superficial CA3 markers (Extended Data Fig. 6e-g). These findings support the notion that earlier-born neurons are enriched for the St18-positive/deep subtype. Taken together, these analyses show that CA3 PN maturation is accompanied by coordinated remodeling of chromatin accessibility and transcriptional programs across development in a subtype-specific manner.

### Subtype-specific *cis*-regulatory sequence features distinguish deep and superficial CA3

To identify sequence features underlying subtype-specific gene regulatory activity in CA3, we trained a sequence-to-function model using the CREsted^23^ peak regression framework to predict chromatin accessibility at accessible genomic regions (peaks) directly from their DNA sequence for each CA3 age-subtype group. The model learns regulatory sequence patterns associated with group-biased accessibility, including TF binding motifs, providing a framework to interrogate the *cis*-regulatory logic distinguishing CA3 groups across development. Model training followed a stepwise refinement strategy adapted from previous work^23,41^ to progressively increase CA3 group specificity (Extended Data Fig. 7a). The model was first trained on a broad hippocampal peak set, then fine-tuned on CA3-specific peaks, followed by peaks with variable accessibility across CA3 groups. A final fine-tuning step used age- and subtype-specific CA3 peaks to enrich for regulatory elements with restricted accessibility across CA3 subtypes. Predicted and observed accessibility were significantly correlated on held-out test regions (Pearson r=0.53, Extended Data Fig. 7b). Observed profiles within groups correlated more strongly between the same CA3 subtype across ages than between different subtypes within an age, indicating that subtype identity accounted for a major component of regulatory similarity. P5 was an exception, showing strong within-age correlations and suggesting a more general developmental state (Extended Data Fig. 7c). Correlations between predicted and observed group accessibility profiles broadly recapitulated this structure, indicating that the model captured the major organization of the accessibility landscape (Extended Data Fig. 7c).

We next quantified sequence contribution scores across the top regions in each regulatory cluster (CR1-CR8) for each age-subtype group (Fig. 4d). We applied Transcription Factor Motif Instance Neighborhood Decomposition and Interpretation (TF-MINDI)^42^ to extract and cluster short DNA sequences with high model attribution scores, known as seqlets, and relate them to known TF motifs. We then asked whether specific motif patterns were preferentially associated with CA3 subtypes. Seqlets were separated according to the sign of their contribution scores, representing predicted accessibility-increasing or accessibility-decreasing effects, and their abundance was compared across subtypes.

Among positively contributing seqlets, AP-1 motifs were more frequent in St18-positive/deep CA3 than in the other subtypes (Leiden (L) 28, L35; Fig. 4e). AP-1 factors, including JUN and FOS family proteins, are major mediators of activity-dependent transcription in neurons and have established roles in synaptic plasticity, neuronal maturation, and experience-dependent gene regulation^43,44^. Model-predicted accessibility for regions containing L35 cluster seqlets showed the highest predicted accessibility in St18-positive CA3 subtypes, consistent with their observed accessibility profiles (Fig. 4f). A nuclear receptor motif (L5) was enriched in St18-positive and St18-negative mix populations compared with superficial proximal CA3 and mossy cells (Fig. 4e). A zinc finger motif (L29) was preferentially associated with the St18-negative proximal subtype (Fig. 4e), which is consistent with the higher predicted and observed accessibility of regions containing L29 seqlets in superficial CA3 subtypes (Extended Data Fig. 7d).

Among negatively contributing seqlets, which are associated with reduced model-predicted accessibility, AAGTT-like motifs (L30, L27) were enriched in St18-positive/deep CA3 (Fig. 4e). These motifs closely resemble the consensus sequence reported for MYT-family neural zinc finger TFs including ST18, MYT1L and MYT1^25,45^. This negative contribution could reflect reduced accessibility in St18-positive/deep CA3 or, alternatively, stronger accessibility associated with the same motif in St18-negative subtypes. Regions containing AAGTT/MYT-like motifs showed lower predicted accessibility in St18-positive than in St18-negative CA3 neurons (Fig. 4g). This pattern was recapitulated in the observed accessibility landscape, with lower accessibility in deep CA3 and higher accessibility in the superficial domain (Fig. 4g).

To relate the AAGTT/MYT-like motif signature to candidate MYT-family TFs, we compared TF expression with accessibility at these regions. Accessibility at AAGTT/MYT-like motif regions was strongly anti-correlated with *St18* expression, with the lowest accessibility observed in *S*t18-positive/deep CA3 neurons (Extended Data Fig. 7e). *Myt1l* expression showed a weaker negative correlation and was broadly expressed across CA3 populations, whereas *Myt1* showed little to no correlation, and no clear subtype-specific pattern. Together, these correlations link the AAGTT/MYT-like motif signature most closely to *St18* and support a role for ST18 in restricting accessibility at deep CA3 regulatory elements.

### Regulatory network inference predicts ST18-mediated repression of superficial CA3 programs

Having identified subtype-specific *cis*-regulatory sequence features and accessibility patterns, we next asked how these differences are organized into broader gene regulatory programs across CA3 subtypes. We therefore used SCENIC+^46^ to infer subtype-specific regulatory programs. As no canonical ST18 motif is available, we incorporated the TF-MINDI-derived AAGTT/MYT-like motif as a custom motif annotated for ST18 and MYT1L. Several enhancer-driven regulons (eRegulons) showed subtype-specific activity across CA3 populations, broadly mirrored by expression of their corresponding TFs (Fig. 5a, b). Among activating eRegulons, *Neurod6*- and *E2f3*-linked programs were preferentially active in St18-negative proximal CA3, whereas *Ar*-, *Nr3c2*-, and *Rfx3*-linked programs were enriched in St18-positive CA3 (Fig. 5a). *Pou6f2* and *Trps1* were enriched in St18-negative proximal CA3 and were associated with both activating and repressive eRegulons within this population. *St18* was predicted to act as a repressor in the St18-positive/deep CA3 subtype, targeting genes preferentially expressed in St18-negative CA3 PNs (Fig. 5a, b).

**Fig. 5.**
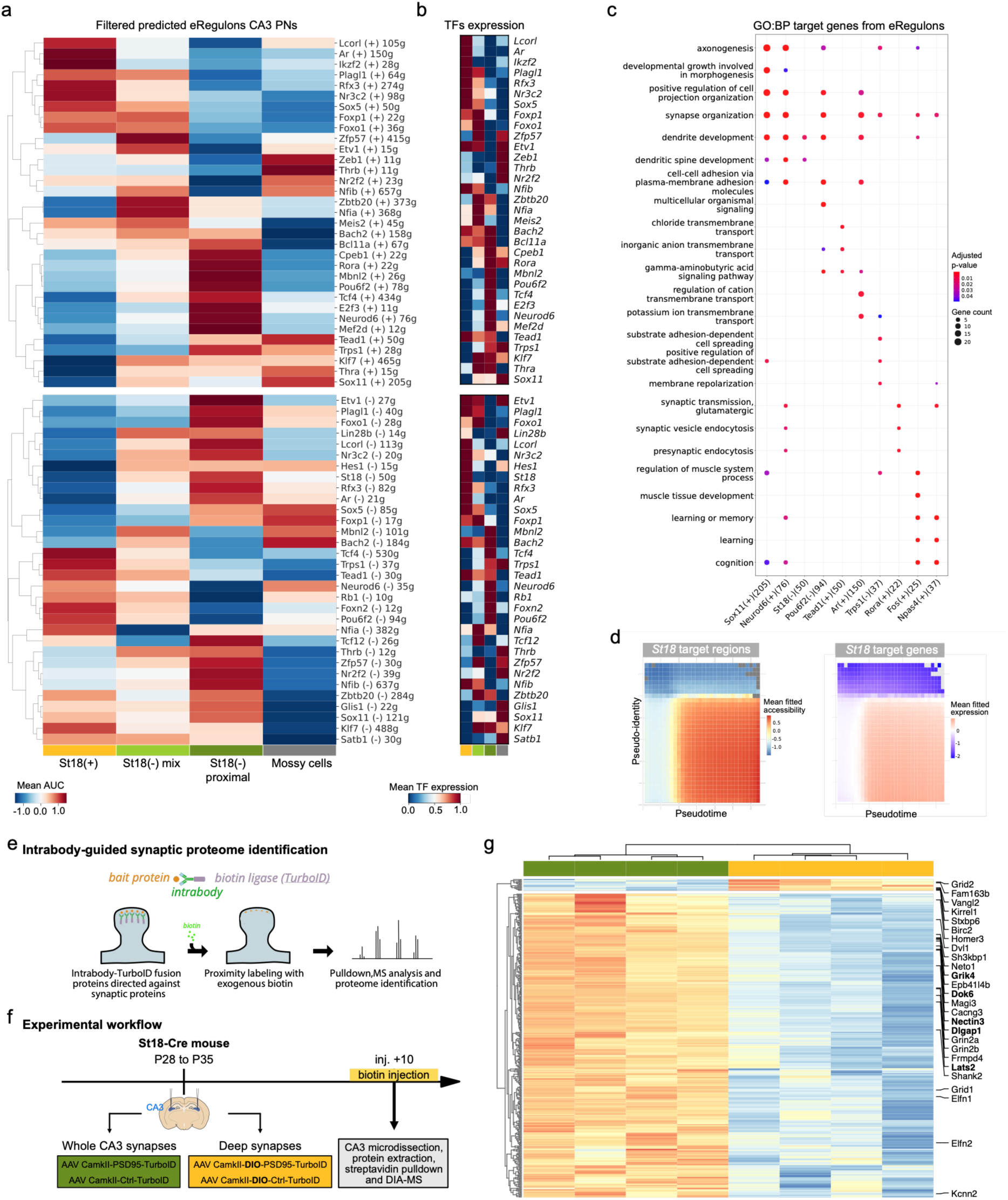
Subtype-specific gene regulatory networks distinguish CA3 pyramidal neuron populations. **(a)** Heatmap of mean gene-based eRegulon activity across CA3 subtypes for selected (see Methods) activating (+) and repressive (-) eRegulons inferred using SCENIC+. Activity was measured using area under the recovery curve (AUC) scores, reflecting the enrichment of predicted target genes within each cell’s expression profile. The number of predicted target genes is indicated for each eRegulon. **(b)** Heatmap showing mean expression of the corresponding transcription factors (TFs) across CA3 subtypes. Colors indicate scaled mean expression. **(c)** Gene Ontology (GO) biological process enrichment of target genes from selected CA3 eRegulons. **(d)** Mean generalized additive model (GAM)-fitted chromatin accessibility of predicted *St18*-linked regulatory regions (left) and expression of their target genes (right) across pseudotime and pseudo-identity. **(e)** Schematic of intrabody-TurboID-mediated postsynaptic protein biotinylation. **(f)** Workflow of intrabody-TurboID in CA3 PNs (n=4 biological replicates per condition, 3/4 animals per replicate, a mix of males and females). **(g)** Row-based hierarchical clustering of 111 differentially enriched proteins (DEPs) shown as a z-scores heatmap (limma moderated test BH FDR=0.05, LFC thresholds are 1 for synaptic vs. control conditions and 0.5 between synaptic conditions).

We then used gene set enrichment analysis to link predicted regulators to the dynamic gene programs (CG1-CG8) (Extended Data Fig. 8a, b; see Fig. 4a-c). Early gene clusters (CG1-CG2) were enriched for eRegulons led by *Sox11, Klf7, Bach2, Nfia, Zfp57, Lcorl,* and *Rfx3*, consistent with the decline of these TFs from P5 to P28 (Extended Data Fig. 2). In contrast, *Npas4*-and *Fos*-linked eRegulons were enriched in the late cluster CG5, consistent with activity-dependent gene regulation^43^. Additional eRegulons were associated with gene clusters that varied along the deep-superficial CA3 axis. Target genes of the activating *Ar*-and *Nr3c2*-linked eRegulons were enriched in CG7 and CG8, both of which include genes with higher expression in deep CA3 neurons (Extended Data Fig. 8a). Similarly, genes predicted to be repressed by *Pou6f2* and *Trps1* were enriched in the deep CA3-associated clusters CG3, CG7, and CG8 (Extended Data Fig. 8b).

GO enrichment analysis further revealed functional differences among these temporally- and identity-associated eRegulons (Fig. 5c). *Sox11* target genes are involved with roles in early postnatal circuit formation such as axonogenesis, developmental growth involved in morphogenesis, and positive regulation of cell projection organization (Fig. 5c). Target genes of several CA3-specific regulatory programs, such as *Ar*-activating and *Pou6f2*-and *Trps1*-repressive eRegulons, were enriched for synapse organization, cell-cell adhesion, and ion transport and signaling, suggesting that these programs may contribute to the differential connectivity of CA3 populations. In contrast, target genes of later-expressed activity-dependent TFs, including *Npas4 and Fos*, were enriched for learning, cognition, and glutamatergic synaptic transmission (Fig. 5c), indicating a transition toward mature synaptic and experience-dependent neuronal functions.

Predicted ST18 regions and target genes are more active at later pseudotime in the superficial CA3 domain (Fig. 5d). Predicted ST18 target genes are enriched for dendrite development and dendritic spine development terms (Fig. 5c) and comprised several functional gene classes (Extended Data Fig. 8c), including cell-surface or synapse-associated proteins such as *Grik4, Nectin3, Chrm3, Lsamp*, and *Cntnap2*; ion channels (*Kcnq5, Kcnc2, Cacna2d3*); TFs (*Pou6f2, Trps1*), and protein kinases (*Lats2, Map3k5, Sgk1, Srpk2*). Several predicted targets were associated with St18-negative/superficial CA3 identity, including *Tafa1* and *Kcnq5.* The predicted target genes *Nectin3* and *Grik4* have established roles in MF-CA3 synaptic organization and function^47,48^, whereas *Kcnq5, Kcnc2, Cacna2d3,* and *Rimbp2* are related to voltage-gated ion channel activity^49–52^. These findings suggest that ST18 may regulate key genes related to CA3 identity, synaptic organization and function, and electrophysiological properties.

### CA3 subtypes are characterized by distinct synaptic signatures

To investigate how the inferred regulatory programs relate to synaptic protein composition, and whether predicted ST18 targets are represented at excitatory postsynaptic sites, we compared the postsynaptic proteomes of St18-positive CA3 PNs and the broader CA3 population. To characterize synaptic proteome composition in a cell-type-specific manner, we applied our intrabody-guided synaptic proximity labeling method^53^that fuses the biotin ligase TurboID to an intrabody targeting the excitatory postsynaptic scaffold protein PSD95 (Figure 5e). We opted for a differential enrichment approach, comparing the postsynaptic composition of *St18*-Cre-labeled CA3 PNs to that of whole CA3 PNs. Because of the higher relative proportion of superficial CA3 PNs compared to deep CA3 neurons, we reasoned that this approach would be sufficient to represent deep vs. superficial synaptic proteome differences.

We stereotactically injected PSD95-TurboID- and Ctrl-TurboID-expressing AAVs in the CA3 region of *St18-*Cre mice, using either constitutive or Cre-dependent constructs (Figure 5f, Extended Data Fig. 8d), to target whole CA3 or *St18*-Cre CA3 postsynaptic proteomes, respectively. Ten days after, mice were treated with biotin for 3 hours, and CA3 was subsequently dissected. Biotinylation levels were similar between *St18*-Cre and whole CA3 samples (Extended Data Figure 8e). After streptavidin-mediated pulldown and proteomic sample preparation^54^, we used data-independent acquisition mass spectrometry (DIA-MS) to identify synaptic and control proteomes (Extended Data Figure 8f-h) and performed PCA and differential enrichment analysis (Extended Data Figure 8i, see Methods). Comparing control proteomes between the two cell populations demonstrated the enrichment of cell-type specific markers in the correct condition (e.g. St18 in the *St18*-Cre condition and Trps1, a superficial CA3-enriched gene, in the whole CA3 condition; Extended Data Fig. 8j), indicating that our experimental strategy can differentiate proteomic composition of deep and superficial PNs. Analysis comparing synaptic conditions against their respective controls yielded 426 synaptic proteins, of which 111 were differentially enriched in the *St18*-Cre population compared to whole CA3 PNs (Fig. 5g).

We found distinct synaptic signatures for *St18*-Cre and whole CA3 samples. Whole CA3 was enriched in cell-surface proteins such as Nectin3 and Elfn1, kainate receptor and auxiliary subunits Grik4 and Neto1, and scaffold proteins such as Homer3 and Shank2, which are present at MF-CA3 synapses^55^ (Figure 5g). In contrast, *St18*-Cre CA3 PN synapses largely lacked this MF-CA3 synapse signature. These synapses were enriched in cell-surface proteins Kirrel1 and Vangl2 (Fig. 5g), of which Vangl2 has been shown to regulate MF-CA3 connectivity and function^56^. Interestingly, the glutamate receptor delta proteins Grid1 and Grid2 were differentially expressed between the two populations (Fig. 5g).

We next compared predicted ST18 target genes (Extended Data Figure 8c) with the synaptic proteins that were differentially enriched between whole CA3 and *St18*-Cre CA3. Five predicted ST18 targets, *Dok6*, *Lats2*, *Grik4*, *Nectin3* and *Dlgap1*, overlapped with synaptic proteins enriched in whole CA3 (Figure 5g, indicated in bold), whereas none overlapped with the *St18*-Cre-labeled synaptic proteome. These findings suggest that part of the molecular program repressed by ST18 is related to superficial CA3 PN synaptic organization and function. Taken together, the results from our gene regulatory and proteomic analyses define subtype-specific regulatory programs across CA3 and suggest that ST18 restricts superficial CA3-associated molecular and synaptic programs in deep CA3 neurons.

### Postnatal loss of ST18 shifts deep CA3 molecular identity toward an *St18*-negative state

Given the enrichment of *St18* in sparsely thorny and athorny deep CA3 PNs and its predicted repression of superficial CA3-associated genes and regulatory programs, we reasoned that ST18 may actively restrict superficial-like molecular, structural and physiological features. We therefore examined the consequences of ST18 loss in deep CA3 PNs.

To induce postnatal loss of ST18, we injected the CA3 region of P0 *St18*-Cre:H11-Cas9 mice with an AAV carrying an hU6-driven gRNA targeting exon 10 of *St18* (AAV- hU6-ST18KO-fDIO-GFP-IRES-tTA) (see Methods) (Fig. 6a). Additionally, the KO construct contained a Flpo-dependent GFP expression cassette, used to assess injection efficiency when co-injected with AAV-TRE-DIO-FLP, and an IRES-tTA sequence for sparse labeling of neurons. Control mice received the corresponding AAV lacking the *St18*-targeting gRNA. These conditions are referred to as *St18*-KO and *St18*-CTRL, respectively. IHC showed a robust reduction in ST18 immunoreactivity in CA3 PNs at P15, confirming efficient loss of ST18 protein in *St18*- KO animals (Fig. 6b).

**Fig. 6.**
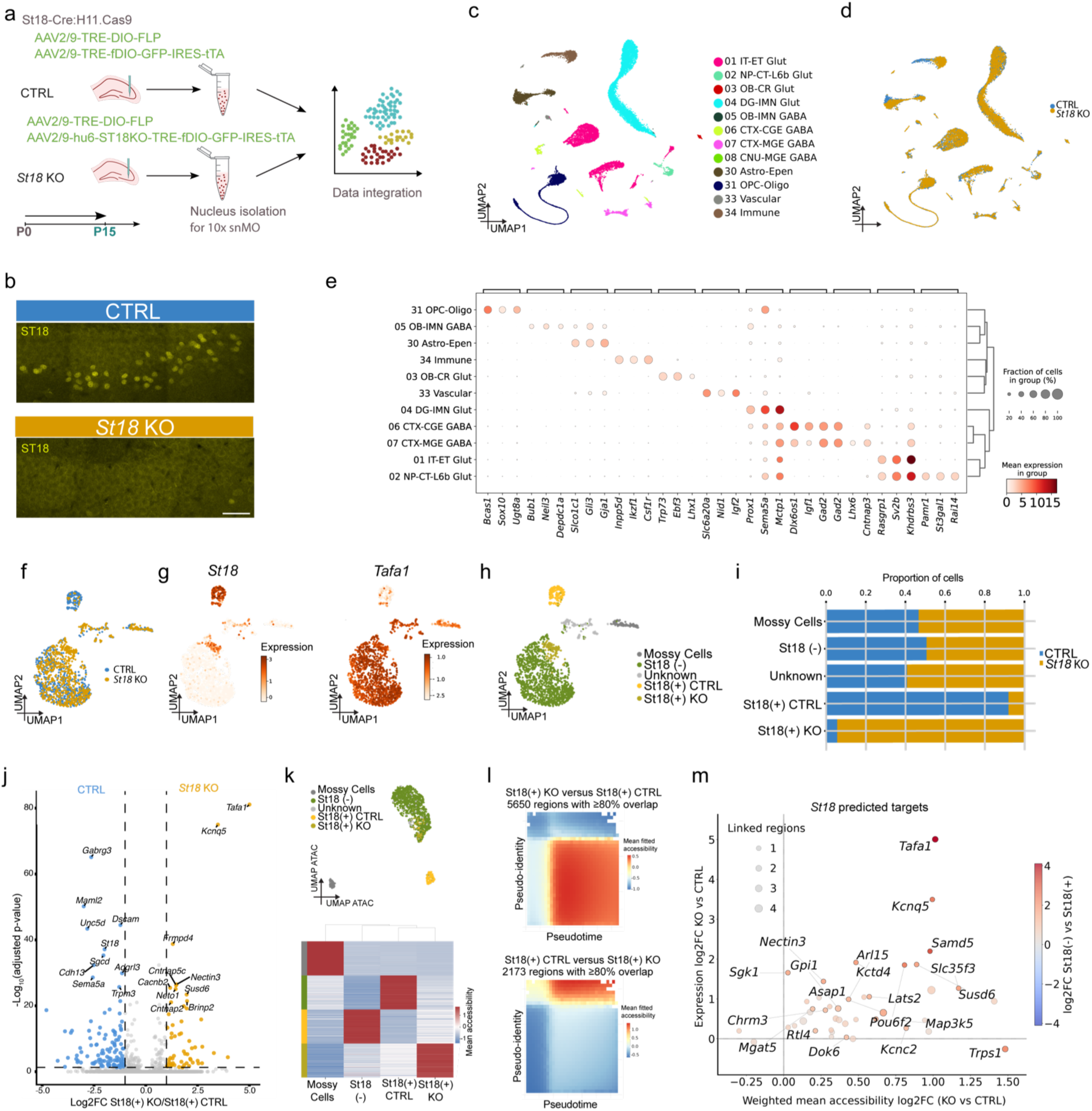
Postnatal ST18 loss remodels the molecular and chromatin accessibility landscape of deep CA3 PNs. **(a)** Schematic of the experimental strategy for postnatal *St18* knockout (KO) in *St18*-Cre:H11-Cas9 mice. AAVs carrying Flpo and either a control construct or an *St18*-targeting gRNA were injected into CA3 at P0, followed by snMO profiling at P15. **(b)** Representative IHC images showing reduced ST18 immunoreactivity in *St18* KO compared to CTRL. Scale bar, 50 µm. **(c, d)** UMAP of integrated snMO profiles colored by major cell class following label transfer from the Allen Brain Cell (ABC) Atlas reference (c) or experimental condition (d). **(e)** Dot plot of the top differentially expressed genes across major cell classes (Wilcoxon rank-sum test). Dot size represents the percentage of cells expressing each gene, and color indicates mean log-normalized expression. **(f)** UMAP of reclustered CA3 neurons colored by experimental condition. **(g)** UMAP showing expression of *St18* and *Tafa1* in reclustered CA3 neurons. **(h)** UMAP of CA3 neurons annotated as mossy cells, St18- negative, St18-positive CTRL, St18-positive KO and unknown populations. **(i)** Relative proportions of CTRL and KO nuclei across CA3 populations. **(j)** Volcano plot showing differential gene expression between *St18*-positive KO and CTRL neurons (MAST test). Positive log₂ fold changes (LFC) indicate increased expression following ST18 loss. **(k)** UMAP of CA3 chromatin accessibility profiles colored by RNA-seq based CA3 population annotations (top) and heatmap of mean accessibility of the top differentially accessible regions across CA3 populations (bottom). **(l)** Mean generalized additive model (GAM)-fitted chromatin accessibility profiles across pseudotime and pseudo-identity for regions gaining (top) or losing accessibility (bottom) following ST18 loss, identified by differential accessibility analysis between St18-positive KO and CTRL neurons (Wilcoxon rank-sum test). **(m)** Relationship between gene expression changes and weighted mean chromatin accessibility changes across linked regulatory regions for predicted *St18* target genes following ST18 loss. Both axes show LFC between St18-positive KO and CTRL neurons. Dot size indicates the number of linked regulatory regions, and color indicates the gene expression LFC between St18-negative and St18-positive CA3 neurons in the unperturbed dataset.

To define the molecular and regulatory changes associated with ST18 loss, we performed hippocampal snMO profiling at P15 following *St18* KO. We assigned cell identities by transferring labels from the ABC Atlas reference^24^ resolving glutamatergic, GABAergic, oligodendroglial, astrocytic, vascular and immune subclasses. This analysis captured the major neuronal and non-neuronal populations in both *St18*-CTRL and *St18*-KO samples (Fig. 6c). *St18*-CTRL and *St18*-KO nuclei showed a similar overall distribution in the integrated UMAP (Fig. 6d), and expression of the top three differentially expressed genes per major cell class supported the transferred annotations (Fig. 6e). To assess the effects of ST18 loss specifically in CA3 PNs, we isolated and reclustered CA3 cells (Fig. 6f, Extended Data Fig. 9a-c). Reclustering resolved two transcriptionally distinct St18-positive populations enriched for CTRL and KO cells, respectively (Fig. 6f, g). Our single-guide KO strategy is expected to induce indels leading to loss of function at the protein level, without extensive genomic deletions. *St18* mRNA is therefore still detected in our snMO data, in both CTRL and KO cells (Fig. 6g).

Notably, St18-positive KO cells expressed *Tafa1*, which in CTRL was mainly restricted to St18-negative CA3 neurons (Fig. 6g). We therefore resolved four CA3 populations: mossy cells, St18-negative cells, St18-positive CTRL and St18-positive KO cells, together with a small population of unassigned CA3 cells (Fig. 6h, i). The mossy cell and St18-negative clusters contained similar proportions of CTRL and KO cells, whereas the two St18-positive clusters showed a strong condition bias (Fig. 6i). The unassigned population, labeled as “unknown,” contained both CTRL and KO nuclei with heterogeneous *St18* expression, preventing their confident assignment to either CA3 subtype.

Differential expression analysis between St18-positive CTRL and KO CA3 neurons identified 56 upregulated and 115 downregulated genes following ST18 loss (Fig. 6j). Among the most strongly upregulated genes were *Tafa1* and *Kcnq5*, whose expression is normally enriched in superficial CA3 and complementary to *St18* (Fig. 2e-g, Extended Data Fig. 4b)^13^. *Nectin3* and *Neto1*, genes implicated in MF-CA3 synaptic organization and function^47,57^, were also upregulated following ST18 loss. The expression of several genes normally enriched in deep CA3 was reduced (Fig. 6j), including *Unc5d, Gabrg3, Dscam, Cdh13* and *Sema5a*, which encode cell-surface proteins involved in inhibitory synaptic transmission, adhesion and axon guidance^58–62^. Together, these changes indicate broad remodeling of the molecular program of deep CA3 neurons following ST18 loss. In contrast, analysis of dentate granule cells and CA1 PNs revealed no major transcriptional changes following ST18 loss (Extended Data Fig. 9d-m).

Given the predicted repressive role of *St18* in deep CA3 PNs, we next examined chromatin accessibility following ST18 loss. St18-positive CTRL and KO neurons showed distinct accessibility profiles (Fig. 6k, Extended Data Fig. 9n-p). Regions gaining accessibility in St18-positive KO cells were preferentially associated with the superficial CA3 domain, whereas regions losing accessibility mapped to the deep domain (Fig. 6l). These changes are consistent with a shift toward a more superficial-like regulatory state in *St18*-KO neurons.

We then asked whether predicted ST18-linked regulatory regions and their target genes were affected by ST18 loss. For each predicted target gene, we compared the expression change between St18-positive KO and CTRL neurons with the weighted mean accessibility change across its linked regulatory regions (Fig. 6m). Several targets showed concordant increases in accessibility and expression following ST18 loss. These targets included *Tafa1*, *Nectin3*, *Kcnq5*, *Samd5* and *Susd6*, consistent with their preferential expression in St18-negative CA3 neurons in the unperturbed dataset (Fig. 6m). This indicates that ST18 loss remodels the inferred regulatory network with coordinated changes in chromatin accessibility and transcription. Taken together, these changes support a role for ST18 in maintaining deep CA3 identity by restricting a superficial CA3-associated regulatory program.

### ST18 loss shifts deep CA3 PNs toward superficial-like morphology and firing properties

To determine whether the shift toward a superficial-like molecular identity in *St18* KO neurons translates into changes in morphology, we reconstructed TEs of sparsely labeled *St18-*Cre-expressing PNs in *St18-*CTRL and *St18*-KO conditions (Fig. 7a). We selected neurons with an intact apical dendritic tree primarily located in distal CA3 of the dorsal hippocampus (Fig. 7b, Extended Data Fig. 10a). Consistent with our previous observations (Fig. 3f-h), *St18-*CTRL CA3 PNs exhibited small and sparsely distributed TEs with limited dendritic coverage (Fig. 7c-e). In contrast, *St18-*KO cells exhibited larger and more densely distributed TEs that occupied a greater fraction of the apical dendrite (Fig. 7c-e). TE volume and dendritic coverage in *St18-*KO cells approached those of St18*-*negative cells at the same developmental time point (Fig. 7e, dotted lines). These findings indicate that ST18 loss results in a shift toward a superficial CA3-like TE architecture.

**Figure 7.**
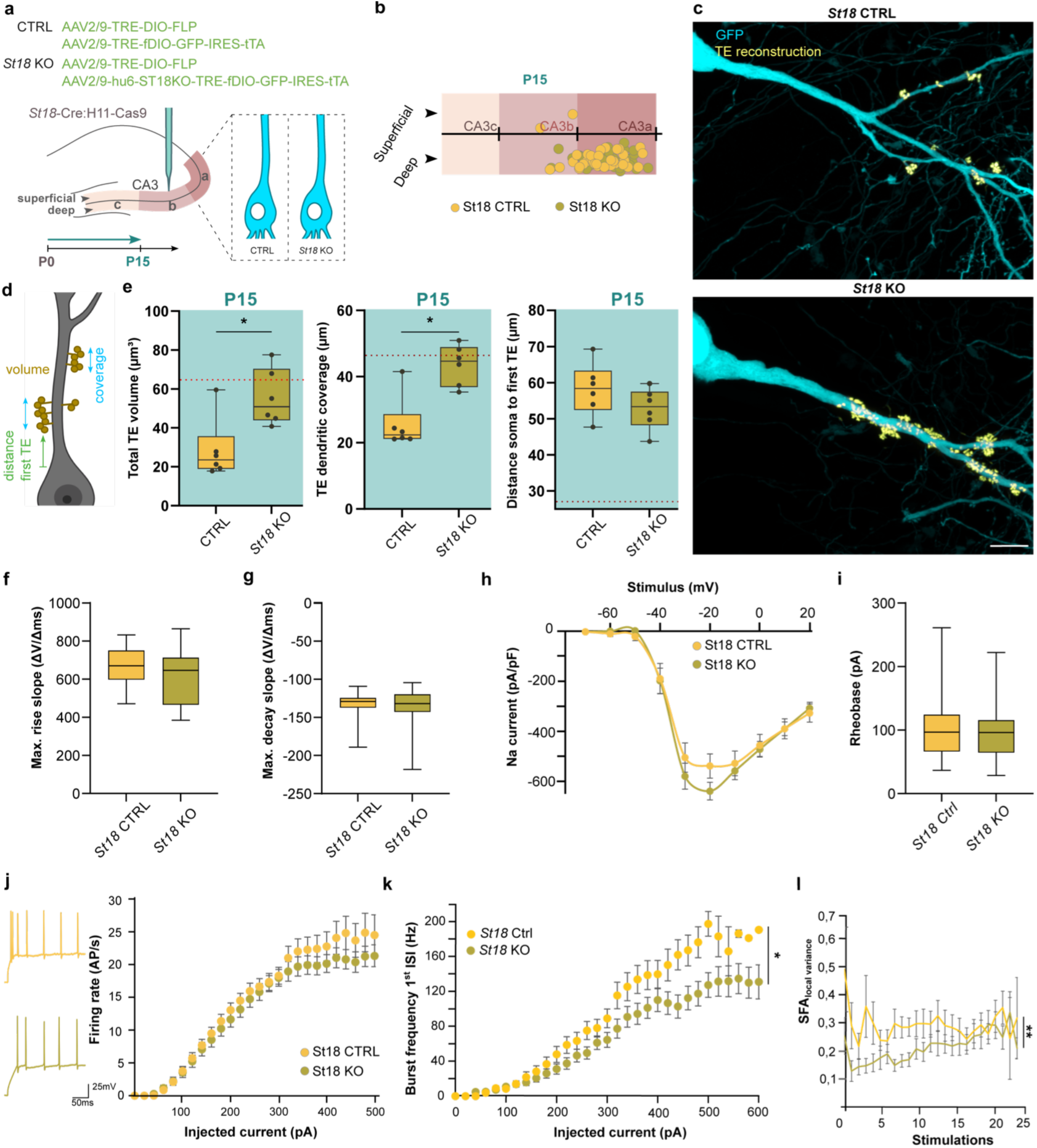
Postnatal loss of ST18 promotes a superficial-like TE identity and reduces burst-firing in CA3 PNs. **(a)** Experimental strategy for CRISPR/Cas9-mediated *St18* KO and Cre-dependent sparse labeling in *St18*-Cre:H11-Cas9 mice. **(b)** Normalized somatic positions of analyzed CA3 PNs along the proximodistal and deep-superficial axes at P15. Each point represents one neuron, classified as *St18*-CTRL (green) or *St18*-KO (yellow). **(c)** Representative images of *St18*-CTRL (top) or *St18-*KO (bottom) CA3 PNs. TE segmentation is shown in yellow. Scale bar 10 µm. **(d)** Schematic of morphological parameters quantified. **(e)** Morphological quantification of *St18-*CTRL and *St18*-KO CA3 PNs at P15: total TE volume (Welch’s t-test, p=0.0114), TE dendritic coverage (Welch’s t-test, p=0.0016) and the distance from soma to first TE (Student’s t-test, p=0.1826) (n=6 animals per condition). The dotted line indicates the mean value of *St18*-negative cells shown in Fig. 3h. **(f, g)** Maximum action potential rise slope (f; p=0.171) and decay slope (g; p=0.880) (Mann–Whitney tests; n=23 *St18*-CTRL and 30 *St18*-KO cells). **(h)** Na^+^ current traces normalized to cell capacitance (two-way repeated-measures ANOVA, p=0.543; n= 38 *St18*-CTRL and 46 *St18*-KO cells). **(i)** Measured rheobase (Mann-Whitney test, p=0.768; n= 28 *St18*-CTRL and 35 *St18*-KO cells). **(j)** Representative action potential firing traces from a single cell during current-clamp recordings (left) and firing rates in response to increasing depolarizing currents (right; mixed-effects model, p=0.397; n= 26 *St18*-CTRL and 39 *St18*-KO cells). **(k)** First interspike interval (ISI) burst frequency to increasing depolarizing current (mixed-effects model, p=0.014; n= 39 *St18*-CTRL and 30 cells *St18*-KO cells). **(l)** Local variation spike frequency accommodation corrected for rheobase differences (mixed-effects model, p=0.006; n= 30 *St18*-CTRL and 39 *St18*-KO cells).

To examine whether loss of ST18 also affects the intrinsic electrophysiological properties of deep CA3 PNs, we performed whole-cell recordings from *St18*-CTRL and *St18*-KO PNs and compared their passive membrane properties, voltage-gated currents, action-potential kinetics, and firing responses. Comparison of membrane capacitance, resting membrane potential, input resistance, and membrane time constant revealed no differences between *St18-*CTRL and *St18*-KO PNs (Extended Data Fig. 10b-e). Thus, passive membrane properties were not altered upon ST18 loss. Action potentials in *St18*-KO PNs had significantly longer rise time (Extended Data Fig. 10f), whereas no significant differences were detected in maximum upstroke slopes and repolarization kinetics (Fig. 7f-g). There were also no significant changes in amplitude, voltage threshold, decay time, half-width, or afterhyperpolarization amplitudes (Extended Data Fig. 10g-l). Moreover, inward Na^+^ and outward K^+^ current densities were similar between *St18-*KO and *St18-*CTRL cells, suggesting that voltage-gated currents are not affected upon ST18 loss (Fig. 7h, Extended data Fig. 10m, n).

Analysis of firing responses to depolarizing current injection revealed no significant difference in rheobase or mean firing rate or between *St18*-CTRL and *St18-*KO PNs (Fig. 7i, j). However, *St18*-KO neurons exhibited significantly lower instantaneous firing frequency over the first interspike interval (ISI; Fig 7k), as well as lower frequencies measured across the first two or first three ISIs (Extended Data Fig. 10o). These findings indicate reduced burst responses upon ST18 loss. Moreover, *St18*-KO neurons exhibited lower spike variance accommodation (SFA), consistent with more regular spike timing (Fig. 7l). Together, these results indicate a role for ST18 in controlling rapid initial and sustained firing properties, potentially by tuning action potential kinetics in CA3 PNs. Loss of ST18 selectively altered these properties rather than completely shifting electrophysiogical properties toward the *St18*-negative state.

## Discussion

Here, we integrated developmental single-nucleus multiomics with cellular and loss- of-function analyses to identify distinct gene regulatory programs associated with deep and superficial CA3 and established the TF ST18 as a regulator that restricts superficial-like molecular, morphological, and electrophysiological features in deep CA3 neurons.

CA3 PNs exhibited substantial molecular heterogeneity throughout postnatal development. Extending earlier descriptions of distinct CA3 expression domains along the proximo-distal and septotemporal axes^13^, our analysis identifies the radial axis as a notable source of molecular variation. The segregation of deep *St18*-expressing and superficial *Tafa1*-expressing populations was already apparent at P5, suggesting that radial molecular identity is established before the completion of CA3 circuit maturation. Birth timing may contribute to this organization, as temporally distinct populations of CA3 neurons can acquire different molecular identities and connectivity patterns along the radial axis^38,40^. Early-generated neurons are enriched in deep CA3a/b, while later-born neurons are more evenly distributed across the pyramidal layer. Consistent with this, genes enriched in CA3 neurons embryonically labeled by viral injection at E14, relative to the heterogeneous population labeled at E17, were preferentially expressed toward the St18-positive/deep end of our molecular pseudo-identity axis. These findings suggest that *St18*-expressing CA3 PNs are generated earlier than superficial CA3 PNs.

To link the molecular identity of *St18*-positive neurons to previously characterized cellular properties of neurons in deep CA3, we generated an *St18*-Cre line. We found that *St18*-expressing neurons were predominantly sparsely thorny, with a small (∼10%) athorny fraction. *St18* thus marks a broader deep CA3 population that includes both athorny and sparsely thorny neurons, consistent with recent work describing both morphologies in deep CA3^11,12^. Electrophysiological recordings from *St18*-expressing neurons revealed the burst-firing pattern characteristic of deep CA3 PNs^8,10,11^. During the preparation of this manuscript, two independent preprints also identified *St18* as a marker of this deep CA3 population and reported reduced TE coverage and burst-firing properties for these cells^63,64^. Together, these studies confidently link *St18* expression to a deep CA3 population previously characterized by its morphological and electrophysiological properties.

Previous work showed that athorny CA3 PNs lack functional MF input^10^. In sparsely thorny CA3 PNs, serial-section electron microscopy showed that TEs were contacted by vesicle-rich MF boutons^12^. Whether these contacts support functional transmission, and how this compares with densely thorny cells, remains to be determined. It is also unknown whether the same granule cells innervate deep and superficial CA3 PNs or whether distinct granule cell populations preferentially target sparsely or densely thorny cells. Birth timing could contribute to such selectivity, given evidence for preferential connectivity between temporally matched granule cells and CA3 neurons65. Finally, whether deep CA3 PNs receive functional MF input will also be important to determine in the context of refining CA3 circuit models, in which information flows from granule cells to superficial CA3 neurons, and from there to neurons in deep CA3 and onward to CA1^11^. The functional relevance of a direct granule cell to deep CA3 connection in this two-layer network remains unclear.

Beyond MF input, recurrent excitatory connections and inhibitory inputs also appear to be organized along the CA3 radial axis^11,12,32^. Electrophysiological recordings and connectomic studies indicate that deep and superficial CA3 PNs receive partially segregated inhibitory inputs, with inhibitory axons preferentially targeting one sublayer^12^. Similar radial differences in inhibitory connectivity have been described in CA1, where parvalbumin (PV)-positive basket cells preferentially innervate deep PNs^66^. This differential innervation is regulated in part by the TF SATB2, which suppresses PV-positive inhibitory synapse formation onto superficial CA1 PNs during development^67^. In cortical L5, CDH13 promotes PV-positive interneuron innervation of deep extratelencephalic neurons^68^. These examples suggest a putative framework that links subtype-specific inhibitory input to transcriptional regulation and cell-surface recognition molecules. The enrichment of *Gabrg3*, *Kirrel3*, and *Cdh13* in deep CA3 PNs in our study suggest that related mechanisms could contribute to their distinct inhibitory connectivity.

Modeling gene expression and chromatin accessibility across pseudotime and molecular identity revealed that deep and superficial CA3 neurons follow a shared developmental trajectory while maintaining distinct transcriptional and *cis*-regulatory programs. These differences in the chromatin landscape may enable common developmental TFs to drive subtype-specific gene expression^69^. Among the sequence features identified by deep learning, the positive AP-1-like contribution in the St18-positive/deep CA3 domain at late developmental stages is particularly interesting, given the intrinsically bursting phenotype of St18-positive CA3 PNs. AP-1 complexes include activity-regulated FOS and JUN family factors, and closely spaced bursts can elicit stronger *c-fos* transcription than equivalent activity delivered at longer intervals^70^. The AP-1-like signal may therefore reflect increased activity-dependent regulation as deep CA3 neurons mature. Whether AP-1 contributes to the development of these properties or is instead a consequence of increasing neuronal activity remains unclear.

The negative contribution of an ST18-like motif in St18-positive/deep CA3 neurons and the predicted repressive role of *St18* are consistent with previous studies describing ST18/MYT3 as a transcriptional repressor in other cellular contexts^25,71^. ST18 shares with other MyT family members a conserved SIN3-interaction domain that, in MYT1 and MYT1L, mediates recruitment of SIN3/HDAC corepressor complexes, suggesting that ST18 may repress transcription through a similar mechanism^72,73^. Whether ST18 engages a similar chromatin-based mechanism in CA3 neurons remains unknown.

Postnatal loss of ST18 shifted deep CA3 PNs toward an *St18*-negative molecular state. Deep CA3-associated genes were downregulated following *St18* deletion, including the expression of *Unc5d*, *Gabrg3*, *Cdh13*, and *Sema5a*. Conversely, superficial CA3-associated genes were upregulated, including *Tafa1* and *Kcnq5*, together with genes with known roles in MF-CA3 synapse organization and function such as *Nectin3* and *Neto1*, encoding a postsynaptic adhesion molecule and kainate receptor auxiliary subunit^47,57^, respectively. Chromatin accessibility shifted in parallel: regions that gained accessibility following ST18 loss were preferentially accessible in superficial CA3, whereas regions with reduced accessibility were associated with deep CA3.

The shift in deep CA3 PN molecular identity was accompanied by a change in cellular properties. ST18 loss increased TE volume and dendritic coverage in deep CA3 neurons toward superficial levels, indicating that ST18 normally constrains TE elaboration in deep CA3 PNs. Upregulation of *Nectin3* and *Neto1* raises the possibility that ST18 loss affects not only TE morphology but also MF synaptic organization and function. The accompanying increase in sEPSC frequency in *St18*-KO deep CA3 PNs supports enhanced synaptic input, although it remains unresolved whether this reflects MF-associated synapses or other inputs. ST18 loss may also affect output synapses of deep CA3 PNs through upregulation of its predicted target *Tafa1*, which encodes a secreted protein that associates with and regulates the post-translational modification of neurexins^29^, presynaptic adhesion molecules that are key regulators of synapse formation and synaptic properties^74,75^. Whether ST18 more broadly regulates the organization of deep CA3 connectivity will be important to investigate.

ST18 loss also reduced the characteristic burst-firing properties of deep CA3 PNs, while leaving most other measured electrophysiological properties, including voltage- gated Na⁺ and K⁺ current densities, largely unchanged. Several predicted ST18 targets involved in ion-channel function or regulation were upregulated following *St18* KO, raising the possibility that changes in intrinsic firing properties reflect the combined effects of multiple downstream genes. Among these, *Kcnq5*, which encodes the Kv7.5 potassium channel, is particularly interesting given the established role of Kv7 channels in regulating afterdepolarisation and burst firing in CA3 PNs^76^. Kcnq5 has also been shown to contribute to afterhyperpolarisation currents in these neurons^77^. Whether *Kcnq5* or other predicted ST18 targets contribute to the altered firing phenotype remains to be determined.

The preservation of most electrophysiological properties alongside changes in burst-firing and synaptic activity in the *St18*-KO suggests that postnatal ST18 loss induces a selective shift in subtype-associated features rather than a complete conversion of neuronal identity. The *St18* KO phenotype may arise from coordinated changes across multiple downstream effectors, as described for other TFs controlling connectivity. Acj6 regulates dendritic targeting in *Drosophila* olfactory projection neurons through combinations of cell-surface molecules^78^, whereas UNC-4 maintains VA motor-neuron input specificity in *C. elegans* by repressing a transcriptional program associated with VB neurons. Loss of UNC-4 shifts connectivity toward the alternative subtype without fully erasing VA identity^17,79,80^. A similar regulatory principle may apply to ST18, whose predicted targets include genes involved in dendrite development, synaptic organization, and electrophysiological properties. The acquisition of superficial-associated features following postnatal ST18 loss suggests that deep-superficial CA3 differences depend not only on initial specification, but also on continued repression of alternative subtype programs during circuit maturation.

In conclusion, our results identify ST18 as a regulator of deep CA3 identity that restricts a superficial-like molecular and postsynaptic program. These findings highlight how transcriptional regulatory programs contribute not only to neuronal molecular diversity but also to the structural and functional properties underlying their integration into neuronal circuits. They also raise the broader question of whether common regulatory principles govern neuronal diversity and circuit integration along other spatial axes of CA3 and across different hippocampal cell types. The postnatal hippocampal snMO atlas and the *St18*-Cre line provide complementary resources to investigate these questions and identify the downstream mechanisms linking subtype-specific regulatory programs to neuronal connectivity and function. Beyond understanding how these programs operate within individual neuronal populations, an important future challenge will be to determine how they are coordinated across synaptically connected neurons to shape hippocampal circuit organization and function.

## Methods

### Animals

All animal procedures were approved by the KU Leuven Ethical Committee for Animal Experimentation (protocol 006-2024) or by the Institutional Animal Care and Use Committee at the Janelia Research Campus (protocol 14-118). Mice were housed in a specific pathogen-free (SPF) facility under standard laboratory conditions, with ad libitum access to food and water. Animals were maintained on a 14h light/10h dark cycle at KU Leuven or a 12h light/12h dark cycle at Janelia Research Campus. WT C57BL/6J mice originated from Charles River. *St18*-Cre C57Bl/6J was generated by the Janelia Genomics core, as described below. H11- Cas9 C57Bl/6J (JAX:028239) was obtained from The Jackson Laboratory.

### Nucleus isolation and library preparation for 10x Genomics snMO

Nucleus preparation from mouse hippocampal tissue was done using the sucrose gradient protocol. Briefly, up to 50 mg of hippocampal tissue was homogenized using 1 ml of homogenization lysis buffer [1× salt-tris buffer: 146 mM NaCl, 10 mM tris (pH 7.5), 1 mM CaCl2, and 21 mM MgCl2; 0.03% (v/v) Tween-20, Sigma-Aldrich; 0.01% (v/v) NP-40, Sigma-Aldrich; 1% bovine serum albumin (BSA), Thermo Fisher Scientific; 25 mM KCl, Thermo Fisher Scientific; 250 mM sucrose, Sigma-Aldrich; 1 mM DTT, Thermo Fisher Scientific; 1× protease inhibitor, Sigma-Aldrich; RNase inhibitor, Promega]. The tissue was homogenized with 10× strokes with pestle A and 10× strokes with pestle B. The homogenized mixture was incubated in homogenization lysis buffer for 5 min. During the incubation, the homogenized sample was passed through 70-μm nylon mesh. The nuclei were pelleted by centrifugation at 500g for 5 min. The nucleus pellet was washed and resuspended with 520 μl of Wash Buffer 1 [1× salt-tris buffer; 1% BSA, Thermo Fisher Scientific; 25 mM KCl, Thermo Fisher Scientific; 250 mM sucrose, Sigma-Aldrich; 1 mM DTT, Thermo Fisher Scientific; 1× protease inhibitor, Sigma-Aldrich; RNase inhibitor (1 U/μl), Promega]. The nucleus suspension was then mixed with an equal volume of gradient medium [1 mM CaCl2, Thermo Fisher Scientific; 50% (v/v) Optiprep, Sigma-Aldrich; 5 mM MgCl2, Thermo Fisher Scientific; 10 mM tris (pH 7.5); Thermo Fisher Scientific; 75 mM sucrose, Sigma-Aldrich; 1 mM DTT, Thermo Fisher Scientific; 1× protease inhibitor, Sigma-Aldrich; RNase inhibitor (0.5 U/μl), Promega]. This was thoroughly mixed, and the mix was layered upon 948 μl of 29% density cushion [29% (v/v) Optiprep, Sigma-Aldrich; 72.5 mM KCl, Thermo Fisher Scientific; 14.5 mM MgCl2, Thermo Fisher Scientific; 29 mM tris (pH 8), Thermo Fisher Scientific; 121 mM sucrose, Sigma-Aldrich; RNase inhibitor (0.25 U/μl), Promega]. The layered nucleus suspension was subjected to centrifugation at 9000g for 20 min at 4°C. After the centrifugation, the debris was removed from the top of the suspension, and if excess debris was present, this could be scooped out with a 1000-μl pipette tip. The supernatant was aspirated out gradually using a glass Pasteur pipette until about 75 μl of the supernatant was left. An equal volume of 0.2× lysis buffer [20 mM tris (pH 7.5), Thermo Fisher Scientific; 20 mM NaCl, Thermo Fisher Scientific; 6 mM MgCl2, Thermo Fisher Scientific; 0.02% Tween 20, Sigma-Aldrich; 0.02% NP-40, Sigma-Aldrich; 0.002% digitonin, Promega; 2% BSA, Thermo Fisher Scientific; 2 mM DTT, Thermo Fisher Scientific; protector RNase inhibitor (2 U/μl), Sigma-Aldrich] was added, and the nuclei were resuspended and incubated for 2 min on wet ice. After the incubation, the lysis was quenched by adding 1 ml of Wash Buffer 2 [10 mM tris (pH 7.5), Thermo Fisher Scientific; 10 mM NaCl, Thermo Fisher Scientific; 3 mM MgCl2, Thermo Fisher Scientific; 0.01% Tween 20, Sigma-Aldrich; 1% BSA, Thermo Fisher Scientific; 1 mM DTT, Thermo Fisher Scientific; protector RNase inhibitor (1 U/μl), Sigma-Aldrich], and the nucleus suspension was resuspended. The nuclei were pelleted by centrifugation at 500g for 5 min. The supernatant was discarded until about 20 μl of the supernatant was left in the Eppendorf. Removal was done carefully without disturbing the nucleus pellet. The nucleus pellet was resuspended with 20 μl of 2× Nuclei Buffer [2× Nuclei Buffer, 10x Genomics, PN-2000153; 1 mM DTT, Thermo Fisher Scientific; protector RNase inhibitor (1 U/μl), Sigma-Aldrich].

The nucleus count and the viability of the samples were accessed using a LUNA dual florescence cell counter (Logos Biosystems), and a targeted cell recovery of 10,000 cells was aimed for each of the samples. Library preparations for 10x Genomics single-cell multiome were performed using 10x Genomics Chromium NextGEM Single cell Multiome ATAC+Gene Expression chemistry (10x Genomics, Pleasanton, CA). Transposition of nuclei and the rest of the 10x Genomics Multiome library preparation were performed as per the protocol recommendations (Chromium Next GEM Single Cell Multiome ATAC + Gene Expression Reagent Kits User Guide; CG000338 Rev. G). At the different checkpoints, the library quality was accessed using Qubit (Thermo Fisher Scientific) and Bioanalyzer (Agilent).

10x libraries were sequenced either on an Illumina platform or on an MGISEQ-2000 sequencing platform (MGI Hong Kong SAR sequencing facility). Before being able to sequence on the MGI platforms, the 10x libraries need to undergo a conversion step using the MGIEasy Universal DNA Library Prep kit. Briefly, the final 10x sequencing libraries were circularized using the splint-ligation step, and the circularized libraries were converted to single-stranded DNA copies. DNA nanoballs were prepared from the circularized single-stranded DNA using rolling circle amplification. DNA libraries generated were then flown through the patterned flow cell of the MGISEQ-2000RS High-Throughput Sequencing kit. For a targeted sequencing saturation of 50 to 60%, sequencing was performed at a depth of 30,000 to 60,000 reads per cell for the RNA and ATAC libraries. For RNA libraries, the sequencing recipe of paired-end reads of 28 base pairs (bp) (read 1), 100 bp (read 2), 10 bp (index 1), and 10 bp (index 2) was used for sequencing. For ATAC libraries, the sequencing recipe of paired-end reads of 50 bp (read 1), 49 bp (read 2), 24 bp (index i5), and 8 bp (index i7) was used for sequencing.

### Single molecule Fluorescent *In* Situ Hybridization RNAscope

Mice were euthanized at P5, P15, and P28, and brains were rapidly dissected, embedded in O.C.T. compound (4583, Sakura Finetek), frozen, and cryosectioned coronally at 10 μm using a CM3050 S cryostat (Leica). Sections were mounted on SuperFrost Ultra Plus adhesion slides (Thermo Fisher Scientific), air-dried at room temperature, and processed using the RNAscope™ Multiplex Fluorescent Detection Kit v2 (ACD Bio, 323110) according to the manufacturer’s instructions. Wild type (WT) C57BL/6J mice were used. Brain sections were mildly fixed through a 50–70–100% ethanol series, followed by 10 min hydrogen peroxide treatment and 10 min Protease IV treatment, with 1× PBS washes between steps. *St18* (Bio- Techne, 443271), *Tafa1* (Bio-Techne, 520731), and *Vgat* (Bio-Techne, 319191) probes were hybridized at 40°C for 2 h, followed by HRP-mediated signal development using OPAL dyes, DAPI counterstaining, and mounted with Mowiol. Images were acquired with a confocal microscope Zeiss Cell Discoverer 7 using a 20X objective.

### Cell lines

HEK293T-17 human embryonic kidney cells (hereafter referred to as HEK cells) were obtained from American Type Culture Collection (ATCC) cat# CRL-11268, and were used for AAV production. HEK293T-17 cells were grown in Dulbecco’s modified Eagle’s medium (DMEM; Thermo Fisher Scientific) supplemented with 10% fetal bovine serum (FBS; Thermo Fisher) and penicillin/streptomycin (Thermo Fisher).

### Plasmids

Plasmids were either obtained (pAAV-TRE-DIO-FLPo, Addgene #118027; pAAV-TRE-fDIO-GFP-ires-tTA, Addgene 118026; pAAV-CAG-GFP, Addgene 11150) or generated in-house (pAAV-TRE-FLPo; AAV-hSyn-NLS-tdTomato; pAAV-hU6-gRNA-TRE-fDIO-EGFP-IRES-tTA) using Gibson assembly and/or Golden Gate assembly.

### *St18*-cre mouse strain

This mouse strain was generated in the laboratory of N. Spruston (HHMI Janelia Research Campus). The St18 T2A frt-Neo-frt Cre construct DNA and Cas9/gRNA were electroporated into embryonic stem (ES) cells derived from F1 embryos (129S6 × C57BL/6J). Following five days of G418 selection, ES cell clones were picked and screened for correct targeting. Positive clones were expanded and aggregated with CD1 embryos, then cultured overnight at 37°C, 5% CO2 incubator. The resulting blastocysts were transferred into the uteri of recipient mice to generate chimeras. Chimeras were confirmed by PCR and subsequently bred with R26FLP mice (JAX#016226) to remove the Neo selection cassette. The offspring that were positive for St18–T2A–Cre and negative for FLP were used for further experiments.

### AAV purification

High-titer AAV vectors were produced and purified as previously described^81^. Briefly, six 70% confluent HEK cells culture plates were co-transfected using polyethyleneimine (PEI) with 20 μg of plasmid, 10 μg of pRepCap 2/9 serotype plasmid (Addgene 112865), and 20 μg of the AAV genome plasmid (Addgene, 112867) per plate. Three days after transfection, cells were harvested by scraping and centrifugation. AAV particles present in the supernatant were precipitated using polyethylene glycol (PEG) and combined with the corresponding cell lysates. Lysates were subsequently treated with Benzonase nuclease to degrade contaminating cellular DNA. Purification of AAV particles was performed by ultracentrifugation on an iodixanol density gradient (15%, 25%, 40%, and 60%) at 50,000 rpm for 1h40 min at 12 °C. The purified viral particles were collected from the 40% iodixanol fraction, followed by desalting and concentration using a 4 mL Amicon centrifugal filter (Sigma).

Low-titer AAVs were produced as previously described^53^. Briefly, HEK cells were plated in 12-well cell-culture and upon reaching ∼80% confluency were transfected with a DNA::PEI complex consisting of 800 ng pDelta F6 plasmid, 400 ng RepCap 2/1 plasmid and 400 ng AAV genome plasmid, as well as 5.7 μl PEI, all in a total volume of 100 μl Optimem (Thermo Fisher). 3 days later, the supernatant was collected, filtered through a 0.45 um filter and concentrated by centrifugation on 100 kDa MWCO 0.5 ml Amicon columns (Sigma) to a volume of approximately 30 μl. The resulting AAVs were used at a dilution of 1:500 to 1:2000 to transduce primary neuronal cultures.

### Stereotactic brain injections

Neonatal injections at P0: neonates were anesthetized by inducing hypothermia for 15min on ice prior to injection and maintained during injection by placing the pup on an ice-cooled glass petri dish. Injections are executed in a Kopf stereotactic frame mounted with a Drummond Nanoject III containing a beveled glass capillary filled with a viral construct. Pups are manually immobilized. The capillary is inserted into the brain at CA3 coordinates (mediolateral 1.5; dorsoventral 1.6; anteroposterior two spots visually determined between bregma and lambda) and 50nl viral construct mix is injected bilaterally in each spot at 10nl/s. Mice are recovered on a 37°C heating plate before being placed back in the nest.

Adolescent injections: Mice are anesthetized with 5%isoflurane and immobilized in a Kopf stereotactic frame mounted with a Drummond Nanoject III containing a glass capillary filled with a viral construct. Body temperature is controlled by a temperature-controlled heating plate. Anesthesia during the surgery is maintained with 2-3% isoflurane. Duratears are applied to the eyes. General analgesia (Buprenorphine 0,05mg/kg) is subcutaneously administered. The incision site on the head is shaved, disinfected with iso-betadine, and local analgesia (lidocaine 6mg/kg) is subcutaneously administered. After a skin incision on the head, the skull is exposed, and unilateral/bilateral coordinates for CA3 (mediolateral 2.5, anteroposterior 1.9, dorsoventral 1.8) are determined from bregma. A small burr hole in the skull is made and the capillary is inserted in the brain to inject a viral construct mix (100-300nl). The capillary is retracted and skin incision is closed with surgical glue (Dermafuse). Mice are recovered on a 37°C heating plate. In general, a two weeks transduction period of the virus is respected before proceeding with tissue collection.

### CRISPR KO validation in neuronal cultures

Neuronal cultures were obtained from dissected E18 H11-Cas9 mice or C57Bl6J WT embryos and plated on poly-d-lysine (Millipore)– and laminin (Invitrogen)–coated six-well plates. Neurons were maintained in Neurobasal medium (Invitrogen) supplemented with B27. Forty-eight to 72 hours after seeding, 10 mM glia inhibitor (5-fluoro-2-deoxyuridine; Sigma-Aldrich) was added to the medium. Two *St18* gRNAs were designed to induce a KO using CRISPick^82^ for *St18* CRISPR-mediated KO. To assess KO efficiency, we transduced Cas9-expressing neuronal cultures with low titer AAVs expressing either gRNA, as well as an AAV expressing an HA-tagged St18 construct under the control of a human Synapsin promoter. Cell Lysis followed by Western blot analysis using an anti-HA antibody, revealed a clear reduction in ST18 expression for one of the gRNA constructs, which was subsequently selected for further in vivo validation using immunohistochemistry.

### Western blot

For Western blot, samples consisted of primary neuronal cultures scraped in RIPA buffer (50 mM Tris pH 8, 150 mM NaCl, 1% SDS, 0.5% Sodium Deoxycholate, 1% Triton-X 100, 1X protease inhibitors) or of CA3 lysates (see below). For either, protein concentration was determined using BCA, and an appropriate amount of lysate corresponding to 5 to 10 ug of total protein was boiled 10 mins at 95 C in Laemmli buffer. Then, samples were loaded on 10% or 7.5% polyacrylamide gels and run at 180 V. Proteins were transferred to 0.2 um nitrocellulose membrane using semi-dry transfer (Biorad Transblot Turbo, mixed-molecular weight program). Immediately after transfer, a total protein stain (LI-COR Revert Total Protein Stain) was used to normalize loading. Membranes were then blocked in 10% milk in TBS-T buffer (25 mM Tris-base pH 7.5, 300 mM NaCl, 0.05% Tween-20) for 30 mins at RT. Primary and secondary antibodies were diluted in 5% milk in TBS-T. Primary antibodies were incubated O/N at 4 °C, secondary antibodies were incubated 2 to 3 h at RT. When probing for biotinylated proteins, membranes were thoroughly washed in TBS-T and HRP-conjugated streptavidin was diluted 1:500 in 5% BSA in TBS-T to avoid binding to the biotin present in the milk. After primary and secondary antibody incubation, membranes were washed 5 times with TBS-T and once with 1X PBS before development at ImageQuant 800 (Cytiva) using SuperSignal West Pico PLUS and Femto Chemiluminescent Substrate (Thermo Fisher 34577 and 34094).

### Immunohistochemistry

Mice were transcardially perfused under lethal dose anesthesia (ketamine/xylaine/saline) with 25 ml of 4%PFA 2% sucrose in 1X PB buffer pH 7.4. Brains were dissected, post-fixed in perfusion solution for 1h, washed in phosphate buffer (PB), and sectioned into 80 μm-thick slices using a vibratome (Campden instruments 7000smz). Immostaining is conducted by blocking in 10% normal horse serum, 0.5M glycin, 0.5%Triton-X-100 in 1X PBS, followed by antibody incubation (anti-ST18^83^ 1:300; anti-RFP 1:500, Proteintech Papbr1-20; anti-GFP 1:1000, Aves 10-10) in antibody solution for 48h at 4°C, repeated washes in 1X PBS, secondary antibody solution for 24h at 4°C, repeated washes in 1X PBS and nuclear staining with DAPI (1:1000; Fisher Scientific 10184322). Primary and secondary antibodies are diluted in 5% normal horse serum, 0.5%Triton-X-100 in 1X PBS. Sections are then mounted on a microscope slide with Mowiol and a coverslip.

Adjustments to the protocol were made for the morphological analysis of CA3 TEs. Perfused brains were sectioned into 200µm thick vibratome slices and cleared according to the manufacturer’s instructions (Life Canvas Technology). In brief, sections are incubated in Shield fixative for 48h at 4°C, Shield-ON+epoxy 24h at 37°C, Shield-ON 24h at 37°C and delipidated in Delipidation buffer for 48h at 42°C prior to immunostaining. Sections are mounted on a microscope slide with a 250µm spacer in Easyindex (RT=1.52). Images were acquired on a confocal microscope Zeiss Cell Discoverer 7 or Zeiss LSM900 Airyscan.

### snMO *St18* KO sequencing

Neonatal offspring from *St18-Cre × H11-Cas9* crosses received stereotactic brain injections as described above. Littermates were injected with either the control or *St18* KO viral mixture. Two weeks after viral transduction (P14–P16), animals were anesthetized with isoflurane and brains were collected for dissection. Hippocampi were dissected in ice-cold HBSS. Successful viral transduction of CA3 was scored by hippocampal GFP fluorescence prior to tissue freezing. Following dissection, hippocampi were snap-frozen in liquid nitrogen and stored at −80 °C until further processing.

### Nuclei isolation

Frozen mouse hippocampal tissue was processed for nuclei isolation without prior thawing. Hippocampal tissue from 1 male and 1 female mice was pooled by experimental condition (WT or KO) prior to nuclei isolation, with approximately 40–80 mg of tissue processed per pooled sample. Frozen tissue was transferred directly from dry ice into a pre- cooled Dounce homogenizer containing 750 μL of homogenization/lysis buffer (HLB; containing 1x ST-buffer (10 mM Tris pH 7.4, 146 mM NaCl, 1 mM CaCl2, 21 mM MgCl2), 1% BSA, 25 mM KCl, 250 mM sucrose, 0.03% Tween-20, 0.03% Igepal-CA630, 1X cOmplete protease inhibitor (Roche), 1 mM DTT, and RNase inhibitor (Promega)). Tissue was allowed to thaw in the buffer for 5 min before homogenization. Mechanical dissociation was performed using 10 strokes with pestle A, followed by a 1-min incubation and 10 strokes with pestle B. The homogenate was passed through a 70-μm cell strainer, and the homogenizer and strainer were rinsed with an additional 250 μL of homogenization/lysis buffer. Following a 5-min incubation on ice, nuclei were collected by centrifugation at 500 × g for 5 min at 4°C using a swinging-bucket rotor. The nuclei pellet was gently resuspended in 520 μl nuclei wash buffer 1 (WB1; 1x ST-buffer, 1% BSA, 25 mM KCl, 250 mM sucrose, 1X cOmplete protease inhibitor, 1 mM DTT, and RNase inhibitor) before density gradient purification. 520 μl gradient medium (GM: 75 mM sucrose, 10 mM Tris pH 7.4, 1 mM CaCl2, 5 mM MgCl2, 50% Optiprep, 0.5X complete protease inhibitor, 1 mM DTT, and RNase inhibitor) was added to the sample. For further purification, the nuclei suspension was carefully layered onto 770 μL of 29% density cushion (29% Optiprep, 75 mM KCl, 15 mM MgCl2, 30 mM Tris pH 8, and 125 mM sucrose) and centrifuged at 9,000 × g for 30 min at 4°C in a swinging-bucket rotor without brake. Following centrifugation, the supernatant was carefully removed. The purified nuclei were permeabilized by gently resuspending the pellet in lysis buffer (20 mM Tris pH 7.4, 20 mM NaCl, 6 mM MgCl2, 0.02% Tween-20, 0.02% Igelpal-CA630, 0.004% Digitonin, 1% BSA, 1 mM DTT, RNase inhibitor) and incubating on ice for 2 min. After incubation, 1000 μL chilled wash buffer 2 (WB2; 10 mM Tris pH 7.4, 10 mM NaCl, 3 mM MgCl2, 1% BSA, 1 mM DTT, RNase inhibitor) was added to the nuclei. The nuclei were spun down at 450g, 4°C for 5 min. The obtained nuclei pellet, after supernatant removal, was resuspended in diluted nuclei buffer (1x nuclei buffer (10x Genomics, Cat. No 2000207), 1 mM DTT, RNase inhibitor). Nuclei yield, morphology and purity was visualized using a LUNA-FL dual Automated cell counter (Logos biosystems) before downstream processing.

### Single-nucleus Multiome ATAC and gene-expression library preparation

Purified nuclei were processed using the 10x Genomics Chromium GEM-X Single Cell Multiome ATAC + Gene Expression workflow (10x Genomics, Pleasanton, CA). For each experimental condition, 75,000 nuclei were used as input for transposition, with a targeted nuclei recovery of ≥20,000 nuclei per sample. Transposition was performed for 30 min, after which tagmented nuclei were processed for simultaneous capture of chromatin accessibility and gene-expression information according to the manufacturer’s GEM-X Multiome workflow. Library concentration and fragment-size distribution were assessed using Qubit (Thermo Fisher Scientific) and Agilent Fragment Analyzer.

### Sequencing and processing of raw FASTQ files

Libraries were submitted to VIB Nucleomics Core (Leuven, Belgium) for sequencing on Element AVITI and AVITI24 platforms using the paired-end Cloudbreak Freestyle 2×75 High kit. ATAC and GEX libraries were sequenced separately to accommodate their different read length requirements. ATAC libraries were sequenced with a 51-8-24-50 cycle configuration (Read 1–i7 index–i5 index–Read 2), and GEX libraries with a 29-10-10-91 cycle configuration (Read 1–i7 index–i5 index–Read 2). Raw sequencing data were demultiplexed using the Bases2Fastq software version.

### Morphological analysis

#### CA3 cell spatial distribution

The spatial localization of CA3 PNs was determined from overview images using FIJI software. The boundary between the deep and superficial layers was defined by dividing the CA3 *stratum pyramidale* (SP) in two equal segments longitudinal to the SP. To distinguish the CA3 subfields (CA3a, CA3b, and CA3c), the total linear length of the CA3 SP layer was divided into three equal segments. The position of each neuron was normalized to both the width and the total length of the SP to account for dorsoventral differences in SP morphology. *St18*-expression was determined by tdTomato-expression.

#### TE morphology

High-resolution images of the entire dendritic tree of PNs within the CA3 *stratum lucidum* were acquired by tile-scanning Z-stacks. The spatial distribution of TEs was determined as described above. Dendritic shafts were manually removed in FIJI to generate a segmentation mask containing only TEs. The segmentation mask was subsequently imported into Imaris (Bitplane) together with the original image to generate an overlay. Using this overlay, TEs were segmented as a new channel, enabling three-dimensional reconstruction and quantitative analysis. Morphological parameters included total TE volume, dendritic TE coverage, and the distance from the apex of the pyramidal soma to the first TE cluster.

### In utero viral injection

Timed-pregnant C57BL/6NCrl damswere used, with embryonic age set by vaginal plug (E14 or E17). Dams were anesthetized with 2 to 2.5% isoflurane and received buprenorphine (0.1mg/kg SC), ketoprofen (5mg/kg) analgesia during surgery. The abdomen was shaved, cleaned, and disinfected, and the uterine horns were exposed and removed from the abdominal cavity. Embryos were injected bilaterally into the lateral ventricles with 140nL per hemisphere of adeno-associated virus (1/50 diluted AAV1-CAG-GFP; Addgene ref 37825), using a Nanoject nanoliter injector (Drummond Scientific). The uterus was returned to the abdomen, the abdominal wall was sutured, the skin was stitched, the dam was allowed to recover and received postoperative analgesia once per day for 2 days. Brains were collected at P30-40 for further tissue processing. Number of animals: E14 n = 7; E17 n = 7.

### Cell sorting and transcriptional profiling

Manual cell sorting was performed as previously described (Hempel et al., 2007; Sugino et al., 2014). Animals that received in utero viral injection (age at collection: P30-P40) were rapidly decapitated following isoflurane anesthesia, and 200 µm slices were digested with Pronase E (1 mg/mL, P5147; Sigma-Aldrich) for 1 hour at room temperature in artificial cerebrospinal fluid (ACSF) containing 6,7-dinitroquinoxaline-2,3-dione (20 µM; Sigma-Aldrich), D-(-)-2-amino-5-phosphonovaleric acid (50 µM; Sigma-Aldrich), and tetrodotoxin (0.1 µM; Alomone Labs). Hippocampal CA3 was micro-dissected and triturated with Pasteur pipettes of decreasing tip size. Cell suspensions were diluted 5 to 20 fold with filtered ACSF containing fetal bovine serum (1%; HyClone) and poured over Petri dishes coated with Sylgard (Dow Corning). For dim cells, glass-bottom Petri dishes were used. Fluorescent cells were aspirated into a micropipette (tip diameter 30 to 50 µm) under a fluorescence stereomicroscope (M165FC; Leica) and washed three times by transfer to clean dishes. After the final wash, pure samples were aspirated in 1 to 3 µL and lysed in 47 µL XB lysis buffer (PicoPure kit, KIT0204; Thermo Fisher) in 200 µL PCR tubes (Axygen), incubated for 30 minutes at 40 °C on a thermal cycler, and stored at −80 °C.

Total RNA was extracted with the PicoPure kit. Either 1 µL of a 10^-5 dilution of ERCC spike-in control (#4456740; Life Technologies) or (number of sorted cells/50) x 1 µL of that dilution was added to the purified RNA, which was concentrated by speed-vacuum to 5 µL and immediately reverse transcribed using the NuGEN Ovation RNA-Seq System V2 (#7102; NuGEN), yielding 4 to 8 µg of amplified DNA. Amplified DNA was fragmented (Covaris E220) to approximately 200 bp and ligated to Illumina sequencing adaptors with the Encore Rapid kit (0314; NuGEN). Libraries were quantified with the KAPA Library Quant Kit (KAPA Biosystems) and sequenced on an Illumina HiSeq 2500 with 4 to 32-fold multiplexing (single end, usually 100 bp read length).

#### Proteomics and proximity labelling

#### Proximity labeling

P28 *St18-*Cre mice were injected stereotactically injected in CA3 (both hemispheres) with the Supernova AAV setup described earlier, combined with either of the following AAVs: pAAV-CamkII-Ctrl-TurboID, pAAV-CamkII-PSD95-TurboID, pAAV-CamkII-DIO-Ctrl-TurboID, pAAV-CamkII-DIO-PSD95-TurboID (4 replicates per AAV/condition, 3 to 4 animals per replicate, a mix of males and females). Seven days after AAV injection, mice were injected subcutaneously with 1 ml of 2 mg/mL biotin in 1X PBS, adjusting the pH to 7.4 to increase biotin solubility. 3 to 5 h after injection, mice were briefly anaesthetized by isoflurane and brains collected in ice cold HBSS + 10 mM HEPES. 500 μm coronal brain sections were produced using an ice-cold brain matrix (AgnThos 69-2165-1), and CA3 was dissected under a fluorescence stereomicroscope using fluorescent signal expressed by infected cells and morphological landmarks. Tissue pieces from each mouse were collected in a separate tube, flash-frozen in liquid nitrogen after dissection buffer removal, and stored at −80 °C until needed. Protein extraction and pulldown was performed as previously described^54^. Briefly, tissue from one mouse was homogenized and lysed in 50 μl 1% SDS RIPA buffer (50 mM Tris pH 8, 150 mM NaCl, 1% SDS, 0.5% Sodium Deoxycholate, 1% Triton-X 100, 1X protease inhibitors) by pipetting and trituration. Lysates were sonicated in a bath sonicator (124-9721 RS-online) at max power three times for 1 min with 30 s rest on ice in between. Lysates were then boiled at 95 °C for 5 min and immediately diluted to 0.5% SDS with SDS-free RIPA buffer. After 3 h incubation at 4 °C with rotation, lysates were cleared by centrifugation at 20.000 x g for 15 min at 4 °C and supernatant collection. Bicinchoninic acid assay (BCA) was used to quantify protein concentration, and streptavidin-HRP dot blots using 1 ul of lysates were performed to assess biotinylation levels. Between 300 μg and 500 μg protein input was used for pulldown for MS and the amount was adjusted to roughly normalize biotinylation levels across replicates. 10 μl of streptavidin magnetic beads (Pierce) per 100 μg of protein input was used for every condition. Beads were washed 3x in 0.5% SDS RIPA buffer using a magnetic rack and incubated overnight at 4 °C with rotation with protein input in 1 ml of 0.5% SDS RIPA buffer final volume. The following day, beads were washed 3x with 0.5% SDS RIPA buffer, 1x with 1 M KCl, 1x with 0.1 M Na2CO3, 3x with 2 M Urea in 50 mM Tris-HCl pH 8.

#### MS sample preparation

Proteins were eluted from the beads and digested using an S-Trap micro kit (ProtiFi) according to the manufacturer’s instructions, with the modifications described below. Briefly, beads were resuspended in 92 µL of 5% SDS in 50 mM triethylammonium bicarbonate (TEAB; pH 8), heated at 95 °C for 5 min, and further cooled down at room temperature. Proteins were reduced by adding 4 µL of 120 mM tris(2-carboxyethyl)phosphine (TCEP) and incubating at 55 °C for 20 min. Cysteines were alkylated by adding 4 µL of 500 mM methyl methanethiosulfonate (MMTS) in isopropanol and incubating at room temperature for 20 min in the dark. Samples were acidified with 10 µL of 27.5% phosphoric acid in distilled water. Beads were separated using a magnetic rack, and supernatants were transferred to new low binding tubes.

Following the addition of 660 µL of binding buffer (100 mM TEAB in 90% methanol), proteins were loaded onto S-Trap columns and centrifuged at 4,000 × g for 1 min. Columns were washed six times with 150 µL of binding buffer. As a next step, proteins were digested by adding 1 µg of trypsin in 20 µL of 50 mM TEAB directly to each column and incubating overnight at 37 °C in a humidified oven. Next day, peptides were sequentially eluted by centrifuging at 4,000 × g for 1 min 1. 40 µL each of 50 mM TEAB, 2. 0.2% formic acid in water, and 3. 0.2% formic acid in 50% acetonitrile. Peptide eluates were dried using a SpeedVac before LC–MS/MS analysis.

#### Mass spectrometry

Peptide samples were analysed using an Evosep ENO liquid chromatography system (Evosep Biosystems) coupled online to an Orbitrap Astral Zoom mass spectrometer (Thermo Fisher Scientific) equipped with an EASY-Spray ion source. Peptides were resuspended in 20 µL of solvent A (0.1% formic acid in water), and 1 µL of each sample was loaded onto an Evotip according to the manufacturer’s instructions. Peptides were separated on an Aurora Elite column (15 cm × 75 µm; IonOpticks), maintained at 50°C, using the Whisper Zoom 40 samples-per-day method. Full MS spectra were acquired in the Orbitrap analyser at a resolution of 240,000 over m/z 380-980, with a normalised automatic gain control (AGC) target of 500% and a maximum injection time of 5 ms. Data-independent acquisition (DIA) MS/MS spectra were acquired in the Astral analyzer using 150 scan events spanning the precursor range m/z 380-980, with a nominal isolation width of 4 Th and window-placement optimization enabled. The MS/MS scan range was m/z 150-2000, the normalized AGC target was 100%, and the maximum injection time was 8 ms. Higher-energy collisional dissociation (HCD) was performed at a normalized collision energy of 25%. Pre-accumulation was enabled, and the loop-control time was set to 0.6 s.

#### MS data analysis

Raw MS data were analysed using DIA-NN version 2.6.0 in library-free mode. The UniProt Mus musculus reference proteome (UP000000589, downloaded in May 2026) and a cRAP contaminant database were used. Enzyme specificity was set to trypsin, allowing up to one missed cleavage, and N-terminal methionine excision was enabled. Peptide lengths of 7-30 amino acids and precursor charge states of 1-4 were specified. Methylthiolation of cysteine (UniMod:39; +45.987721 Da), corresponding to MMTS alkylation, was specified as a fixed modification using the option --fixed-mod UniMod:39,45.987721,C. Methionine oxidation and protein N-terminal acetylation were specified as variable modifications, with a maximum of one variable modification per peptide. MS1 and MS2 mass tolerances were fixed at 4 and 10 ppm, respectively. Match between runs and peptidoform scoring were enabled. Unless otherwise specified, default DIA-NN settings were used.

Downstream analyses in R used the DIA-NN unique-gene quantification matrix. Abundances were log2-transformed, and proteins quantified in all replicates of at least one condition were retained. Missing values were imputed using impSeq (rrcovNA, version 0.5.3) used otherwise. Principal component analysis was performed on the processed data. Differential abundance analysis was conducted using limma (version 3.62.2).

### Patch-clamp electrophysiology in acute slices

Electrophysiological recordings were performed at the Electrophysiology Expertise Unit of VIB-KU Leuven Center for Neuroscience. Neonatal offspring from St18-Cre or St18-Cre × H11-Cas9 received stereotactic brain injections at P0 as described above. Animals (*St18-*negative vs *St18-*positive; n=12 animals between P13-21), (*St18-*CTRL vs *St18-*KO; n= 6 for each condition between P13-19) were anesthetized with isoflurane. After decapitation, the brain was quickly removed and transferred into ice-cold cutting solution (87 NaCl, 2.5 KCl, 1.25 NaH2PO4, 10 glucose, 25 NaHCO3, 0.5 CaCl2, 7 MgCl2, 75 sucrose, 1 kynurenic acid, 5 ascorbic acid, 3 pyruvic acid, pH 7.4 with 5% CO2/95% O2). After brain dissection, whole brain sagittal slices (300 µm) were cut with a vibratome (Leica VT1200) using a 10% angle. Afterwards slices were transferred to 33°C cutting solution for 45 minutes to recover and finally maintained at room temperature until used for recordings. During recordings brain slices were continuously perfused in a submerged chamber (Warner Instruments) at a rate of 3-4 ml/min with (in mM): 127 NaCl, 2.5 KCl, 1.25 NaH2PO4, 25 NaHCO3, 1 MgCl2, 2 CaCl2, 25 glucose and 1 µM bicuculline at pH 7.4 with 5% CO2/ 95% O2. Whole-cell patch-clamp recordings were done using borosilicate glass recording pipettes (resistance 3.5–5 MΩ, Sutter P-1000), using a double EPC-10 amplifier under control of Patchmaster v2 x 32 software (HEKA Elektronik, Lambrecht/Pfalz, Germany) or a DigiData1550B combined with Multiclamp 700B using Clampex 10.7 (Axon Instruments). The following internal medium was used (in mM): 135 K-Gluconate, 4 KCl, 10 HEPES, 4 Mg-ATP, 0.3 Na2-GTP, 10 Na2-phospocreatine, 3 biocytin (pH 7.3). Cell intrinsic properties were recorded in current clamp, while Na^+^/K+ currents and sEPSCs were recorded in voltage clamp using Vhold = −70 mV. Currents were recorded at 20 Hz and low-pass filtered at 3 kHz when stored. Single action potentials were recorded at 50 Hz and low-pass filtered at 10 kHz. The access resistance and membrane capacitance were manually compensated and subsequently 70-75% series resistance compensation was used for voltage clamp experiments. Only cells with an access resistance less than 20 MΩ were included for analysis. Junction potential was calculated to be approximately 12 mV, data shown are not compensated. Cell intrinsic properties were extracted using Easy Electrophysiology (easyelectrophysiology.com). Spontaneous input was analysed using Mini Analysis program (Synaptosoft).

### Bioinformatics Analyses

#### Read mapping snMO read mapping

The generated raw FASTQ files were mapped to the *Mus musculus* (mm10) reference genome using the count pipeline of Cell Ranger ARC (10x Genomics), version (v) 2.0.2 for the developmental snMO dataset and v2.1.0 for the *St18* KO snMO dataset, with intronic reads included by default.

#### Gene expression data quality control

##### snMO

Gene expression data quality control was performed independently for each sample using the Scanpy^84^ tool v1.8.2. For each sample, genes expressed in at least 3 cells were retained, as well as cells expressing a minimum of 200 genes and with a mitochondrial DNA content below 5%. Doublets were then identified and removed using the Scrublet^85^ tool, either by directly removing identified doublets or by excluding clusters with a high mean doublet score.

#### Cell type annotations and merge samples snMO gene expression data

Cell type annotation was performed via label transfer from the Allen Institute’s high-resolution transcriptomic atlas (ABC atlas) of the adult mouse brain using the 10x v2/v3 hippocampal formation reference dataset^24^. Datasets were integrated using a canonical correlation analysis (CCA) based workflow using IntegrateLayers() function with Seurat^86^ v5.2.1. Replicates from P28 samples (n=2) and P15 samples (n=2) were merged using Seurat v5.3.0 merge() function. Labels were transferred from the Allen reference^24^ to the P28 dataset, with FindTransferAnchors() and MapQuery() in Seurat v5.3.0 and PCA as the reference reduction. We then applied a hierarchical, age-sequential label-propagation strategy, using the annotated P28 dataset as the reference for P15, the annotated P15 dataset as the reference for P10, and the annotated P10 dataset as the reference for P5. Label transfer was performed at the cluster level: each target cluster was assigned the transferred cluster identity, and all associated higher-level annotations. Samples were merged using Seurat merge() function resulting in a total of 123,093 cells. Following integration, nuclei were clustered using the CCA- integrated reduction (dimensions 1-30) with the FindNeighbors() and FindClusters() functions in Seurat, at a resolution of 2. Cluster annotations were manually assessed for consistency with the transferred cell-type identities and marker gene expression. One cluster containing nuclei assigned to multiple cell types, without a consistent subclass-level annotation, was classified as “unknown.” An additional cluster, predominantly annotated as ‘064 STR-PAL Chst9 Gaba’, was also reclassified as “unknown” due to the absence of expected cell-type-specific marker expression. Finally, 26 nuclei with more than 100,000 total RNA counts were excluded as outliers. Lastly, nuclei with a label-transfer confidence score below 0.2 were annotated as low confidence annotations. This resulted in 119,380 annotated high quality nuclei. Mitochondrial genes, uncharacterized/predicted genes, and a list of sex-related genes^22,87^ were then removed. Nuclei were further filtered on the basis of snATAC-seq quality (see below for further details), leading to 94,981 cells, with a median of 4,267 unique molecular identifiers and 2,048 expressed genes per nucleus.

Marker genes for each cell class containing at least 50 cells were identified using the Wilcoxon rank-sum test implemented in rank_genes_groups() from Scanpy v1.8.2, comparing each class against all remaining cells using log-normalized gene expression values. Tie correction was enabled, and p-values were adjusted for multiple testing using the Benjamini-Hochberg method. The three highest-ranked marker genes for each cell class were visualized using rank_genes_groups_dotplot(). Dot size represents the fraction of cells within each class expressing the gene, while color represents the mean expression level within the class.

#### Hippocampal temporal gene expression analysis

Hippocampal gene expression data was normalized using NormalizeData() and was subsequently scaled using ScaleData() from Seurat v5.3.0. Developmental stages were ordered chronologically as P5, P10, P15 and P28. Temporal differential expression analysis was performed independently for each annotated cell subclass using TDEseq^88^ v1.1. Subclasses containing fewer than 20 cells were excluded from the analysis. Within each subclass, the 2,000 most highly variable genes (HVGs) were identified and retained for temporal modelling using FindVariableFeatures() method with “vst” method. TDEseq was applied at the single-cell level using tdeseq() function with a linear mixed-effects model, with developmental age specified as the stage variable and sample batch identity specified as the sample-level grouping variable. The FastLMM implementation was used for model fitting. Genes were filtered using a minimum detection fraction of 0.1 and a minimum log2 fold change (LFC) threshold of 0.1, adjusted p-value threshold of 0.05, and developmental stages represented by fewer than three cells were excluded. GO enrichment analysis was performed for genes assigned to the Recession, Peak, and Growth temporal expression patterns within the CA3 glutamatergic neuronal subclass (017 CA3 Glut). Only 13 genes were assigned to the “trough” pattern in CA3. Given the small number of genes and the limited functional enrichment, this trajectory was not characterized further. GO enrichment was performed independently for each temporal expression pattern using the enrichGO function in clusterProfiler^89^ v4.14, restricted to the BP ontology. Enriched GO terms were identified using p-value and q-value cutoffs of 0.05. Up to six enriched GO terms were retained for each temporal expression pattern.

#### snMO snATAC-seq analysis

For hippocampal analysis, each sample was processed separately using pycisTopic v2.0a0 (python implementation of cisTopic^90^). Using cell type annotations derived from the gene expression analysis, pseudobulk accessibility profiles were generated per cell subclass and per age-subclass. Cells with low-confidence or unknown annotations and subclasses represented by fewer than 10 cells were excluded. Peaks were called on each pseudobulk profile using MACS2^91^ v2.2.9.1 with the following parameters: --format BEDPE, --gsize mm, --qvalue 0.01, --keep-dup all, --shift 73 and --ext_size 146. To derive consensus peaks, an iterative peak filtering approach^92^ implemented in pycisTopic was applied using get_consensus_peaks() function with peak_half_width of 250 bp and the mm10 blacklist v2.

Quality control (QC) was performed separately for each sample using the pycisTopic qc() function. High-quality nuclei were retained on the basis of a minimum TSS enrichment score of 5, a fraction of reads in peaks (FRIP) greater than 0.30 and a minimum of 1,000 unique fragments. Sample specific cisTopic objects were generated from nuclei passing these quality control thresholds and matching the cell barcodes retained in the gene expression data QC analysis and subsequently merged into a single cisTopic object. Doublets were identified using Scrublet, assuming an expected doublet rate of 0.1 and applying a doublet-score threshold of 0.60, and predicted doublets were excluded from downstream analyses resulting in a total of 94,981 cells. The snATAC-seq nuclei yielded 710,988 accessible regions, with a mean of 7,216 accessible regions and 12,534 unique fragments per cell, a 60.56% FRIP, and a median TSS enrichment of 14.85.

For topic modelling, latent Dirichlet allocation^93^ using collapsed Gibbs sampling was performed as implemented in pycisTopic using Mallet v202108. Models containing [60, 65, 70, 75, 80, 85, 90, 95, 100, 110, 120, 130, 135, 140, 145, 150] topics were generated using 500 iterations, with an initial alpha of 50 with topic-specific alpha optimization enabled, and eta set to 0.1. Based on pycisTopic model evaluation, the model containing 140 topics was selected for downstream analysis. Next, region-topic distributions were binarized using both Otsu thresholding^94^ and the ntop method, selecting the top 3,000 regions per topic. Cell-topic distributions were binarized using Li thresholding with 100 bins. To identify DARs, normalized imputed accessibility was obtained using calculate_per_region_mean_and_dispersion_on_normalized_imputed_acc() function scaling factor of 10⁶ and normalized with 10⁴ scaling factor. Highly variable regions were selected using find_highly_variable_features() function with a minimum dispersion of 0.05, a minimum mean accessibility of 0.0125 and a maximum mean accessibility of 3. Supertype-specific and age-subclass DARs were identified using the Wilcoxon test on the imputed accessibility matrix, with an adjusted p-value threshold of 0.05 and a LFC threshold of log₂(1.5).

For CA3 analysis, cells assigned CA3 glutamatergic subtype (017 CA3 Glut) were extracted from the integrated dataset after removal of cells with low confidence or unknown annotations. Pseudobulk accessibility profiles were generated using export_pseudobulk() function in pycisTopic v2.0a0 for the CA3 annotated subtypes (leiden levels 0.3 and 0.1) and for age-subtypes (leiden level 0.1). Peaks were called on the pseudobulk profiles using MACS2 v2.2.9.1 with the following parameters: --format BEDPE, --gsize mm, --qvalue 0.05, --keep-dup all, --shift 73 and --ext_size 146. Consensus peaks were obtained using the iterative peak calling procedure implemented in pycisTopic using get_consensus_peaks() function with a peak_half_width of 250 bp and the mm10 blacklist version 2.

Quality control was performed separately for each sample. Nuclei were retained on the basis of a minimum TSS enrichment score of 5, a FRIP greater than 0.30 and a minimum of 1,000 unique fragments. Nuclei passing quality control thresholds in both snRNA-seq and snATAC- seq were retained. Doublets were identified using Scrublet v0.2.3 assuming an expected doublet rate of 0.1 and applying a doublet-score threshold of 0.60, and predicted doublets were removed from downstream analyses. The different quality control criteria resulted in a total of 7,624 cells with high-quality gene expression and snATAC-seq profiles. The 7,624 snATAC-seq nuclei yielded 417,244 accessible regions, with a mean of 11,315 accessible regions and 22,724 unique fragments per cell, a 62.50% FRIP, and a median TSS enrichment of 14.70. For topic modeling, latent dirichlet allocation using Gibbs sampling was performed, as implemented in pycisTopic using Mallet v202108 with 500 iterations across models of [30,40,45,50,60,65,70,75,80,85,90,95,100] topics. On the basis of pycisTopic model selection metrics, we selected the model with 95 topics.

Region-topic distributions were binarized using both Otsu thresholding and the ntop method, retaining the top 3,000 regions per topic, whereas cell-topic distributions were binarized using Li thresholding with 100 bins. Accessibility values were imputed using impute_accessibility() function with a scaling factor of 10⁶ and subsequently normalized using normalize_scores() function with a scaling factor of 10⁴ (Nuclei belonging to the age-leiden 0.3 resolution groups with less than 20 nuclei were filtered out, total of 44 nuclei). Highly variable regions were selected using find_highly_variable_features() function with a minimum dispersion of 0.05, a minimum mean accessibility of 0.0125 and a maximum mean accessibility of 3 resulting in 77,060 variable regions.

#### CREsted model hippocampus

We trained a sequence-to-accessibility regression model on hippocampal snATAC-seq data using CREsted^23^ v1.5.0 with age-subclass pseudobulk chromatin accessibility profiles. Cell populations represented by fewer than 100 cells were excluded prior to model training resulting in 78 groups. Accessibility counts were extracted from pseudobulk BigWig files across the hippocampal consensus peak region set (710,988 regions) using import_bigwigs() function with a target region width of 1,000 bp. Accessibility profiles were normalized using normalize_peaks() function, with normalization weights estimated from the top 3% most accessible regions. For sequence modeling, regions were resized to 2,114 bp. Consensus regions were split by chromosome into training, validation, and test sets, with chromosomes 8 and 10 used for validation, chromosomes 9 and 18 used for testing, and the remaining chromosomes used for training.

A base model was first trained on the complete hippocampal consensus peak universe using the CREsted dilated CNN architecture with an input sequence length of 2,114 bp and one output for each age-subclass accessibility profile. Training was performed with a batch size of 256, stochastic sequence shifts of up to 3 bp, and reverse-complement augmentation. The model was optimized using Adam with a learning rate of 0.001 and the CREsted CosineMSELogLoss (max_weight = 100, multiplier = 1). Training was performed for a maximum of 60 epochs, with learning-rate reduction after three epochs without improvement and early stopping after six epochs without improvement.

To improve model performance on regulatory elements displaying age-subclass specific accessibility, we subsequently fine-tuned the base model on a specificity-enriched region set. Variable regions were identified using filter_regions_on_specificity() function with a Gini standard deviation (SD) threshold of 1.0. This filtering resulted in a specificity-enriched set of 112,442 regions used for model fine-tuning. The previously trained base model was then fine-tuned on these regions using a batch size of 64, stochastic shift of 3, reverse-complement augmentation strategy, an Adam learning rate of 0.00001, and CosineMSELogLoss (max_weight=100, multiplier=1). Fine-tuning was performed for a maximum of 60 epochs, with learning-rate reduction after three epochs without improvement and early stopping after six epochs without improvement.

Predictions correlations of predictions vs ground truth and self-correlation heatmaps of ground truth for all cell types were calculated and visualized using heatmap.correlations_predictions() and heatmap.correlations_self() functions in CREsted, respectively.

#### CREsted model hippocampus downstream analysis

To interpret the cis-regulatory sequence features learned by the trained CREsted model, predictions were generated for all specificity-filtered regions using the finetuned model. For each region and age-subclass profile, experimentally observed chromatin accessibility and model-predicted accessibility were averaged to obtain a combined accessibility score. The 2,000 most specific regions per age-subclass profile were selected using sort_and_filter_regions_on_specificity() function with Gini-based specificity. Nucleotide-level contribution scores were subsequently calculated for these regions for their corresponding output classes using contribution_scores_specific() function. Next, contribution scores were analyzed using TF-MINDI^42^. High-contribution sequence segments (seqlets) were extracted using extract_seqlets() function with an importance threshold of 0.02 and three additional flanking nucleotides on either side. Seqlet contribution matrices were compared with the pycisTarget v10nr_clust_public motif collection using calculate_motif_similarity(), retaining the 80 most similar motif matches per seqlet. Seqlets were subsequently embedded and clustered using the TF-MINDI workflow with Leiden clustering at a resolution of 1.0, allowing recurrent model-derived sequence features to be grouped and annotated according to their associated motif and DBD families. Cluster-level sequence patterns were generated using the create_patterns() function with Tomtom-based motif matching, sampling up to 3,000 seqlets per Leiden cluster. The resulting patterns were used to obtain representative sequence logos and DBD-family annotations for each cluster. Seqlets were visualized in a tSNE embedding using tSNE() function and colored according to their assigned DBD family. Representative sequence patterns were additionally overlaid on the tSNE embedding using tsne_logos(). For visualization, patterns were trimmed at an information-content threshold of 0.2, and patterns shorter than four nucleotides after trimming were excluded. To compare the representation of learned sequence features across cell populations, seqlet counts were summarized by assigned DBD family. Counts were z-scored within each subclass. Cell populations and DBD families were hierarchically clustered and visualized using the dbd_heatmap() function with custom modifications.

#### GRN inference

GRNs were inferred using SCENIC+^46^ applied separately to the hippocampal dataset and CA3 cells. SCENIC+ Snakemake pipeline v1.0a2 and pycisTarget v1.1 were used. Motif enrichment was performed on DARs and binarized pycisTopic regions using *Mus musculus* custom motif annotations and custom motif ranking databases generated from the corresponding consensus peak sets. Specifically, for CA3 analysis, the motif annotation database was additionally supplemented with a custom motif derived from the TF-MINDI analysis and annotated to the TFs *St18* and *Myt1l*, as described in the deep learning section below. Both direct and orthology-based motif-TF annotations were considered. Gene annotations were obtained from the archived Ensembl BioMart release 98 (September 2019).

For motif enrichment, the motif similarity false discovery rate (FDR) threshold was set to 0.001 and the orthologous identity threshold to 0. Differential motif enrichment used a maximum of 500 background regions, a balanced 1kb promoter representation, an adjusted p-value threshold of 0.05, an LFC threshold of 1 and a motif enrichment score threshold of 3. cisTarget enrichment used AUC, NES and rank thresholds of 0.005, 3 and 0.05, respectively. pycisTarget derived cistromes consisted of TFs and their associated motif-enriched regulatory regions.

TF-to-gene and region-to-gene relationships were inferred using gradient boosting machine models, with region-to-gene inference restricted to a search space encompassing gene bodies and extending up to 150 kb upstream of gene starts and downstream of gene ends. Region-to-gene relationships were additionally evaluated using Spearman rank correlation with a correlation threshold of 0.05. eRegulons linking TFs, regulatory regions and target genes were inferred using GSEA with 3,000 permutations. Region-to-gene relationships were selected using importance quantile thresholds of 0.85, 0.90 and 0.95 and the top 5, 10 or 15 regions per gene, and eRegulons with fewer than 10 target genes were excluded. Cell-level enrichment scores for eRegulon target genes and target regions were calculated using AUCell^95^. eRegulons were further restricted to activators, with positive TF-target gene and region-target gene correlations, and repressors, with negative TF-target gene and positive region-target gene correlations. These were denoted TF(+) and TF(−), respectively. PCA followed by UMAP was performed on normalized and scaled gene-based eRegulon AUC scores. In total, GRN analysis resulted in 228 eRegulons in hippocampal data and 124 in CA3 analysis.

#### eRegulon selection

CA3 eRegulons in Figure 5 were selected using two complementary approaches. First, Spearman correlation was calculated between the gene-based AUC activity of each eRegulon and expression of its leading TF across nuclei. P-values were corrected for multiple testing using the Benjamini–Hochberg procedure, and eRegulons were retained when the adjusted p-value was <0.01 and the absolute Spearman correlation coefficient was >0.3. eRegulons were also selected if they showed differential activity between a given CA3 population and all remaining populations using a Wilcoxon rank-sum test using sc.tl.rank_genes_group() function from Scanpy v1.8.2. Within each population comparison, p-values were corrected for multiple testing across eRegulons using the Benjamini-Hochberg procedure. To additionally account for testing across the four CA3 population comparisons, the significance level was Bonferroni corrected, and eRegulons were retained when the adjusted p-value was <0.01/4 (0.0025), the absolute LFC was >0.5, and the eRegulon was active in >20% of nuclei in the corresponding population. In addition, the leading TF of each eRegulon was required to be differentially expressed in the same population using a Wilcoxon rank-sum test with an adjusted p-value <0.01/4 (0.0025), an absolute LFC >0.5, and expression in >20% of nuclei in that population. Only eRegulons for which the leading TF fulfilled these differential-expression criteria were retained by this second approach. eRegulons identified by either of the two approaches were combined. Extended eRegulons were retained only when the corresponding direct eRegulon was not available. In total, 65 eRegulons were selected.

For visualization, activating (+) and repressive (-) eRegulons were displayed separately. Gene based eRegulon AUC values were averaged within the St18-positive, St18-negative mixed, St18-negative proximal and mossy-cell populations. For each eRegulon, mean AUC values were z-score standardized across populations and truncated to the range −3 to 3. eRegulons were hierarchically clustered using Euclidean distance and complete linkage. Heatmaps were generated using clustermap() from the seaborn Python package v0.13.2. GO enrichment analysis was performed for selected CA3 eRegulons using their SCENIC+ inferred target genes with the enrichGO() function from the clusterProfiler R package v4.6.2 with mouse gene annotations from org.Mm.eg.db and was restricted to the BP ontology. P-values were corrected for multiple testing using the Benjamini-Hochberg procedure. GO terms were retained using a p-value cutoff of 0.05 and a q-value cutoff of 0.05. For visualization, the three GO terms with the smallest adjusted p-values were selected for each eRegulon. Enrichment results were visualized as dot plots using ggplot2, with dot size representing the number of eRegulon target genes associated with each GO term and color representing the adjusted p-value.

#### NMF analysis

To identify broader regulatory programs across hippocampal cell types and developmental ages, NMF^96^ was applied to activating (+) eRegulons. Gene-based eRegulon AUC scores were averaged for each subclass-by-age combination at P5, P10, P15 and P28. For each eRegulon, the resulting activity profile was z-score normalized across subclass-by-age groups. Because NMF requires non-negative input values, each standardized eRegulon profile was subsequently shifted by subtracting its minimum value. NMF was performed using the NMF implementation in scikit-learn v1.3.2 with eight components, Nonnegative Double Singular Value Decomposition with average filling (NNDSVDa) initialization, a random seed of 0 and a maximum of 1,000 iterations. The resulting factorization provided an eRegulon-by-module weight matrix and a module-by-subclass–age activity matrix. The eight modules were annotated according to their predominant activity across hippocampal cell populations. For visualization, module activities were averaged across developmental ages for each cell type and cell types were hierarchically clustered using correlation distance and average linkage.

#### CA3 gene expression analysis

CA3 gene expression data were extracted from the previously filtered and integrated snMO dataset. Nuclei carrying low-confidence (lowConfScre) or unknown annotations at the class, subclass, cluster or supertype level were excluded. The analysis was subsequently restricted to the five CA3 glutamatergic supertypes “0075 CA3 Glut-1” to “0079 CA3 Glut-5” resulting in 8,232 cells. Genes detected in fewer than three nuclei were removed. Gene expression counts were library size normalized and log-transformed using sc.pp.normalize_total() and sc.pp.log1p(), respectively, followed by scaling. Previously calculated CCA embeddings from the integrated dataset were imported and used to construct the cell-cell nearest-neighbor graph. UMAP visualization and Leiden clustering were performed on the CCA-derived neighbor graph. Leiden clustering was evaluated across resolutions ranging from 0.01 to 1.0. A resolution of 0.1 was used to define four broad CA3 populations, annotated as St18(-) proximal, St18(-) mix, St18(+) and mossy cells. For analyses requiring matched gene expression and chromatin accessibility measurements, for each sample, ATAC barcodes passing quality thresholds were intersected with high-quality CA3 nuclei identified from the gene expression data resulting in 7,624 CA3 cells. Differential gene expression was assessed in Scanpy v1.8.2 using sc.tl.rank_genes_groups() with a Wilcoxon rank-sum test. *P-*values were corrected for multiple testing using the Benjamini-Hochberg procedure.

#### CA3 Pseudotime/identity inference

To reconstruct continuous transcriptional trajectories across CA3 development and molecular identity, CA3 gene expression counts were library-size normalized and log-transformed using Scanpy v1.8.2 functions and penalized ordinal regression was performed using the ordinalNet^97^ R package v2.12. Mossy cells were excluded before modeling, and the analysis was restricted to CA3 PN populations. The 2,000 most HVGs were selected across the remaining CA3 nuclei using the variance-stabilizing transformation implemented in Seurat v5.3.0 FindVariableFeatures. The resulting log-normalized expression matrix was used as input for both developmental and spatial models.

Two independent cumulative ordinal regression models were fitted. For the temporal model, developmental age was encoded as an ordered outcome (P5<P10<P15<P28). For the molecular identity model, CA3 PNs were represented by two ordered transcriptional states, St18(+) and combined St18(−) (including St18(−) proximal and St18(−) mix populations). Both models were fitted using elastic-net regularization. The mixing parameter was evaluated from 0 to 1 in increments of 0.2, with 50 regularization parameters evaluated along the lambda path and a minimum lambda ratio of 0.0001. Hyperparameters were optimized using sample-aware cross-validation. For the developmental model, individual biological samples from ages represented by more than one sample were withheld in turn, and folds were retained only when all four developmental ages remained represented in the training set. Within each fold, predictors were standardized using the mean and SD calculated from the training data, and the same transformation was applied to the held-out nuclei. Model performance was quantified as recall for the held-out developmental age and averaged across retained folds. For the identity model, leave-one-sample-out cross-validation was performed, excluding folds in which either the training or test set lacked one of the two spatial classes. Performance was quantified as balanced accuracy, defined as the mean sensitivity across the two spatial classes. For each model, the combination of α and lambda index yielding the highest mean performance across retained folds was selected. The final models were refitted to all nuclei after gene-wise standardization.

Genes with non-zero coefficients from each ordinal regression model were ranked by absolute coefficient magnitude, and the 200 highest-weighted genes were retained separately for the developmental and identity models. For each nucleus, developmental and spatial transcriptional scores were calculated as the weighted sum of log-normalized expression values for these genes using their corresponding ordinal regression coefficients. The scores were subsequently standardized and used as continuous developmental (pseudotime) and spatial (pseudo-identity) coordinates for downstream analyses.

#### Pseudotime/identity gene expression modeling and clustering

To identify genes exhibiting coordinated changes across this two-dimensional landscape, GAMs were fitted independently for each gene using bam() from the mgcv^98^ package v1.9.1. Gene expression was modeled as the sum of smooth effects of pseudotime and pseudo- identity and a tensor product interaction between the two coordinates, using six basis functions for each dimension and a Gaussian error distribution. Fitted expression values were calculated for each nucleus. Genes for which at least one of the three smooth terms had a nominal p-value below 0.01 were retained as dynamically expressed genes, and their fitted expression profiles were z-score standardized across nuclei.

Dynamic gene expression patterns were subsequently grouped using fuzzy c-means clustering implemented with the cmeans() function from the e1071 v1.7-17 package in R v4.4.3. Fuzzy c-means clustering was performed using eight clusters, a fuzzification parameter of 2, Euclidean distance, and a maximum of 200 iterations. Genes with a membership score of at least 0.20 were assigned to the corresponding expression program, allowing individual genes to belong to more than one program. For visualization of the resulting transcriptional landscape, nuclei were divided into 32 quantile bins along each coordinate and fitted expression values were averaged within each pseudotime-by-pseudo-identity bin. GO enrichment analysis was performed separately for each fuzzy c-means^99^ gene expression program using the enrichGO() function from the clusterProfiler R package v4.14.0 with mouse gene annotations obtained from org.Mm.eg.db. Gene sets corresponding to the fuzzy expression programs were supplied as gene symbols, and enrichment was restricted to the BP ontology. A predefined background gene set consisting of all expressed genes in CA3 gene expression data was used as the enrichment universe. p-values were adjusted for multiple testing using the Benjamini–Hochberg method, and GO terms were retained with both an adjusted p-value<0.05 and q<0.05.

To determine which SCENIC+ inferred eRegulons were associated with CA3 pseudotime/identity gene programs, gene set enrichment analysis (GSEA) was performed using GSEApy^100^ v0.10.8. For each CA3 spatiotemporal gene cluster, genes were ranked in decreasing order according to their fuzzy-clustering membership score. Pre-ranked GSEA was then performed with prerank() function, using the SCENIC+ eRegulon target gene sets as gene sets. Gene sets containing fewer than 10 or more than 500 genes were excluded, and enrichment significance was estimated using 2,000 permutations with a random seed of 5. Enrichment results were summarized using NES, and eRegulon-program associations with an FDR<0.05 were retained. For visualization, activating (+) and repressive (-) eRegulons were displayed separately. Only positive NES values were visualized. Hierarchically clustered heatmaps generated with clustermap() function from seaborn package v0.13.2.

#### Pseudotime/identity accessibility modeling and clustering

Imputed chromatin accessibility profiles for CA3 variable regions obtained through pycisTopic analysis (77,060 regions) were modeled across pseudotime and pseudo-identity CA3 coordinates. For each region, accessibility was fitted using bam() from the mgcv package v1.9.1. Chromatin accessibility was modeled as the sum of smooth effects of pseudotime and pseudo-identity and a tensor product interaction between the two coordinates, using six basis functions for each dimension and a Gaussian error distribution. Fitted accessibility values were calculated for each nucleus. Regions for which at least one of the three smooth terms had a nominal p-value below 0.01 were retained as dynamically accessible regions, and their fitted accessibility profiles were z-score standardized across nuclei.

Dynamic regulatory regions were grouped according to their fitted accessibility profiles across the pseudotime/identity landscape using fuzzy c-means clustering implemented with the cmeans() function from the e1071 package in R v4.4.3. Prior to clustering, GAM-fitted accessibility profiles were z-score standardized across nuclei. Fuzzy c-means clustering was performed using eight clusters, a fuzzification parameter of 1.5, Euclidean distance, and a maximum of 200 iterations. Regions with a membership score of at least 0.20 for a given cluster were assigned to the corresponding regulatory program, allowing individual regions to belong to more than one program.

To associate dynamic chromatin accessibility programs with putative target genes, region-to-gene relationships inferred by SCENIC+ were used. To focus on positively associated regulatory relationships, links were restricted to those with correlation coefficient ρ>0.2. For each target gene, the ten region-gene links with the highest importance × rho scores were retained. For each regions cluster, putative target genes were defined as the unique genes linked to regions belonging to that cluster.

GO enrichment analysis was performed separately for the target gene sets of region clusters CR-G1 and CR-G5 using the enrichGO() function from the clusterProfiler R package v4.14.0, and mouse annotations from org.Mm.eg.db. Gene clusters containing fewer than five genes were excluded from enrichment analysis. Enrichment was restricted to the BP ontology. P-values were corrected for multiple testing using the Benjamini-Hochberg procedure, with a raw p-value cutoff of 0.05 and a q-value cutoff of 0.05.

#### Bulk RNA-seq analysis

Adapters and poly(A) tails were removed using Cutadapt^101^ v4.9 with NextSeq quality trimming (Q20), and quality control was performed using FastQC v0.12.1a and RSeQC^102^ v5.0.1. Filtered and trimmed reads were aligned to the mouse GRCm39 genome with GENCODE vM38 annotations using two-pass STAR^103^ mapping, with special multimapping parameters --winAnchorMultimapNmax --outFilterMultimapNmax 100). Gene and transposable element subfamily counts were generated using TEcount (TEtranscripts^104^ v2.2.3), in unstranded multimapping mode with RepeatMasker transposable element annotations. Transcript abundances were additionally quantified using Salmon^105^ v1.11.4 with mapping validation and GC-bias correction and summarized to genes using tximport^106^ v1.38.2. Differential expression of TEcount-derived gene and transposable element counts was analyzed in R v4.5.2 using DESeq2^107^ v1.50.2, comparing E17 with E14 samples and adjusting for sequencing run. Gene and transposable elementss with Benjamini-Hochberg-adjusted p-value < 0.05 and absolute LFC >= log2(1.5) were considered differentially expressed.

#### CREsted model CA3

To model the cis-regulatory sequence features underlying CA3 chromatin accessibility, we trained a multi-output peak-regression model using CREsted v1.5.0. Pseudobulk snATAC-seq BigWig tracks corresponding to CA3 age-by-subtype populations were used as accessibility targets. Accessibility signal was quantified using the count target over the central 1,000 bp of each regulatory region, while the corresponding DNA sequences were resized to 2,114 bp for model input. Accessibility profiles were normalized across populations using normalize_peaks() function with the top 3% of accessible regions used to estimate normalization scaling factors. Chromosomes 8 and 10 were used for validation, chromosomes 9 and 18 were used for testing, and all remaining chromosomes were used for training.

Initial model training was performed on a hippocampus-wide consensus peak universe comprising 710,988 regions. A CREsted dilated CNN was trained to predict accessibility across the CA3 age-by-subtype profiles. Training used a batch size of 256, random sequence shifts of up to 3 bp and reverse-complement augmentation. The model was optimized using Adam with a learning rate of 0.001 and CosineMSELogLoss with max_weight = 100 and multiplier = 1. Training was performed for a maximum of 60 epochs, with learning-rate reduction after three epochs without improvement and early stopping after six epochs without improvement.

The resulting model was subsequently finetuned through progressively restricted CA3 regulatory region sets. First, a CA3 consensus peak set was generated using the pycisTopic pipeline. Pseudobulk accessibility profiles were generated using export_pseudobulk() for CA3 Leiden clusters at resolutions 0.1 and 0.3, both across developmental ages and separately for each age, excluding the ‘unknown’ cluster at Leiden resolution 0.3. Peaks were called using MACS2 v2.2.9.1 through peak_calling(), and consensus peaks were obtained using the iterative peak calling procedure implemented in get_consensus_peaks(), with a peak_half_width of 250 bp and the mm10 blacklist version 2, yielding 439,492 regions. The base model was fine-tuned on these regions using a 1,000-bp target window and a 2,114-bp input sequence. Accessibility profiles were normalized across populations using normalize_peaks() function with the top 3% of accessible regions used to estimate normalization scaling factors. Finetuning was performed with a batch size of 128 and an Adam learning rate of 0.00001, while retaining the same loss function, 3-bp stochastic sequence shifts and reverse-complement augmentation. Training was continued for a maximum of 60 epochs, with learning-rate reduction and early-stopping patience of three and six epochs, respectively. The CA3 finetuned model was next trained on 74,312 variable CA3 peaks, obtained by pycisTopic find_highly_variable_features() function, using CA3 cells normalized imputed matrix. The same sequence length, accessibility quantification, chromosome split and peak-normalization procedure were applied. Finetuning was initialized from the selected CA3-consensus model and performed with a batch size of 64 and an Adam learning rate of 0.000001. The same CosineMSELogLoss, stochastic 3-bp sequence shifts, reverse- complement augmentation and stopping criteria were retained. Finally, the model was finetuned on regulatory regions with the strongest specificity across CA3 profiles. The 74,312 variable regions were filtered using crested.pp.filter_regions_on_specificity() with gini_std_threshold = 1.0. This resulted in 11,847 high specificity CA3 regions. Finetuning was initialized from the CA3-variable-region model and performed using a batch size of 64 and an Adam learning rate of 0.0000001, while retaining the same loss function, sequence augmentation and training-stopping criteria.

#### CA3 CREsted model downstream analysis

To interpret the cis-regulatory sequence features learned by the final CA3 CREsted model, nucleotide level contribution scores were calculated for the top 1,000 regulatory regions associated with each CA3 accessibility cluster (ordinal regression, pseudotime/identity inference). Contribution scores were calculated independently for all CA3 age-by-subtype output classes using integrated gradients implemented in CREsted.

Seqlets were identified using a TF-MINDI custom recursive seqlet-calling approach. Four-channel nucleotide contribution scores were first projected onto the nucleotide present in the input sequence to generate a one-dimensional signed attribution track. Attribution tracks were smoothed using a 9bp triangular kernel and normalized within each region by the 99th percentile of the absolute attribution values. Seqlet calling was then performed on the absolute normalized attribution signal using a recursive caller with a threshold of 0.05, a minimum seqlet length of 4 bp, a maximum seqlet length of 25 bp, and three additional flanking nucleotides on either side. No additional confidence or perturbation-based filtering was applied.

Because seqlet detection was performed on the absolute attribution signal, both positively and negatively contributing sequence features could be recovered. To retain this directional information, the original signed attribution values were summed across each called seqlet interval, and seqlets were classified as positively or negatively contributing according to the sign of the summed attribution. For motif analysis, seqlet contribution matrices were normalized by their maximum absolute contribution and sign-oriented according to the mean contribution across the seqlet. Similarity between model-derived seqlets and reference motifs was calculated using tfmindi.pp.calculate_motif_similarity() retaining the 100 nearest motif matches for each seqlet. Seqlets were annotated according to their originating CA3 age-by-subtype profile and clustered using Leiden clustering at a resolution of 0.5, and each cluster was assigned a DNA-binding domain (DBD) annotation based on enrichment of reference motif matches, as implemented in tfmindi.tl.cluster_seqlets(). DBD annotations for selected clusters were manually refined by examining their top-scoring reference motif matches and associated TF families.

#### KO snMO gene expression analysis and cell type annotation

The *St18-*CTRL and *St18*-KO samples were processed and quality controlled separately. Gene expression count matrices were processed using Scanpy v1.8.2. Nuclei with fewer than 200 detected genes were removed, and genes detected in fewer than three nuclei were excluded. Mitochondrial genes were identified based on the “mt-“ prefix and nuclei with ≥5% mitochondrial counts were excluded from downstream analysis. Potential doublets were assessed independently in each sample using Scrublet as implemented in Scanpy. The distribution of Scrublet doublet scores was inspected separately for the CTRL and KO samples, and a manual doublet-score threshold of 0.04 was applied to both samples. Nuclei with a doublet score ≥0.04 were removed resulting in 12492 nuclei for CTRL sample and 13079 nuclei for the KO sample.

For both samples, cell type annotation was performed independently by mapping each dataset to the Allen 10x v2/v3 reference^24^ using Seurat v5.3.0. Each query dataset was normalized using NormalizeData(), and transfer anchors between the query and reference were identified using FindTransferAnchors() with the first 30 dimensions of the reference PCA space. Reference annotations comprising class, subclass, supertype and cluster identity were transferred to each query nucleus using MapQuery(), with the reference PCA space used for mapping and the reference UMAP model used to project query nuclei onto the reference embedding. Nuclei with a label-transfer prediction score <0.2 were annotated as low-confidence (54 nuclei from CTRL sample and 71 nuclei from KO sample).

The KO and CTRL Seurat objects were merged using merge() function from Seurat. Nuclei with total_counts > 100,000 were annotated as outliers (83 nuclei, 19 of which have low confidence score annotation) and excluded from downstream analyses. Nuclei were clustered using FindClusters() at a resolution of 1. Clusters containing nuclei with ambiguous cell type annotations were assigned cluster-specific unknown labels (815 nuclei). Nuclei with low-confidence annotations (106 nuclei) or unknown cluster assignments were excluded from downstream analyses,resulting in 24588 high-quality annotated cells (12,473 KO cells and 12,115 CTRL cells). Lastly, mitochondrial genes, uncharacterized/predicted genes, and a list of sex-related genes, were removed (see above^22,87^). The merged object was normalized using NormalizeData(), highly variable features were identified using FindVariableFeatures(), and expression values were scaled using ScaleData(). PCA was performed with RunPCA() and UMAP dimensionality reduction was performed using RunUMAP() with the first 30 principal components. Marker genes for broad cell class level were identified with Scanpy v1.8.2 rank_genes_groups() function using the Wilcoxon rank-sum test with tie correction, and p-values were corrected for multiple testing using the Benjamini-Hochberg procedure (method=”wilcoxon”, tie_correct=True, corr_method=”benjamini-hochberg”). Classes represented by fewer than 50 nuclei were excluded from this analysis. Marker expression was visualized using Scanpy^327^ rank_genes_groups_dotplot() function.

Cells annotated as ‘017 CA3 Glut’ at the subclass level were extracted resulting in 1,747 nuclei (885 CTRL nuclei and 862 KO nuclei). Highly variable features were identified using FindVariableFeatures(), and expression values were scaled using ScaleData() in Seurat. PCA was performed using RunPCA(), followed by nearest-neighbor graph construction using FindNeighbors() with the first 30 principal components. Clustering was performed using FindClusters() at a resolution of 0.1, and UMAP dimensionality reduction was performed using RunUMAP() with the first 30 principal components.

Differential gene expression between St18-positive CA3 nuclei from KO (108 nuclei) and CTRL (161 nuclei) samples was assessed using the FindMarkers() function in Seurat v5.3.0 with the MAST^108^ model. Genes with an adjusted p-value <0.05 and an absolute average LFC >1 were considered significantly differentially expressed. For visualization, results were displayed as volcano plots using ggplot2 v4.0.1. The ten most significant differentially expressed genes in each direction were labeled, ranked first by adjusted p-value and subsequently by absolute LFC.

#### KO snMO snATAC-seq analysis

Quality control of *St18*-CTRL and *St18*-KO snATAC-seq samples was performed using pycisTopic. Pseudobulk accessibility profiles were generated using pycisTopic v2.0a0 the subclass and condition-by-subclass levels. Groups containing fewer than 10 nuclei were excluded from downstream analyses. For consensus peak definition, peaks were called on pseudobulk profiles using MACS2 with the following parameters: --format BEDPE, --gsize mm, --qvalue 0.01, --keep-dup all, --shift 73 and --extsize 146. Consensus peaks were obtained using the iterative peak calling procedure implemented in pycisTopic with a peak_half_width of 250 bp and the mm10 blacklist version 2.

Quality control was performed separately for each sample using get_barcodes_passing_qc_for_sample(). Nuclei were retained using a minimum TSS enrichment score of 5, a FRIP threshold of 0.20 and a minimum of 1,000 unique fragments, with automatic threshold selection disabled. Nuclei passing QC thresholds in both gene expression and chromatin accessibility analyses are retained resulting in 19667 nuclei. Doublets were identified using Scrublet, assuming an expected doublet rate of 0.1 and applying a doublet-score threshold of 0.60, and predicted doublets were excluded from downstream analyses resulting in a total of 19,531 nuclei and 362,169 regions. Cells assigned to the CA3 glutamatergic population were extracted (1598 nuclei passing gene expression and snATAC quality thresholds, and 161445 regions). For topic modeling, latent Dirichlet allocation models generated using pycisTopic were evaluated using the pycisTopic model selection metrics implemented in evaluate_models(), and the model containing 45 topics was selected for downstream analysis. UMAP dimensionality reduction was generated from the cell-topic space. Region-topic distributions were binarized using both Otsu thresholding and the ntop method, retaining the top 3,000 regions per topic, whereas cell-topic distributions were binarized using Li thresholding with 100 bins. For differential accessibility analysis, cells annotated as unknown were excluded. Accessibility values were imputed using impute_accessibility() with a scaling factor of 10⁶ and subsequently normalized using normalize_scores() with a scaling factor of 10⁴. Highly variable regions were selected using find_highly_variable_features() with a minimum dispersion of 0.05, a minimum mean accessibility of 0.0125, a maximum mean accessibility of 3 and 20 bins. DARs were identified using the pycisTopic find_diff_features() function with an adjusted p-value threshold of 0.01 and an absolute LFC threshold of 1. Population specific DARs were first identified for each CA3 annotation. Additional predefined contrasts were performed between the *St18*-positive KO, *St18*-positive CTRL, and *St18*-negative populations. DARs were visualized using seaborn clustermap function.

*St18-*predicted regions were matched to regions in the CA3 accessibility matrix using a 70% overlap threshold. For each matched region, accessibility changes were calculated as log₂(KO/CTRL) using a pseudocount of 10⁻⁶. For genes linked to multiple regulatory regions, accessibility changes were summarized as the weighted mean of the region-level LFC using importance_x_abs_rho as the weight. Gene expression changes between *St18*-positive KO and *St18*-positive CTRL CA3 cells were calculated from log-normalized expression values. For visualization, each point represented one St18-linked gene, with the x-axis indicating the weighted mean accessibility LFC of its linked regulatory regions and the y-axis indicating its gene expression LFC between KO and CTRL. Point size represented the number of regulatory regions linked to each gene. Points were colored according to the LFC from the corresponding precomputed differential expression analysis of *St18*-positive and *St18*-negative CA3 (unperturbed samples) populations using Seurat v5.3.0 FindMarkers() function with MAST test.

#### Data availability

All data will be made available upon publication.

## Acknowledgments

We thank David Hunt for contributions related to bulk RNAseq and initial identification of *St18* as a marker gene for sparsely spiny CA3 pyramidal neurons. We thank Kendra Morris, Sarah Lindo, Cassie Ramirez, Ken Sugino, Caiying Guo, Xianling Zhao, Lihua Wang, and Andrew Lemire at Janelia for their contributions with the experiments related to bulk RNA-seq. We thank Seppe De Winter and Gert Hulselmans for their help with the computational analyses. We thank Guoqiang Gu (Vanderbilt University School of Medicine, Nashville, TN USA) for generously providing the anti-ST18 antibody. We thank VIB Single Cell Core for the technical and methodological support for the knockout experiments. We thank VIB/CNL Bioimaging Core, VIB Nucleomics Core, and VIB Proteomics Leuven - Laboratory of Applied Mass Spectrometry (LAMaS) for access to instruments and technical support.

## Author contributions

Conceptualization: D. Daa., J.Vandens., D.Das., B.L.-E., S.A., and J.d.W.

Data curation: D.Daa., J.Vandens., D.Das., K.D., N.A., G.M., and K.W.

Methodology: J.Vandens., D.Daa., D.Das., B.L.-E., P.M., S.P., N.S., K.W., S.A., and J.d.W.

Investigation: D.Daa., J.Vandens., D.Das., B.L.-E., J.Vandenb., E.L., M.R., I.V., L.V., M.W., S.P., P.M., G.M., and K.W.

Validation: D.Daa., J.Vandens., D.Das., N.A., G.M., N.S., K.W., S.A., and J.d.W. Visualization: D.Daa., J.Vandens., D.Das., and K.W.

Formal analysis: D.Daa., J.Vandens., D.Das., M.R., I.V., L.V., N.A., G.M., and K.W.

Supervision: D.Das.., S.P., K.D., P.M., N.K., N.S., K.W., S.A, and J.d.W. Resources: S.P., P.M., N.S., K.W., S.A., and J.d.W.

Software: D.Daa., D.Das., N.A., N.K., and P.M.

Project administration: D.Daa., J.Vandens., D.Das., S.A., and J.d.W. Funding acquisition: N.S., S.A., and J.d.W.

Writing—original draft: D.Daa., J.Vandens., D.Das., and J.d.W.

Writing—review and editing: D.Daa., J.Vandens., D.Das., B.L.-E., N.A., P.M., N.K., G.M., N.S., K.W., S.A., and J.d.W.

