## Supplemental Information for "A subtype-specific gene regulatory program controls differential circuit integration of deep CA3 pyramidal neurons"

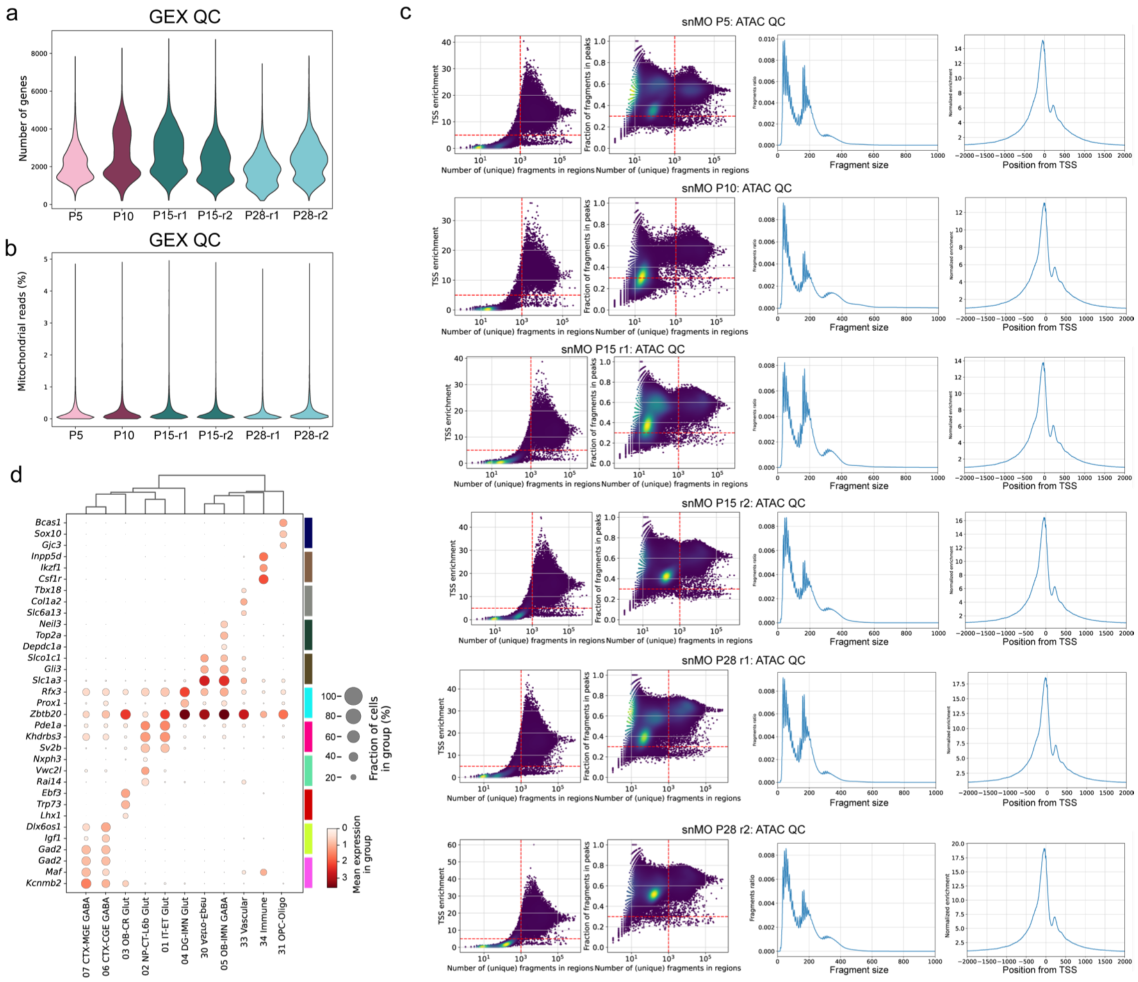


**Extended Data Fig. 1 | Quality control of postnatal hippocampal single-nucleus multiome datasets. (a, b)** Violin plots showing the number of detected genes per nucleus (a) and percentage of mitochondrial reads (b) across developmental timepoints and biological replicates. **(c)** ATAC-seq quality-control metrics for each sample. From left to right: transcription start site (TSS) enrichment versus the number of unique fragments in regions, fraction of fragments in peaks (FRiP) versus the number of unique fragments in regions, fragment-size distributions and aggregate TSS enrichment profiles. Red dashed lines indicate quality-control thresholds used for nuclei filtering. **(d)** Dot plot of the top differentially expressed genes across major cell classes (Wilcoxon rank-sum test). Dot size represents the percentage of cells expressing each gene, and color indicates scaled mean log-normalized expression.

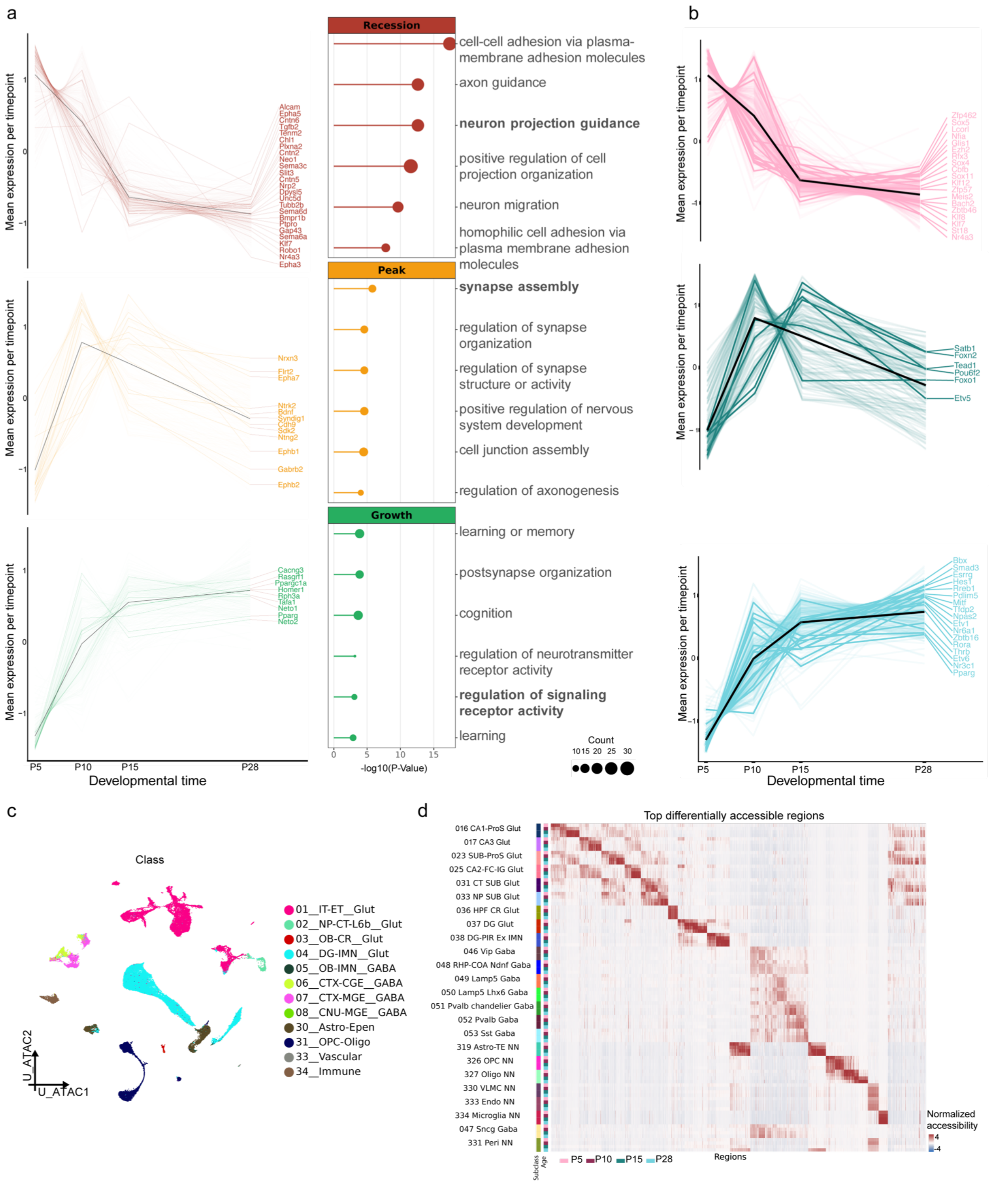


**Extended Data Fig. 2 | Gene expression and chromatin accessibility dynamics in the postnatal mouse hippocampus. (a)** Temporal gene expression trajectories and enriched Gene Ontology (GO) biological process terms for recession, peak and growth expression patterns in CA3 PNs. The trough pattern was excluded from the analysis due to low number of genes. Labeled genes correspond to those associated with the highlighted GO terms. Black lines indicate mean expression trajectories. **(b)** Temporal expression trajectories of transcription factors assigned to recession, peak and growth patterns in CA3. **(c)** UMAP of snMO ATAC profiles colored by major cell class. **(d)** Heatmap of the top differentially accessible regions (DARs) across hippocampal subclass-by-age groups. Colors indicate normalized chromatin accessibility.

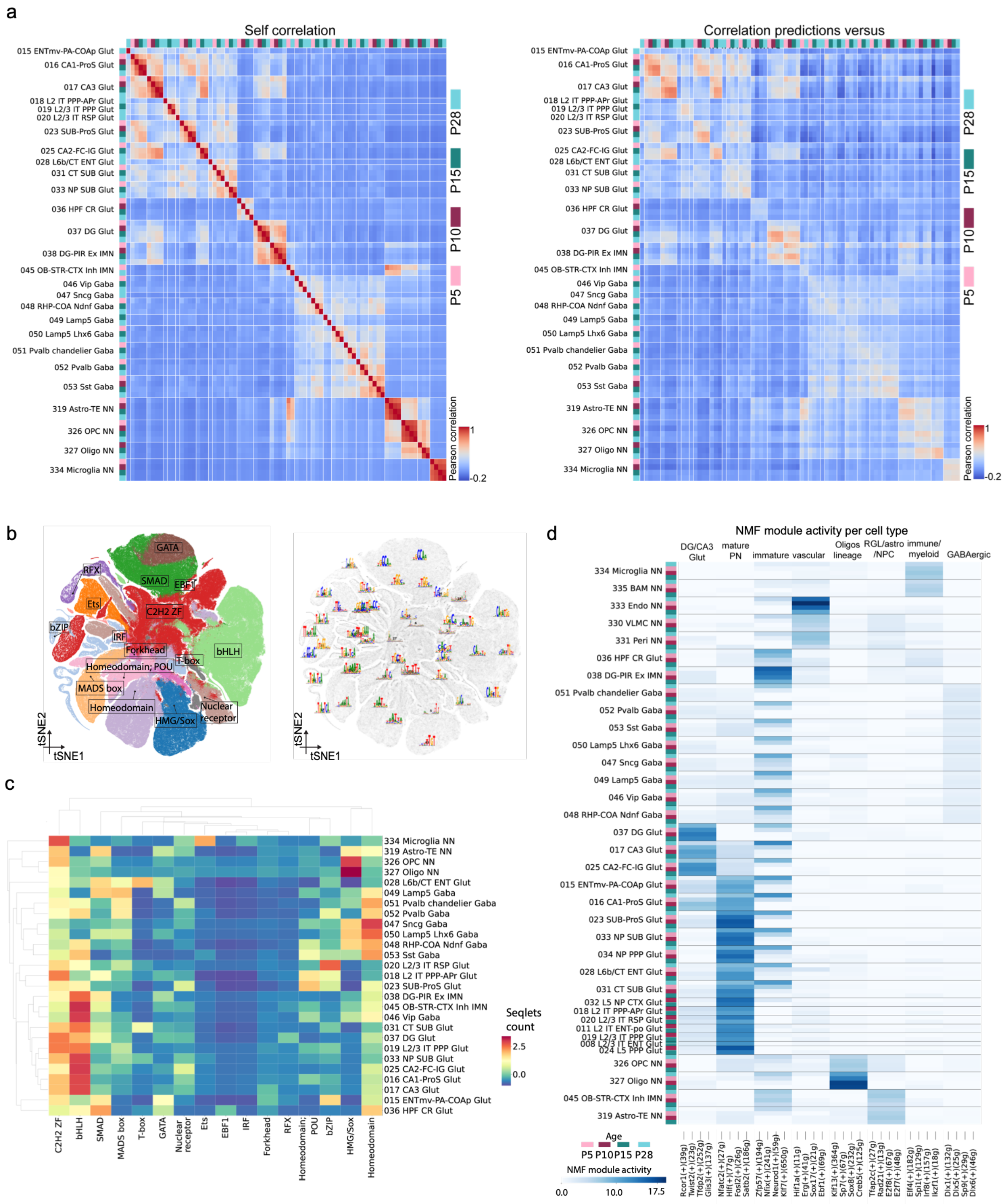


**Extended Data Fig. 3 | Cell-type-specific cis-regulatory features and gene regulatory programs across the postnatal hippocampus. (a)** Heatmaps of pairwise Pearson correlations between observed chromatin accessibility profiles (left) and between CREsted DeepPeak predictions and observed accessibility profiles (right) across hippocampal subclass-by-age groups. **(b)** t-SNE visualization of TF-MINDI-derived seqlets extracted from CREsted DeepPeak contribution scores and clustered based on sequence similarity. Seqlets are colored by their inferred DNA-binding domain (DBD) family (left), with representative consensus motif logos shown for the identified Leiden clusters (right). **(c)** Heatmap showing the number of seqlets assigned to each DBD family across hippocampal subclasses. **(d)** Heatmap showing non-negative matrix factorization (NMF) module activity across hippocampal subclasses and developmental timepoints. Columns represent NMF modules annotated with their top four contributing eRegulons, and rows represent subclass-by-age groups.

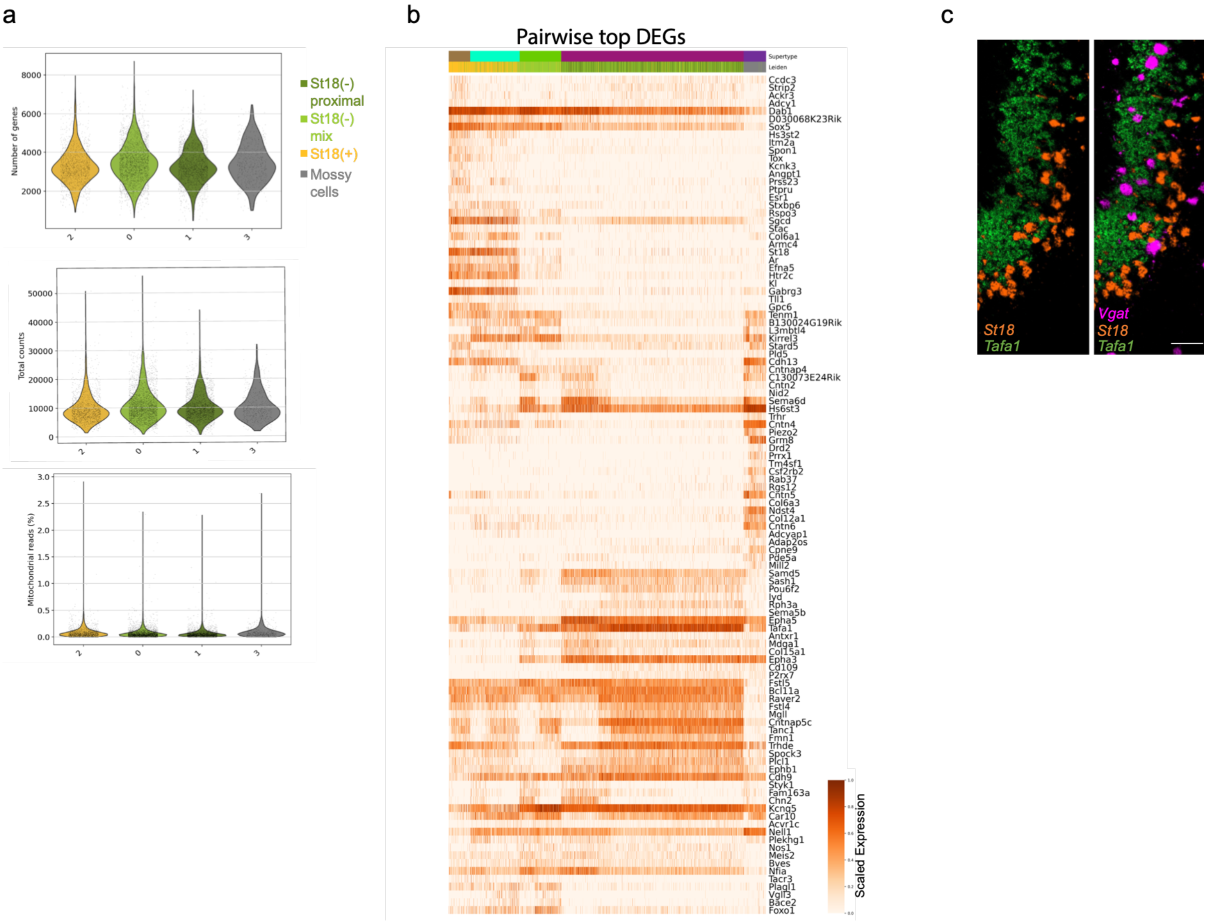


**Extended Data Fig. 4 | Quality control and molecular characterization of CA3 pyramidal neurons. (a)** Violin plots showing the number of detected genes, total RNA counts and percentage of mitochondrial reads across CA3 clusters. **(b)** Heatmap of top differentially expressed genes identified by pairwise comparisons between CA3 supertypes (Wilcoxon rank-sum test). **(c)** smFISH RNAscope images showing expression of *St18* (orange) in deep-layer CA3, *Tafa1* (green) in superficial-layer CA3 and *Vgat* (magenta) in CA3. Scale bar, 50 μm.

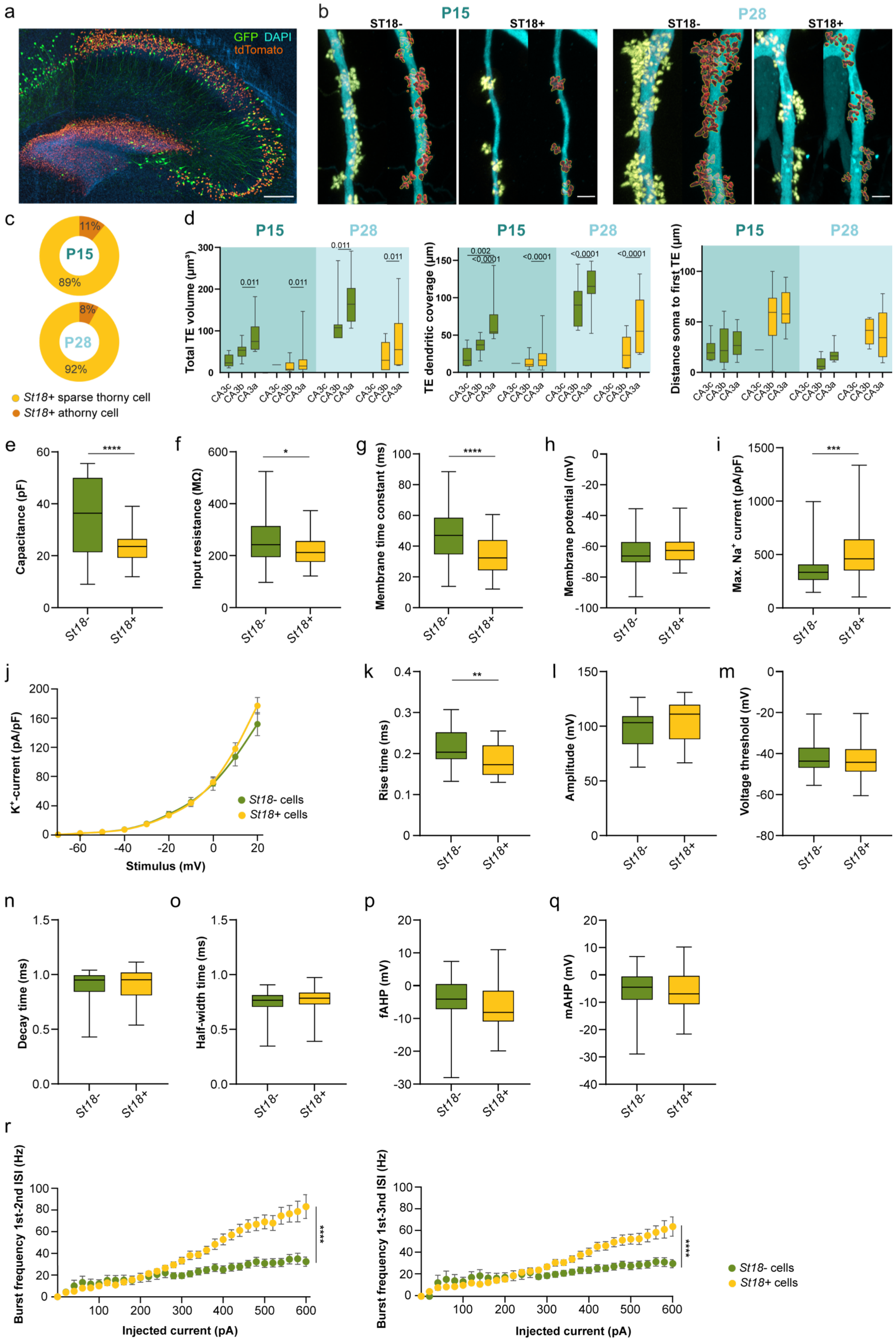

a
GFP DAPI
tdTomato
b
P15
ST18-
ST18+
P28
c
P15
11%
89%
P28
8%
92%
St18+ sparse thorny cell
St18+ athorny cell
d
Total TE volume (µm³)
TE dendritic coverage (µm)
Distance soma to first TE (µm)
CA3c
CA3b
CA3a
e
Capacitance (pF)
f
Input resistance (MΩ)
g
Membrane time constant (ms)
h
Membrane potential (mV)
i
Max. Na⁺ current (pA/pF)
St18-
St18+
j
K⁺-current (pA/pF)
Stimulus (mV)
St18- cells
St18+ cells
k
Rise time (ms)
l
Amplitude (mV)
m
Voltage threshold (mV)
n
Decay time (ms)
o
Half-width time (ms)
p
fAHP (mV)
q
mAHP (mV)
r
Burst frequency 1st-2nd ISI (Hz)
Burst frequency 1st-3nd ISI (Hz)
Injected current (pA)

**Extended Data Fig. 5 | Morphological and intrinsic properties of *St18*-positive and *St18*-negative neurons. (a)** Representative 200 µm-thick cleared sagittal *St18*-Cre hippocampal section showing sparse GFP labeling and Cre-dependent nuclear tdTomato expression. **(b)** Representative dendritic segments from *St18*-negative and *St18*-positive CA3 PNs at P15 (left) and P28 (right) from Fig. 3f, showing segmented TEs (yellow) and corresponding 3D reconstructions (red). **(c)** Proportions of sparsely thorny and athorny neurons among analyzed *St18*-positive CA3 PNs at P15 and P28. **(d)** Quantification of TE morphological parameters along the proximodistal axis of CA3 at P15 and P28: total TE volume (two-way ANOVA Tukey's multiple comparisons test, p-values indicated); TE dendritic coverage (two-way ANOVA Sidak multiple comparisons test, p-values indicated); and the distance from soma to first TE (two-way ANOVA Tukey's multiple comparisons, no significance). **(e–h)** Measured passive membrane properties, including membrane capacitance (e; $p<0.0001$; n=43 *St18*-negative and 52 *St18*-positive cells), input resistance (f; $p=0.028$; n=57 *St18*-negative and 61 *St18*-positive cells), membrane time constant (g; $p<0.0001$; n=57 *St18*-negative and 61 *St18*-positive cells), and resting membrane potential (h; $p=0.181$; n=61 *St18*-negative and 64 *St18*-positive cells) (all Mann–Whitney tests). **(i)** Measured maximum $Na^+$ current (Mann-Whitney test, $p=0.0001$; n=43 *St18*-negative and 52 *St18*-positive cells). **(j)** $K^+$ currents normalized to cell capacitance (two-way repeated-measures ANOVA, $p=0.537$; n=43 *St18*-negative and 52 *St18*-positive cells). **(k–q)** Action potential properties, including rise time (k; $p=0.002$), amplitude (l; $p=0.062$), voltage threshold (m; $p=0.535$), decay time (n; $p=0.613$), half-width (o; $p=0.428$), fast afterhyperpolarization (p; $p=0.198$) and medium afterhyperpolarization (q; $p=0.865$) (all Mann–Whitney tests; n=30 *St18*-negative and 38 *St18*-positive cells). **(r)** Burst frequency calculated from the first-second (left; $p<0.0001$) and first-third (right; $p<0.0001$) interspike intervals (ISIs) (all mixed-effects models; n=55 *St18*-negative and 62 *St18*-positive cells). Scale bars, 200 µm (a), 3 µm (b).

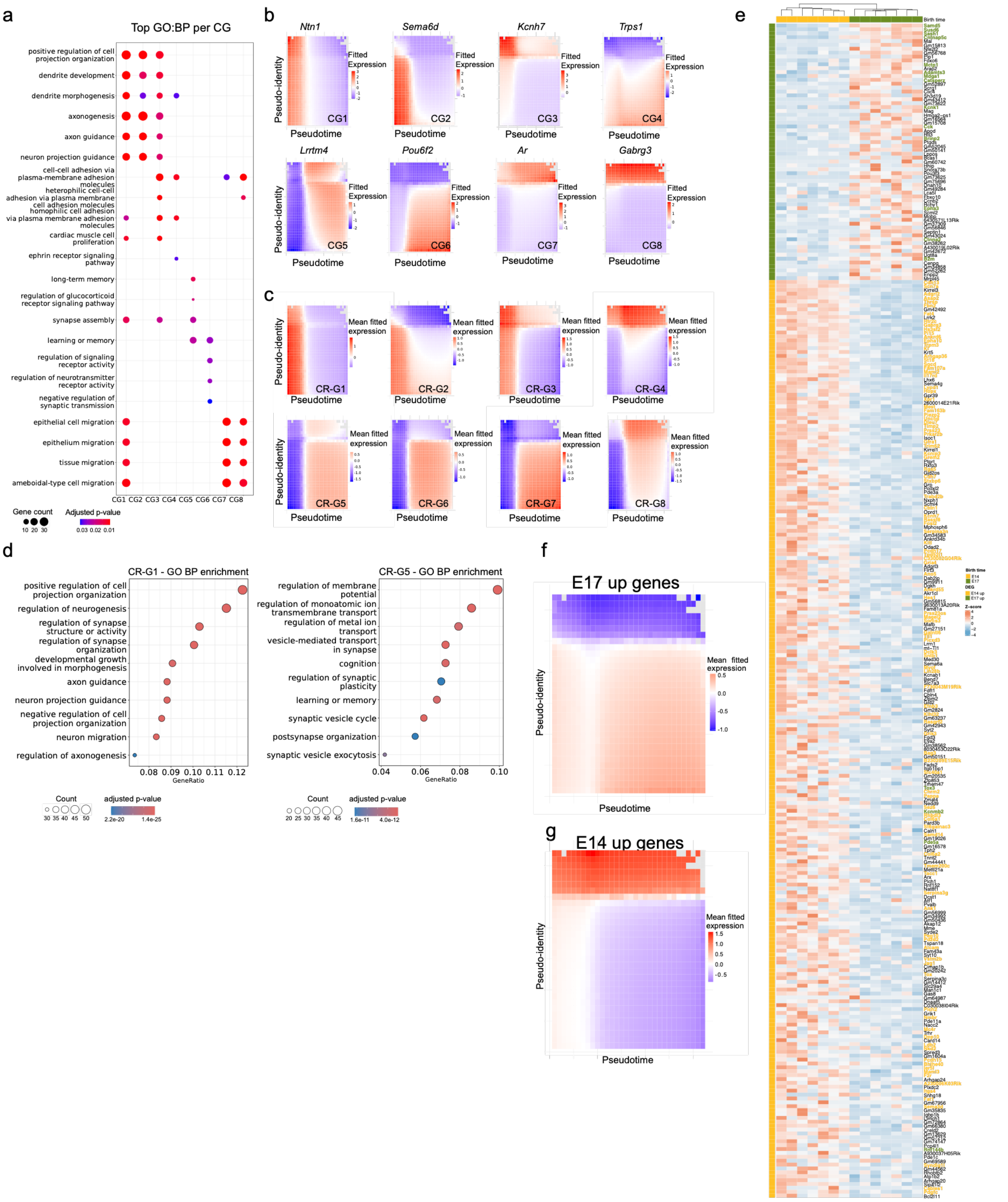


**Extended Data Fig. 6 | CA3 subtype-specific transcriptional and chromatin accessibility dynamics across postnatal development. (a)** Gene Ontology (GO) biological process enrichment across the eight dynamic gene expression clusters (CG1-CG8). Dot size indicates gene count, and color indicates adjusted p-value. **(b)** Generalized additive model (GAM)-fitted expression profiles of representative genes from each dynamic expression cluster across pseudotime and pseudo-identity. **(c)** Mean GAM-fitted expression profiles across pseudotime and pseudo-identity for genes linked to dynamically accessible regions in each chromatin regulatory cluster (CR1-CR8). **(d)** GO biological process enrichment of genes linked to regulatory regions in CR1 and CR5. **(e)** Heatmap of genes differentially expressed between E14- and E17-born CA3 neurons from adult mice. Columns represent individual samples, and colors indicate scaled variance-stabilized expression. Genes that are differentially expressed in *St18*-positive CA3 are colored in yellow and genes differentially expressed in *St18*-negative CA3 are colored in green. **(f, g)** Mean GAM-fitted expression profiles across pseudotime and pseudo-identity for genes upregulated in E17-born (f) and E14-born (g) CA3 neurons, respectively.

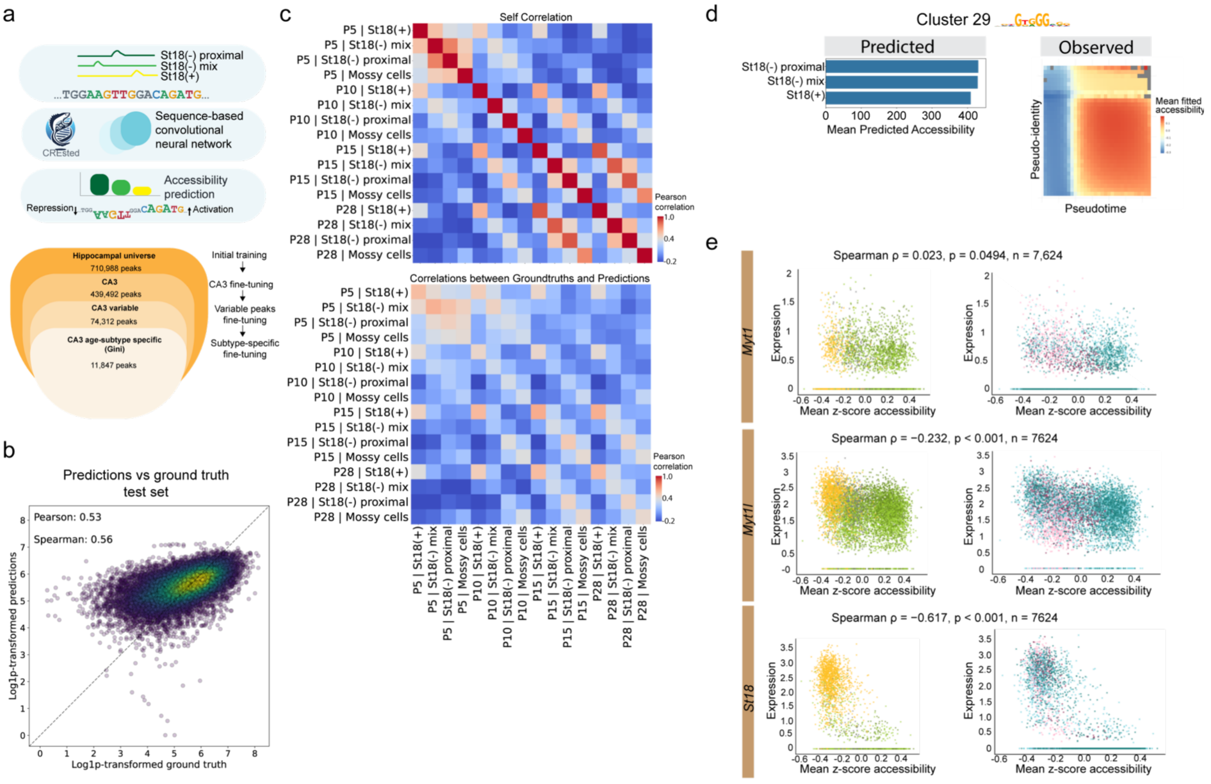


**Extended Data Fig. 7 | CREsted model performance and regulatory sequence features underlying CA3 subtype-specific chromatin accessibility. (a)** Schematic overview of the CREsted DeepPeak model and sequential training strategy, from hippocampal accessible regions to CA3 subtype-specific regulatory regions. **(b)** Scatterplot comparing CREsted-predicted and observed chromatin accessibility across held-out test regions. Pearson and Spearman correlation coefficients are indicated. **(c)** Heatmaps of pairwise Pearson correlations between observed chromatin accessibility profiles (top) and between CREsted DeepPeak predictions and observed accessibility profiles (bottom) across CA3 subtype-by-age groups. Model predictions were log1p-transformed before correlation analysis. **(d)** Mean CREsted-predicted accessibility across CA3 subtypes (left) and mean GAM-fitted observed accessibility across pseudotime and pseudo-identity (right) for regions containing seqlets assigned to TF-MINDI Leiden cluster 29. **(e)** Scatterplots showing the relationship between *Myt1*, *Myt1l* and *St18* expression and mean z-scored accessibility of regions containing AAGTT/MYT-like motif-associated seqlets (Leiden cluster 30). Nuclei are colored by CA3 subtype (left) or developmental age (right). Spearman correlation coefficients, p-values and number of nuclei are indicated.

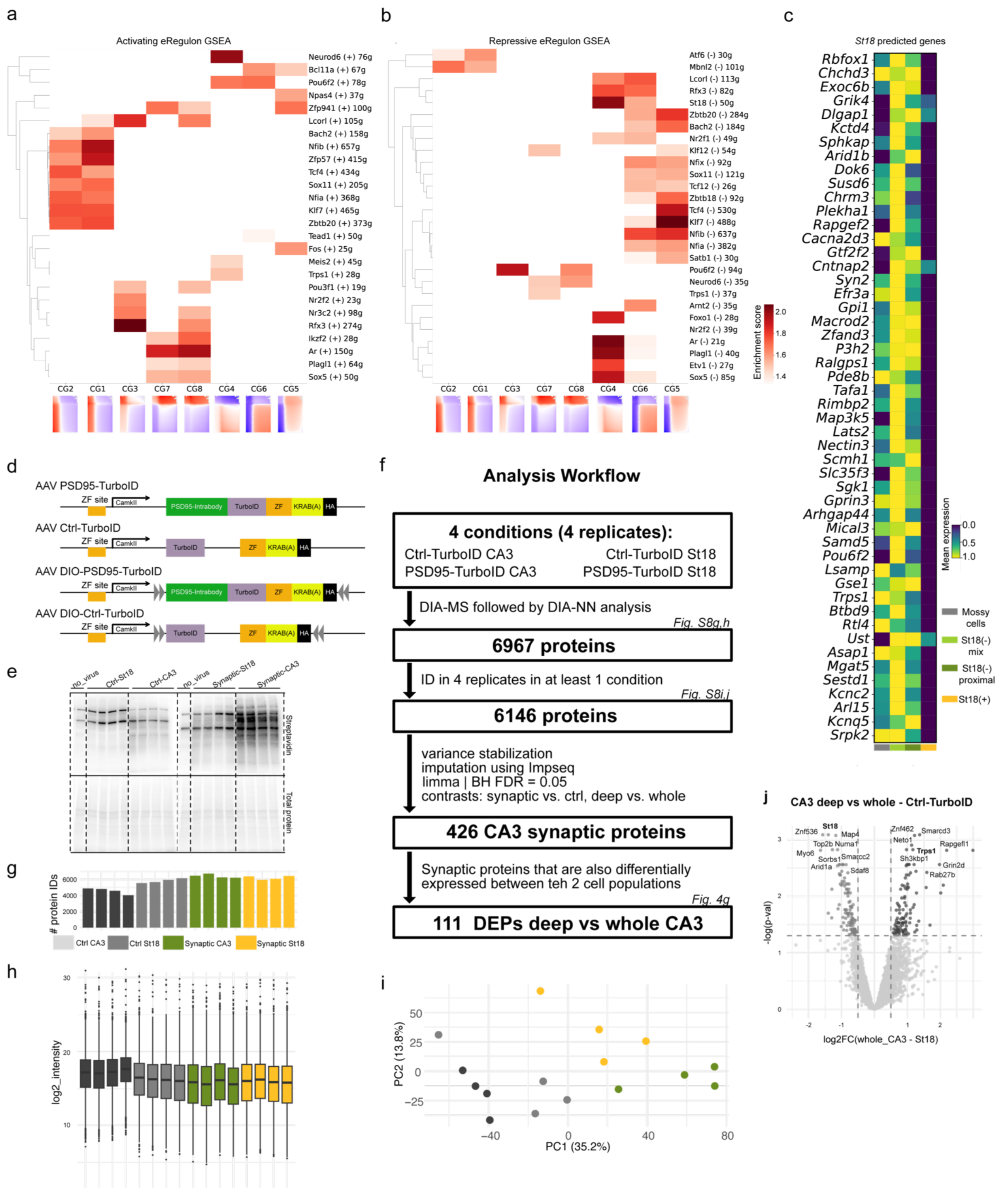


**Extended Data Fig. 8 | Gene regulatory networks and proteomic profiling of CA3 pyramidal neuron subtypes. (a, b)** Gene set enrichment analysis (GSEA) of activating (+) (a) and repressive (−) (b) SCENIC+ eRegulons across eight dynamic CA3 gene expression clusters (CG1-CG8). Genes were ranked by their fuzzy-clustering membership scores for each cluster. Rows represent eRegulons and columns represent gene expression clusters. Colors indicate normalized enrichment scores (NES); only positive enrichments with FDR < 0.05 are shown. The number of predicted target genes is indicated for each eRegulon. Heatmaps below show the mean fitted expression profiles of the corresponding gene clusters across pseudotime and pseudo-identity. **(c)** Heatmap showing mean expression of predicted *St18* target genes across CA3 subtypes. **(d)** Schematic of the AAV constructs used in this study. Intrabody-TurboID constructs consist of target-specific intrabodies fused to TurboID, as well as a zinc-finger-KRAB repressor, using flexible linkers. The repressor binds a sequence upstream of the promoter, thereby inhibiting its own transcription in an unbound state. Ctrl-TurboID construct is identical, but without any intrabody. DIO-constructs are identical but with inverted LoxP sequences. All constructs are downstream of a short neuron-specific CamkII promoter. **(e)** Western blot showing biotinylation levels (streptavidin-HRP), and total protein levels for replicates used in the experiment. **(f)** Data analysis workflow. **(g)** Number of protein IDs in each sample. **(h)** Distribution of MS intensities and in each sample. Box-plots show median, minima, maxima and interquartile range. **(i)** PCA showing separation of different replicates based on condition. **(j)** Volcano plot showing DEPs in Ctrl-TurboID conditions between deep and whole CA3 PNs (limma moderated test BH FDR=0.05, log2FC threshold is 0.5 between conditions).

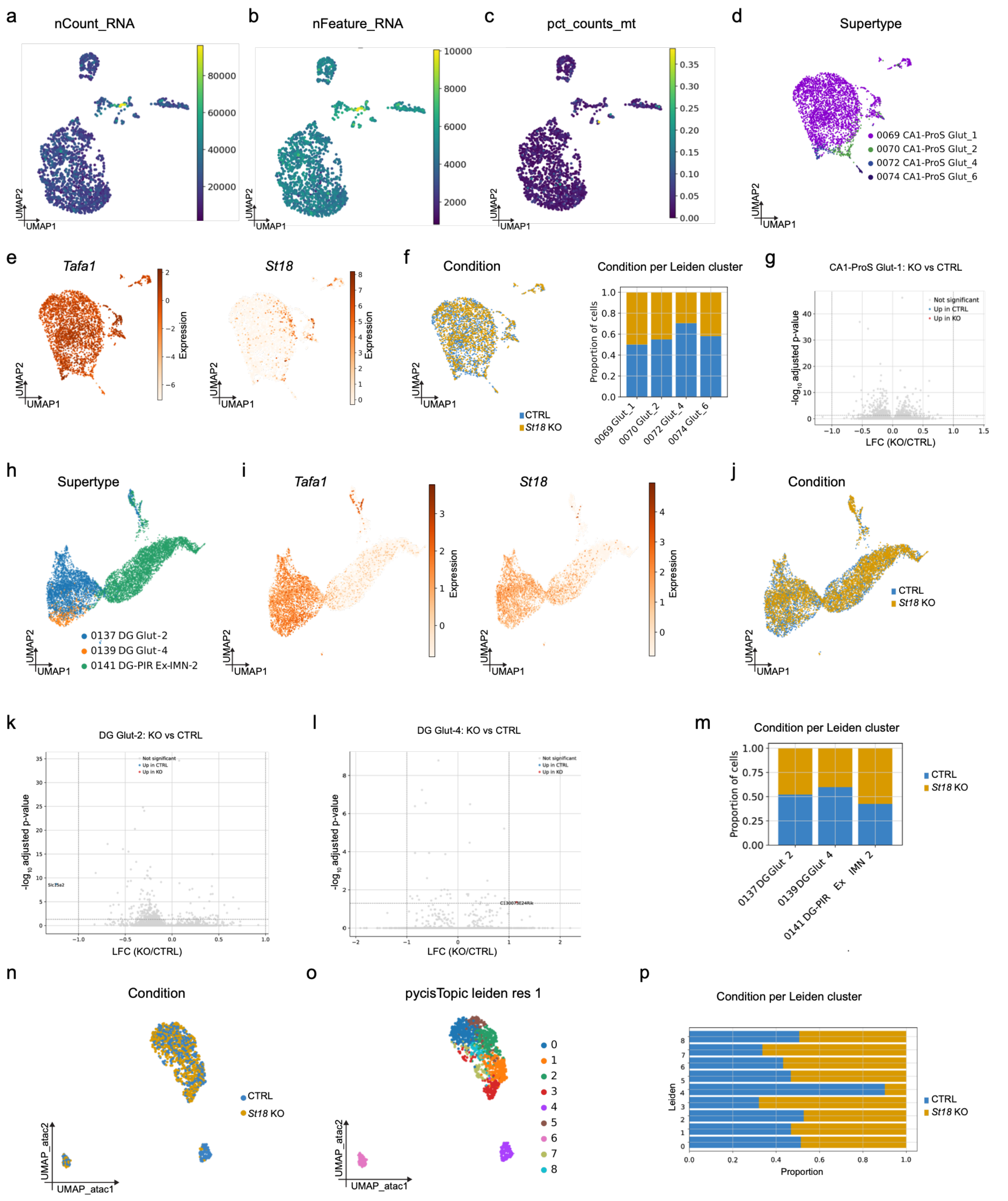


**Extended Data Fig. 9 | Transcriptional and chromatin accessibility characterization of neuronal populations following *St18* deletion. (a-c)** UMAP representations of CA3 PNs colored by the number of RNA counts (nCount_RNA; a), number of detected genes (nFeature_RNA; b) and percentage of mitochondrial transcripts (pct_counts_mt; c). **(d)** UMAP showing the distribution of CA1-ProS neuronal supertypes. **(e)** UMAPs showing the expression of *Tafa1* and *St18* across CA1-ProS neurons. (**f**) UMAP showing the distribution of control (CTRL) and *St18* knockout (KO) cells (left), and stacked bar plot showing the proportion of cells from each condition across CA1-ProS supertypes (right). (**g**) Volcano plot showing differential gene expression between *St18* KO and CTRL neurons in the CA1-ProS Glut-1 population. **(h)** UMAP showing the distribution of dentate gyrus (DG) neuronal supertypes. **(i)** UMAPs showing the expression of *Tafa1* and *St18* across DG neurons. **(j)** UMAP showing the distribution of CTRL and *St18* KO cells across DG neuronal populations. **(k,l)** Volcano plots showing differential gene expression between *St18* KO and CTRL neurons in DG Glut-2 (k) and DG Glut-4 (l) populations. **(m)** Stacked bar plot showing the proportion of CTRL and *St18* KO cells across DG neuronal supertypes. **(n)** UMAP representation of single-nucleus ATAC-seq profiles of CA3 pyramidal neurons (PNs), colored by condition (CTRL or *St18* KO). **(o)** UMAP showing Leiden clustering of CA3 PNs based on chromatin accessibility profiles. **(p)** Stacked bar plot showing the proportion of CTRL and *St18* KO nuclei across CA3 Leiden clusters.

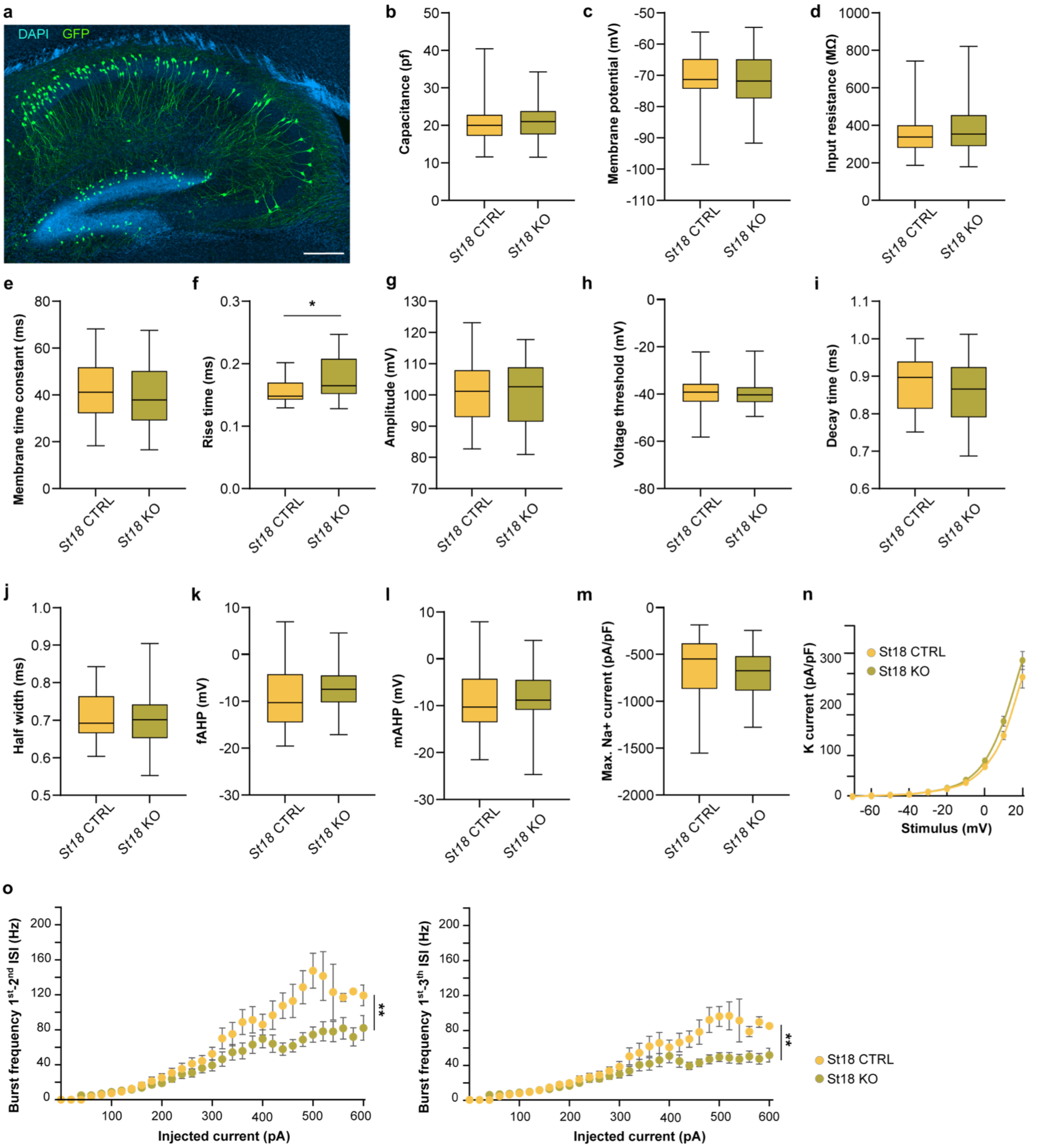


**Extended Data Fig. 10 | Intrinsic electrophysiological properties of *St18*-CTRL and *St18*-KO neurons. (a)** Representative 200 µm thick cleared sagittal hippocampal section showing sparse GFP labeling of PNs. Scale bar 200 µm. **(b-e)** Passive membrane properties including: (b) capacitance (p=0.394; n= 38 *St18*-CTRL and 46 *St18*-KO cells), (c) resting membrane potential (p=0.950; n= 30 *St18*-CTRL and 42 *St18*-KO cells), (d) input resistance (p=0.497; n= 30 *St18*-CTRL and 39 *St18*-KO cells), and (e) membrane time constant (p=0.489; n= 30 *St18*-CTRL and 39 *St18*-KO cells) (all Mann-Whitney tests). **(f-l)** Action potential properties, including (f) rise time (p=0.013), (g) amplitude (p=0.908), (h) voltage threshold (p=0.539), (i) decay time (p=0.177), (j) half width (p=0.378), (k) fast afterhyperpolarization (p=0.063), (l) medium afterhyperpolarization (p=0.214) (all Mann-Whitney tests; n= 23 *St18*-CTRL and 30 *St18*-KO cells). **(m)** Measured maximum $Na^+$ current (Mann-Whitney test, p=0.147; n=

43 *St18*-CTRL and 52 *St18*-KO cells). **(n)** $K^+$ current traces normalized to cell capacitance (two-way repeated-measures ANOVA, p=0.091; n= 38 *St18*-CTRL and 46 *St18*-KO cells). **(o)** Burst frequency calculated from the first–second (left; p=0.002) and first–third (right; p=0.005) interspike intervals (ISIs) (all mixed-effects models, n= 39 *St18*-CTRL and 30 *St18*-KO cells).
